# Two distinct *Trypanosoma* eIF4F complexes co-exist, bind different mRNAs and are regulated during nutritional stress

**DOI:** 10.1101/2024.04.15.589194

**Authors:** Bernardo Papini Gabiatti, Eden Ribeiro Freire, Jimena Ferreira da Costa, Mariana Galvão Ferrarini, Tatiana Reichert Assunção de Matos, Henrique Preti, Isadora Munhoz da Rocha, Beatriz Gomes Guimarães, Susanne Kramer, Nilson Ivo Tonin Zanchin, Fabíola Barbieri Holetz

**Author notes:** The authors share equal contribution to this work (Bernardo Papini Gabiatti, Eden Ribeiro Freire). To whom correspondence should be addressed: Fabíola Barbieri Holetz.

## Abstract

Many eIF4F subunits and PABP paralogues are found in trypanosomes: six eIF4E, five eIF4G, one eIF4A and two PABPs. They are expressed simultaneously and assemble into different complexes, contrasting the situation in metazoans that use distinct complexes in different cell types or developmental stages. Each eIF4F complex has its own proteins, mRNAs and, consequently, a distinct function. We set out to study the function and regulation of the two major eIF4F complexes of *Trypanosoma cruzi* and identified the associated proteins and mRNAs of eIF4E3 and eIF4E4 in cells in exponential growth and in nutritional stress. Upon stress, eIF4G/eIF4A and PABP remain associated to the eIF4E, but the associations with other 43S pre-initiation factors decrease, indicating that ribosome attachment is impaired. Most eIF4E3-associated mRNAs encode for metabolic proteins, while eIF4E4 associate to mRNAs encoding ribosomal proteins. Interestingly, for both eIF4E3/4, more mRNAs were associated in stressed cells than in non-stressed cells, even though these mRNAs have lower translational efficiencies in stress. In summary, trypanosomes have two co-existing eIF4F complexes involved in translation of distinct mRNA cohorts important for growth. Under stress conditions, both complexes exit translation but remain bound to their mRNA targets.

## 3. Introduction

Translational control is an important post-transcriptional mechanism of gene expression regulation [1]. Translation initiation depends on eukaryotic translation initiation factors (eIFs) to assemble the elongation-competent 80S ribosome with a tRNA initiator positioned at the start codon of the mRNA [2]. Two main mechanisms regulate translation initiation. One targets the start codon recognition by the ternary complex (TC). This regulation is mediated by stress-activated kinases which phosphorylate the eIF2 subunit of the TC and decrease its availability, resulting in bulk translational arrest [3]. The other targets the ribosome attachment to the mRNA [4] and involves the 43S pre-initiation complex (43S PIC), formed by eIF1, eIF1A, eIF3, the small ribosomal subunit and the TC. The mRNA and the 43S PIC are connected by the eIF4F complex.

The eIF4F complex is formed by the cap-binding subunit eIF4E, the scaffolding subunit eIF4G and the RNA helicase eIF4A. The mRNA is recruited to translation by association of eIF4E to the 5’ end cap and the poly-(A) binding protein (PABP) to the 3’ end poly(A) tail. eIF4G is a large protein that binds to eIF4E, eIF4A, mRNA, PABP and eIF3. One classical example of translational control during stress is the competition between 4E-BP (eIF4E-binding protein) and eIF4G for eIF4E, which results in a decrease in translation, especially for TOP mRNAs (mRNAs with a 5’ terminal <u>o</u>ligo<u>p</u>yrimidine tract) [5]. The interaction between eIF4G and the eIF3 complex connects the eIF4F complex to the 43S PIC. Because eIF4G binds to both the cap-binding eIF4E and the poly(A) binding PABP, it facilitates a ‘closed loop’ conformation of the mRNA [6]. This is believed to speed up recycling of ribosomes within the polysomes, increasing translational efficiency.

Many other translational control mechanisms evolved within eukaryotic taxa to regulate translation initiation [7]. One is the duplication of genes encoding the eIF4F complex subunits [8] and PABP [9]. Duplicated genes evolved to encode different paralogues that can assemble into distinct eIF4F complexes, each specialised to a specific environmental or developmental condition [10]. Typically, one ‘canonical’ eIF4F complex regulates the translation of bulk mRNAs while others fulfill special functions, for example during hypoxia in metazoans [11] or during development in nematodes [12]. The distinct eIF4F isoforms bind to distinct sets of mRNAs, mediated by (i) different affinities of eIF4E isoforms to the cap, 4E-BP and/or eIF4G [13] (ii) different affinities of eIF4F isoforms to other RNA binding proteins (RBPs) and (iii) localisation of eIF4F isoforms to distinct ribonucleoprotein granules [14]. The largest functional diversity of eIF4F subunit paralogues is found in protists [15] [16].

Kinetoplastida include the human and animal pathogenic trypanosomatids *Trypanosoma* and *Leishmania*. They are digenetic and shuttle between an invertebrate and vertebrate host in replicative and non-replicative stages. Trypanosomes are great models to study translational control because of their reliance in post-transcriptional mechanisms for gene expression regulation across a complex life cycle [17], [18]. The reason is the absence of gene-specific promoters, due to the highly unusual way of mRNA transcription and processing: kinetoplastid mRNAs are transcribed as long polycistrons (>100 kb) and processed by *trans*-splicing at the 5’ end coupled to 3’ polyadenylation of the upstream transcript [19], [20]. In *trans*-splicing, a short 39-nucleotide spliced leader (SL) sequence previously modified at the 5’-end with a unique cap structure named cap-4 is added to the 5’ end of every mRNA [21]. Trypanosomatid genomes encode multiple paralogues of the eIF4F subunits: six eIF4E (eIF4E1-6), five eIF4G (eIF4G1-5) and one eIF4A (eIF4A1). Moreover, *Trypanosoma* genomes encode two PABP isoforms (PABP1-2); *Leishmania* and other trypanosomatids encodes a third one (PABP3). The functional characterization of eIF4F-like complexes in trypanosomatids has mostly focused on *T. brucei* and *Leishmania* [22], while less is known about *T. cruzi*.

Of the six trypanosomatid eIF4E paralogues, eIF4E3 and eIF4E4 appear to play the dominant role in translation [23] because (i) they are the most abundant [23], (ii) they are essential for survival [24], [25], (iii) they activate the expression of a reporter mRNA in tethering assays [26] and (iv) they associate to polysomes [27]. Moreover, both can form ‘eIF4F-like’ complexes with eIF4G4 and eIF4G3, respectively (but only eIF4G3 binds to eIF4A1) [28]. Only recently, *T. brucei* eIF4E3 and eIF4E4 were reported to associate to different sets of mRNAs in procyclic cells (the replicative stage of the invertebrate host) [29]. eIF4E3 seems to be the ‘canonical’ eIF4E and associates to mRNAs coding for the most abundant protein in this stage, procyclin, RBPs and metabolic proteins. In contrast, eIF4E4 is specialised to mRNAs encoding ribosomal proteins [29]. How the selectivity of the cap-binding proteins to certain mRNAs is achieved, when every trypanosomatid mRNA has the same capped SL at their 5’ end, is still unknown.

The Kinetoplastida eIF4E3 and eIF4E4 paralogues have unique discorded amino-terminal extensions with potential to interact with PABP. This extension contains two to three PABP-association motifs (PAM2) [30]. PAM2 motifs interact with the MLLE domain of the PABP C-terminus [31] and are present in several other mRNA metabolism proteins [32]. The presence of PAM2 motifs in an eIF4E was so far only reported in the trypanosomatid eIF4E3 and eIF4E4. While the eIF4E4:PABP1 interaction has been demonstrated experimentally in *Leishmania* [31], the eIF4E3:PABP interaction has not yet. As eIF4E3 and eIF4E4 have both cap– and PABP-binding sites, the eIF4G:PABP interaction conserved in all other eukaryotes could be redundant in kinetoplastids. Regarding the interaction between eIF4G3 or eIF4G4 and PABP, results have not been clear or reproductible [28], [31], [33], [34]. The amino acids that mediate the PAM2:MLLE interaction [35] are conserved in Kinetoplastida eIF4E3/eIF4E4:PABP1/PABP2, however, there is specific *in vivo* preference for interactions: when used as bait, PABP1 specifically co-precipitates with eIF4E4 [36], [37] and this is confirmed in the reverse assay; eIF4E4 co-precipitates PABP1 [24], [34]. PABP2 is more promiscuous and co-precipitates most translation initiation factors, including eIF4E3 [36], but there is no experimental data supporting a direct eIF4E3:PABP2 interaction so far. How the interaction preferences between the two PABP and the eIF4E orthologs are mediated when the interaction motifs are the same is not known.

The life cycle of trypanosomatids is coupled to extreme environmental changes. Usually, the replicative form of the invertebrate host encounters different types of stress and this triggers a response to differentiation into the non-replicative infective form. *T. cruzi* replicative epimastigotes proliferate in the hindgut of the triatomine bug. As epimastigotes migrate into the intestinal tract of the insect, nutrients become scarce, triggering differentiation into the non-replicative, infective metacyclic trypomastigotes. This differentiation is called metacyclogenesis and can be induced *in vitro*: epimastigotes, usually cultured in carbon-rich medium (liver infusion tryptose, LIT), are concentrated and transferred to an acidic medium lacking carbon sources (triatomine artificial urine medium, TAU) and subsequently to TAU supplemented with three amino acids (TAU3AAG, proline, glutamate, aspartate) for four days [38] (Figure 1A). Experiments that aim to investigate molecular mechanisms involved in *T. cruzi* differentiation focus mainly on two stages of this process: (i) starvation in TAU medium for two hours, which induces global translation arrest, polysome disassembly, eIF2α phosphorylation [39], [40] and formation of stress granules [41], [42], [43], and (ii) recovery in TAU3AAG, which causes a partial return in translation [40].

**Figure 1.**
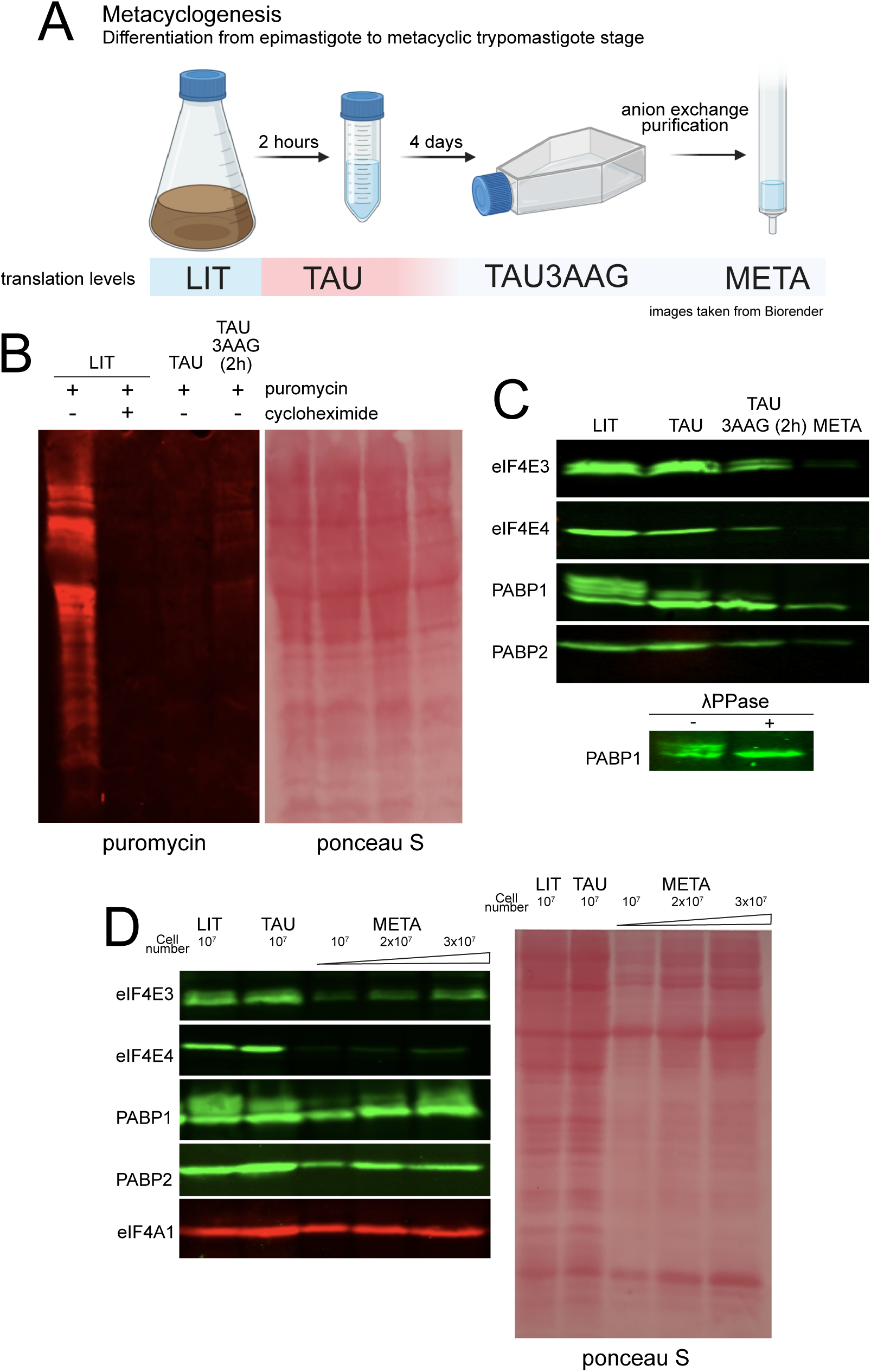
Expression of the proteins of interest and analysis of translation levels during differentiation. (A) Scheme of differentiation from epimastigotes to metacyclic trypomastigotes, a process known as metacyclogenesis. Epimastigotes cultured in a carbon-rich media (LIT) are concentrated and transferred to an acidic medium lacking carbon sources (TAU) for two hours and later in TAU supplemented with three amino acids (TAU3AAG, proline, glutamate, aspartate) for 4 days. An affinity purification by anion exchange chromatography is necessary to purify fully differentiated metacyclic trypomastigotes. Translation levels are reduced in TAU and there is a small return in translation in the first two hours of TAU3AAG [40]. It is unknown how translation is regulated during the four days incubation and quantitative measures of translation in metacyclics are missing. Images were taken from Biorender. (B) SUnSET assay (puromycin incorporation) in cells in LIT, TAU for 2h and TAU3AAG for 2h (after 2h of TAU). Puromycin incorporation detected with anti-puromycin antibodies in western blotting. Ponceau was used to check for loading. Image of one experiment. (C) Expression of proteins of interest in epimastigotes in LIT, TAU and TAU 3AAG and metacyclic trypomastigotes. Image of one experiment. Confirmation of PABP1 phosphorylation by treatment of the lysate from cells in LIT media with lambda phosphatase and detection by western blotting. Image of one experiment. (D) Expression of proteins of interest in epimastigotes in LIT, TAU and metacyclic trypomastigotes with increased number of cells. Ponceau was used to check for loading. Image of three independent experiments.

Translational control is important for stage differentiation in trypanosomatids, as the differences in mRNA translation efficiency between stages are much greater than the respective differences in mRNA abundances [44], [45], [46]. We previously compared the translational efficiency of mRNAs between the replicative (epimastigote) and non-replicative (metacyclic trypomastigote) stages by ribosome profiling (Ribo-seq): mRNAs encoding stress-responsive proteins are preferentially associated to ribosomes in the metacyclic stage, while mRNAs encoding for ribosomal proteins and proteins involved in metabolism are polysome-associated mainly in the epimastigote stage [44]. We recently extended this analysis to include one intermediate stage (epimastigotes stressed in TAU for two hours) and found better correlation between Ribo-seq of stressed epimastigotes and metacyclics than stressed and growing epimastigotes (manuscript in preparation). This finding suggests the existence of a pre-programmed mechanism for efficient global translation arrest and selective translation of stress– and differentiation-specific mRNAs. [47][40][47]. However, the mechanisms promoting translation of selected mRNAs during differentiation are not clear. In this work, we investigated whether and how the distinct *T. cruzi* eIF4F complexes are involved in this regulation, focusing on the essential proteins eIF4E3 and eIF4E4 [22]. We found two co-existing eIF4F complexes, which associate with specific proteins and mRNAs, but not with stress-responsive mRNAs, indicating this function may lie with another eIF4E. Our data also contributes to the understanding of the universal function of eIF4E upon nutritional stress.

## 4. Materials and methods

### Plasmids and cloning

The gene sequences of the translation factors used in this work were retrieved from the TriTrypDB database (https://tritrypdb.org/tritrypdb/app) and their respective accession numbers are given between parentheses. Full-length or truncated genes were amplified by PCR from *Trypanosoma cruzi* Dm28c genomic DNA using the primers described in **Supplemental Table 1**. The genes were expressed comprising the following amino acids sequences: eIF4E3_1-457_, eIF4E3_1-254_ (TcCLB.508827.30 or C4B63_6g269), eIF4E4_1-410_ (TcCLB.509037.40 or C4B63_42g204 and C4B63_42g205), PABP1_1-566_, PABP1_204-409_, PABP1_487-566_ (TcCLB.506885.70 or C4B63_2g331), PABP2_1-550_, PABP2_92-205_, PABP2_467-550_ (TcCLB.508461.140 or C4B63_2g174), eIF4A1_1-404_ (TcCLB.510155.180 or C4B63_85g83) and eIF4G4_1-678_ (TcCLB.510285.100 or C4B63_40g97). Please note the primers were designed based on the CL-Brenner Non-Esmeraldo like genome [47] since eIF4E4 is annotated as two partial fragments in the Dm28c 2018 genome (C4B63_42g204 and C4B63_42g205) due to an incorrect start codon [48]. This is supported by sequence analysis of all eIF4E4 orthologs found in kinetoplastids (data not shown).

The DNA sequences encoding eIF4E3_1-447_, eIF4E4_1-410_, PABP1_1-566_, PABP2_1-550_, eIF4A1_1-404_, and eIF4G4_1-678_ were amplified by PCR and cloned into the entry plasmid pDONR211^TM^ of the Gateway cloning system (Invitrogen). The coding sequences of eIF4E3_1-447_, eIF4E4_1-410_, eIF4A1_1-404_ PABP1_1-566_ and PABP2_1-550_ were subcloned from pDONR211^TM^ into various destination vectors of the Gateway system for various applications. pDEST17^TM^ (Invitrogen) was used for eIF4E3_1-447_, eIF4E4_1-410_ and eIF4A1_1-404_ expression with a 6xhis tag. eIF4E3_1-447_, eIF4E4_1-410_ were subcloned into pTcGW version 2.0 for episomal expression in *T. cruzi* of the proteins with a GFP tag at the C-terminus (pTcGFPN-CO) [49]. eIF4E3_1-447_, eIF4E4_1-410_, PABP1_1-566_, PABP2_1-550_ and eIF4G4_1-678_ were subcloned into pDEST22^TM^ and pDEST32^TM^ (Invitrogen) for N-terminal tagging with DNA binding and activation domains used in yeast two hybrid assays. All cloning steps were performed according to manufacturer’s protocol (Invitrogen). eIF4E3_1-254_, PABP1_204-409_ and PABP2_92-205_ were cloned into pET28a (Novagen) with a 6xhis tag for protein expression and purification. PABP1_487-566_ and PABP2_467-550_ were cloned into pGEX-4T-2 (Amersham) with a GST amino-terminal fusion for pulldown assays.

### *Trypanosoma cruzi* culture

Epimastigotes form the *Trypanosoma cruzi* Dm28c strain [50] were cultured in liver infusion tryptose (LIT) medium with 10% heat-inactivated fetal bovine serum (FBS) in axenic conditions at 28 °C [38], [51]. Cultures were kept in logarithmic growth by continuous sub-culturing at a density of 10^6^ cells/mL every three days. Analyses of exponential growing epimastigotes were made on the third day of growth at density of 2 – 3 x 10^7^ cells/mL. Cells were counted on a Z2 Coulter counter (Beckman Coulter). Transfection of pTcGFPN-CO containing eIF4E3_1-447_ and eIF4E4_1-410_ was performed as described previously [52] and transfectants selected with neomycin (500 μg/ml), followed by confirmation by flow cytometry and western blotting.

Epimastigotes were stressed in triatomine artificial urine (TAU) [38]. For this, exponentially growing epimastigotes were harvested by centrifugation, washed once in PBS and suspended at a density of 5 x 10^8^ cells/mL in TAU and incubated for two hours at 28 °C. Epimastigotes stressed in TAU were further stressed in TAU3AAG as follows: TAU-stressed cells were harvested by centrifugation and resuspended at a density of 5 x 10^6^ cells/mL in TAU3AAG for two hours at 28 °C. Metacyclic trypomastigotes were differentiated *in vitro* from epimastigotes cultivated in LIT medium with 5% FBS, 5% CO_2_ at 28 °C for twelve days [53]. Metacyclic trypomastigotes were purified by diethylaminoethylcellulose (DEAE-cellulose, Sigma) chromatography [38].

### Production of polyclonal antibodies and western blotting

eIF4E3, eIF4E4, PABP1, PABP2 and eIF4A1 antibodies were raised by immunisation of mice and rabbits with recombinant proteins. Proteins were expressed in various *Escherichia coli* strains, affinity purified with Ni-NTA resin (Qiagen) according to manufacturer’s protocol and quantified by SDS-PAGE analysis relative to a bovine serum albumin standard (Thermofisher). Rabbit immunization was performed by the “Célula B Serviço de Produção de Anticorpos” company (Federal University of Rio Grande do Sul, Brazil). Mice were immunised at the Animal Facility of the Carlos Chagas Institute within the protocol P-47/12-3 with the license number LW-19/19 authorized by the ethics committee for animal experimentation of FIOCRUZ.

Rabbit antibodies were affinity purified by immunoadsorption as follows: 100 μg of recombinant protein was transferred onto a nitrocellulose membrane, stained with Ponceau and excised. The membrane was blocked with 5% milk in PBS for one hour, washed three times in PBS for 5 minutes and rotated with 1 mL of rabbit sera at 4 °C overnight. Membrane was washed four times in 0.05% Tween in PBS (PBS-T) for 5 minutes. Antibodies were eluted with 200 μL of 100 mM glycine pH 2.5, the pH adjusted with 10 μL of 1 M Tris HCl pH 8.0, quantified as IgG in Nanodrop One (Thermofisher) and stored at –20°C until further use.

For western blotting, proteins were resolved by SDS-PAGE, transferred onto Amersham™ Protean® Premium Western blotting membranes (Merck) and blocked with 5% low fat milk in PBS for one hour and immunoblotted with the primary antibodies at 4 °C with gentle rocking overnight. The dilution of rabbit purified antibodies in PBS-T was 1:100 for eIF4E3, eIF4E4 and PABP2 and 1:1000 for PABP1. Mice sera for and eIF4A1 were diluted at 1:5000. After washing three times with PBS-T for 5 minutes, the membranes were incubated in PBS-T with IRDye™ 800LT anti-rabbit IgG or IRDye™ 680LT anti-mouse IgG (Li-Cor Cooperate) (both at 1:20,000 dilution) for one hour. The membranes were washed three times with PBS-T and one with PBS for 5 minutes and scanned using an Odyssey® CLx apparatus (Li-cor Cooperate). The ImageStudio™ software (Li-Cor Cooperate) was used to quantify protein bands.

### Translation assay

Translation was assayed by using the SUnSET method [54] as follows; 3 x 10^8^ epimastigotes or stressed epimastigotes were treated with 2 mM puromycin for 20 minutes at 28 °C, washed twice in PBS and lysed in sample buffer. Proteins were resolved by SDS-PAGE, transferred onto nitrocellulose membranes and checked by Ponceau staining. Puromycin incorporation was probed with Anti-Puromycin Antibody (clone 12D10, Sigma,1:5000 dilution) as described in western blotting.

### Dephosphorylation assay

3 x 10^8^ exponential growing epimastigotes were harvested, washed once in PBS and lysed in 500 μL of lambda protein phosphatase buffer (20 mM Tris-HCl, 150 mM NaCl, 1mM EDTA, 1mM EGTA, 1% Triton X-100 and 1mM MnCl_2_ and protease inhibitor cOmplete™, Mini, EDTA-free Protease Inhibitor Cocktail) on ice. 500 μL of lysate was treated with lambda protein phosphatase (the plasmid encoding the phosphatase was a kind gift from Dr. Stênio Perdigão Fragoso, Carlos Chagas Institute, FIOCRUZ, Brazil) for 30 minutes at 30 °C. The lysate was diluted in sample buffer, boiled for 5 min at 95 °C and probed in western blotting (PABP1 and eIF4E3).

### Yeast two-hybrid assays

The haploid yeast host strains Y8800 (MATa) and Y8930 (MATα) [leu2-3,112 trp1-901 his3Δ200 ade2-101 ura3-52 gal4Δ gal80Δ GAL2::ADE2 GAL1::HIS3@LYS2 GAL7: LacZ@met2 cyh2R] used in two-hybrid analyses contains yeast HIS3, LacZ and ADE2 genes integrated into the genome as two-hybrid interaction reporters. The Y8800 strain was transformed with the plasmid pDEST^TM^22, which encodes the activation domain (AD) fusions. The Y8930 strain was transformed with plasmid pDEST^TM^32 encoding the DNA binding domain (DBD) fusions. Transformation was done with the TE/PEG/lithium acetate method with salmon carrier DNA [55]. Diploid yeast strains were generated by mating on YEPD plates and selected on synthetic complete (SC) medium in the absence of leucin and tryptophan. For the phenotypic assay, the diploid strains were transferred to plates containing SC medium and selections: –Leu-Tryp-Ade and –Leu-Tryp-His + 5mM 3AT and 10mM 3AT. Gateway® control vectors were used as assay controls (pEXP^TM^32/Krev (BD) + pEXP^TM^22/RalGDS-wt (AD): positive control and pEXPTM32/Krev (BD) + pEXPTM22/RalGDS-m2 (AD): negative control). Plating was performed using an automated Tecan Freedom 200 system.

### Protein expression, purification, and in vitro pulldown assays

The N-terminal region of eIF4E3 (eIF4E3_1-254_ or eIF4E3N) as well as the C-terminal domain of PAPB1 (PABP1_487-566_ or PABP1C) and PABP2 (PABP2_467-550_ or PABP1C), the latter two in fusion with GST, were expressed in *Escherichia coli* BL21 (DE3) slyD^-^ cells [56] carrying the plasmid pRARE2 (Novagen). Recombinant GST (vector pGEX-4T2 (Amersham)) was also produced to be used as control. Cells transformed with the respective expression vectors were grown at 37°C in LB medium plus the selection antibiotics. Expression was induced at 18°C (OD600 ∼0.7) for sixteen hours with 0.25 mM isopropyl-β-D-thiogalactopyranoside (IPTG). Cells from 100 mL culture corresponding to the expression of GST-PABP1_487-566_, GST-PABP2_467-550_ and GST, respectively, were harvested by centrifugation, resuspended in 2.5 mL of PBS buffer pH 7.3 (140 mM NaCl; 2.7 mM KCl; 10 mM Na_2_HPO_4_; 1.8 mM KH_2_PO_4_), supplemented with EDTA-free protease inhibitor cocktail (Roche cOmplete™) and 10 mM 2-Mercaptoethanol (BME). After incubation with lysozyme (150 µg/mL), cells disruption was completed by sonication and the extracts were clarified by centrifugation at 20,000 x g for 30 minutes at 4°C. The soluble fractions were incubated with 100 µL of Glutathione Sepharose 4B resin (Cytiva) equilibrated with PBS + 10 mM 2-Mercaptoethanol, for one hour at 4°C under slow agitation. Pull-down assays were performed by gravity-flow chromatography using empty polypropylene columns. After collecting the unbound fraction, the resin was washed five times with 500 µL of PBS + 10 mM BME and one time with 500 µL buffer A (50 mM Tris-HCl, 100 mM NaCl pH 8.8) + 10 mM BME. In parallel, cells from 3 x 100 mL culture corresponding to the expression of eIF4E3_1-254_ were harvested by centrifugation, resuspended in (3 x) 2.5 mL (2.5 mL for each pull-down assay) of buffer A + 10 mM BME and disrupted following the same procedure described above. The soluble fractions were incubated, respectively, with Glutathione Sepharose 4B resin with immobilized GST-PABP1_487-566_, GST-PABP2_467-550_ and GST for one hour at 4°C under slow agitation. After collecting the unbound fraction, the resin was washed three times with 500 µL and one last time with 150 µL of buffer A. Target proteins were eluted with two times 150 µL of buffer B (50 mM Tris-HCl pH 9.0, 10 mM glutathione).

### Preparation of lysates for immunoprecipitation assays

All assays were performed with lysates obtained by nitrogen cavitation, a physical method of lysis where cells are resuspended in a hypotonic buffer, without any detergent. It has been used for translation *in vitro* assays in *Leishmania tarantolae* and we believe it is the best method to preserve the protein-protein interactions necessary for translation [57]. In this work, epimastigotes, stressed epimastigotes (2h TAU) or purified metacyclic trypomastigotes (10^10^ cells) were resuspended in 10 mL of lysis buffer (20 mM HEPES KOH pH 7.6, 75 mM potassium acetate, 4 mM magnesium acetate, 2 mM DTT) supplemented with one tablet of cOmplete™, Mini, EDTA-free Protease Inhibitor Cocktail and phosphatase inhibitor cocktail PHOstop™ (Roche). Subsequently, transferred to a Parr cavitation chamber (Parr Instrument Company, IL, USA), set at 1000 psi and kept on ice for 40 min. After sudden decompression, the lysates were centrifuged at 17,000 x g for 10 minutes at 4 °C to remove cell debris and used straight away in immunoprecipitation assays.

### Immunoprecipitation assays

We employed two strategies for the immunoprecipitation assays:

1) Lysates from wild type cells and capture of eIF4E3 and eIF4E4 using affinity-purified rabbit antibodies coupled to magnetic beads Dynabeads™ Protein A (Thermofisher). A commercial, non-reactive isotype immunoglobulin Rabbit IgG Polyclonal Antibody (Merck) was used as a negative control.
2) Lysates from cells expressing eIF4E3-GFP and eIF4E4-GFP fusion proteins and capture of eIF4E3-GFP and eIF4E4-GFP using a llama antibody fragment that recognizes GFP (clone LaG-16–G_4_S–LaG-2) [58] coupled Dynabeads™ M270-epoxy (Thermofisher). Lysates from a cell line with episomal expression of GFP-FLAG was used as a negative control [59].

In strategy 1, we prepared fresh beads for each experiment. For each assay, 10 μg of affinity-purified rabbit polyclonal antibodies were incubated with 50 μL of Dynabeads™ Protein A for one hour according to manufacturer’s protocol. Beads were washed once in PBS Tween 0.1% and resuspended in immunoprecipitation buffer (20 mM HEPES KOH pH 7.6, 75 mM potassium acetate, 4 mM magnesium acetate). In strategy 2, the antibody fragment was expressed in *E. coli* and purified as described in Fridy et al [58], and coupled to Dynabeads™ M270-epoxy (Thermofisher) as described by Obado et al [60] and stored in 4 °C until use.

Immunoprecipitation assays were performed with 1 mL of lysate incubated with the indicated beads for 2 hours at 4 °C with rotation. After incubation, the beads were washed three times with 1 mL of lysis buffer on ice. Subsequently, the beads were processed differently depending on the analysis. To identify proteins associated to eIF4E3 and eIF4E4, we resuspended the beads in sample buffer, boiled for 5 min at 95 °C and analysed by mass spectrometry. To identify mRNAs associated to eIF4E3 and eIF4E4, RNA was extracted using Trizol (Thermofisher), quantified with Qubit RNA HS Assay Kit (Thermofisher) according to manufacturer’s protocol and stored at –80°C until library preparation. All assays were done in technical triplicates.

To test the stringency of our lysis buffer, we compared one extra replicate done exactly as described with another replicate with a stringent buffer (HEPES 20 mM pH 7.4, sodium citrate 50 mM, magnesium chloride 1 mM, calcium chloride 10 mM). Cells were lysed by nitrogen cavitation with this buffer and lysate was supplemented with glycerol 10%, Triton X-100 0.1% and RNAse A 40 μg/ml.

### Mass spectrometry analysis

The samples from the boiled beads were loaded on SDS-polyacrylamide gels and run until the sample migrated approximately 1 cm inside the separating gel. Gel slices corresponding to each sample lane were excised and stained with Coomassie colloidal blue (Sigma). All steps from in-gel digestion and mass spectrometry data acquisition were carried out at the mass spectrometry platform of the Carlos Chagas Institute, FIOCRUZ-Paraná, Brazil. The gel slices were de-stained, then dehydrated in 100% ethanol for 10 min at 25 °C and dried in a speed-vac for 7 min. After this, the proteins were reduced (10 mM DTT in 50 mM ammonium bicarbonate, ABC) for 60 min at 56 °C and alkylated (55 mM iodoacetamide in 50 mM ABC) for 45 min at 25 °C protected from light. Digestion buffer (50 mM ABC) was added for 20 min at 25 °C, followed by dehydration in 100% ethanol for 10 min at 25 °C. The gel slice dehydration step was performed twice with subsequent drying in speed-vac for 7 min. In sequence, the slices were rehydrated in trypsin solution (trypsin 12,5 ng/μL in 50 mM ABC) for 20 min at 4 °C followed by removal of the supernatant and incubation at 37 °C by 16-18 h in digestion buffer. After trypsinization, peptides were extracted twice from gels through incubation in extraction buffer (3% trifluoroacetic acid and 30% acetonitrile) for 10 min at 25 °C. Samples were dried in speed-vac to 10-20% of the original volume and the peptides were purified using Stage-Tips-C18. The peptides were eluted in 40 μL formic acid 0,1%, acetonitrile 40%, dried in speed-vac for 30 min, diluted in formic acid 0,1%, DMSO 5%, acetonitrile 5% and analysed by LC-MS/MS on a LTQ Orbitrap XL mass spectrometer (Thermo Fisher Scientific).

The collected spectra were analysed by PSM with the Andromeda algorithm embedded into the MaxQuant platform [61]. Mass spectrometry analyses were performed using two technical replicates (runs) for each sample, to increase sequencing coverage. The genome of *T. cruzi* strain Dm28c 2018 (release 55) available at the TriTrypDB database was used in the searches for protein identification. The search parameters were set as default, where carbamidomethylation on cysteine was set as a fixed modification, oxidation on methionine and acetylation on protein N-terminus were set as variable modification. In the search, a minimum peptide length of 7 amino acids was required and 1% FDR was applied both at the peptide and protein levels. During the first search in the identification of peptides, tolerance of 20 ppm was used. Trypsin digestion sites were used with up to 2 missed cleavages.

Identification of potential interactors e.g., proteins enriched in the immunoprecipitations relative to the controls was done as described in [62]. Briefly, LFQ intensities were log_2_-transformed and missing values imputed from a normal distribution of intensities. A *Student’s t-test* compared the values of the three replicates of immunoprecipitation to the three replicates of the control. Criteria for potential interactors was four times enrichment in the immunoprecipitation (log_2_(FC) > 2) with statistically significance of *p*-value < 0.01 (-log_10_(*p*-value) > 2). Due to poor annotation of the Dm28c 2018 genome, several proteins of the 43S PIC complex were incorrectly annotated as conserved hypothetical. Their identification was done using the syntheny and orthology analysis with the Trypanosoma brucei 927 genome [63] available in the TriTrypDB [64].

### RNAseq analysis

The libraries were constructed with TruSeq Stranded mRNA kit (Illumina), prepared according to the manufacturer’s instructions, with adaptations. Briefly, we skipped the purification and fragmentation steps due to the already purified and enriched samples. All the samples were prepared in three independent replicates. RNAseq was performed on an Illumina HiSeq 2500 (Illumina, single-end 50-bp SR mid output run) on the High Performance Sequencing Platform (Oswaldo Cruz Institute, FIOCRUZ, Brazil). Sequencing reads were mapped against the *T. cruzi* Dm28c genome (Ensembl Assembly ASM317710v1) with STAR v2.7.10a [65] with default parameters. Gene counts were obtained for uniquely mapped reads with featureCounts v2.0.1 from the Subread package [66]. Whenever uniquely mapped read counts were set to zero due to duplicated regions or multi-mapped reads, we further verified these regions within the multi-mapped read counts available with featureCounts. To identify mRNAs enriched in the immunoprecipitation eluate relative to the control, the differential expression analysis DESeq2 v1.34.0 [67] package in R was used with uniquely mapped counts as input. Criteria for enrichment was a two-fold change (log_2_(FC) > 1) with False Discovery Rate < 0.05 (adjusted *p-value*). To calculate changes in eIF4E:mRNA association (assay) during stress (condition), we used the likelihood ratio test (LRT) to compare the full (∼ assay + condition + assay:condition) to the reduced model (∼ assay + condition) to identify significant genes (FDR < 0.05) with a Wald test (see Table S8 for the script used). mRNAs with FC > 0 were more associated in stress while < 0 less associated.

## 5. Results

### PABP1 is dephosphorylated when global translation is inhibited

Translation is highly regulated during differentiation from the replicative epimastigotes to the non-replicative metacyclic trypomastigotes. *In vitro*, a first step in this process can be achieved by transferring cells from carbon-rich LIT media to acidic carbon-poor TAU medium and exposed to nutritional stress for two hours. Subsequently, the cells are kept in TAU supplemented with an amino acid-/carbon-source (TAU3AAG) for five days. A purification step is necessary to isolate fully differentiated metacyclic cells (Figure 1A). To understand how differentiation is triggered, cells are analyzed after two hours in TAU and after the first two hours in TAU3AAG. After two hours in TAU, global translation is arrested; two hours later in TAU3AAG, translation returns to some extent, conceivably for differentiation-associated mRNAs. This was demonstrated with polysome gradients previously [40] and we now confirm this observation using the SUnSET method [54]. We observed a strong, cycloheximide-sensitive signal in the control, which is lost in TAU, and some signal in TAU3AAG cells, confirming that the *in vitro* metacyclogenesis protocol causes the expected regulation in translation levels (Figure 1B).

Abundance of eIF4E3, eIF4E4, PABP1, PABP2 and eIF4A1 during *in vitro* metacyclogenesis was analyzed by western blot, using antibodies produced against the recombinant proteins. In the case of PABP1 and PABP2, to avoid antibody cross-reactivity [68], truncated versions of the proteins, including only low identity regions, were used to inoculate the animals. Antibody specificity was subsequently validated (Supplementary Figure S1). Our results show that there is no apparent variation in protein levels between control cells and TAU-starved cells, for any of the initiation factors analyzed, while their levels were reduced in TAU3AAG and even more in metacyclics (Figure 1C and Figure 1D).

Interestingly, eIF4E3 and PABP1 migrated as multiple bands, indicating post-translational modifications. For eIF4E3, the band pattern did not change during metacyclogenesis, but the higher molecular weight (MW) bands of PABP1 are absent in TAU, TAU3AAG or metacyclics. eIF4E3 and PABP1 are known phosphoproteins in *T. brucei* [69]. We therefore treated cell lysates of epimastigote cells with lambda phosphatase and probed for eIF4E3 and PABP1 on a western blot. We could detect the loss of the higher MW bands of PABP1 and thus confirm that PABP1 is a phosphoprotein in *T. cruzi* (Figure 1C); eIF4E3 was not detected after incubation and was likely degraded. To distinguish whether PABP1 dephosphorylation is caused by differentiation or by translational repression, we allowed epimastigotes to grow from log phase to stationary phase. This reduces translation without inducing differentiation [42] (Figure 2A). PABP1 was phosphorylated in log phase and dephosphorylated in stationary phase, indicating that PABP1 phosphorylation correlates to translational repression rather than differentiation (Figure 2B). When cells are incubated in TAU3AAG and some carbon sources are available, translation returns in small levels, expectedly for stress-specific mRNAs, but PABP1 phosphorylation does not return.

**Figure 2.**
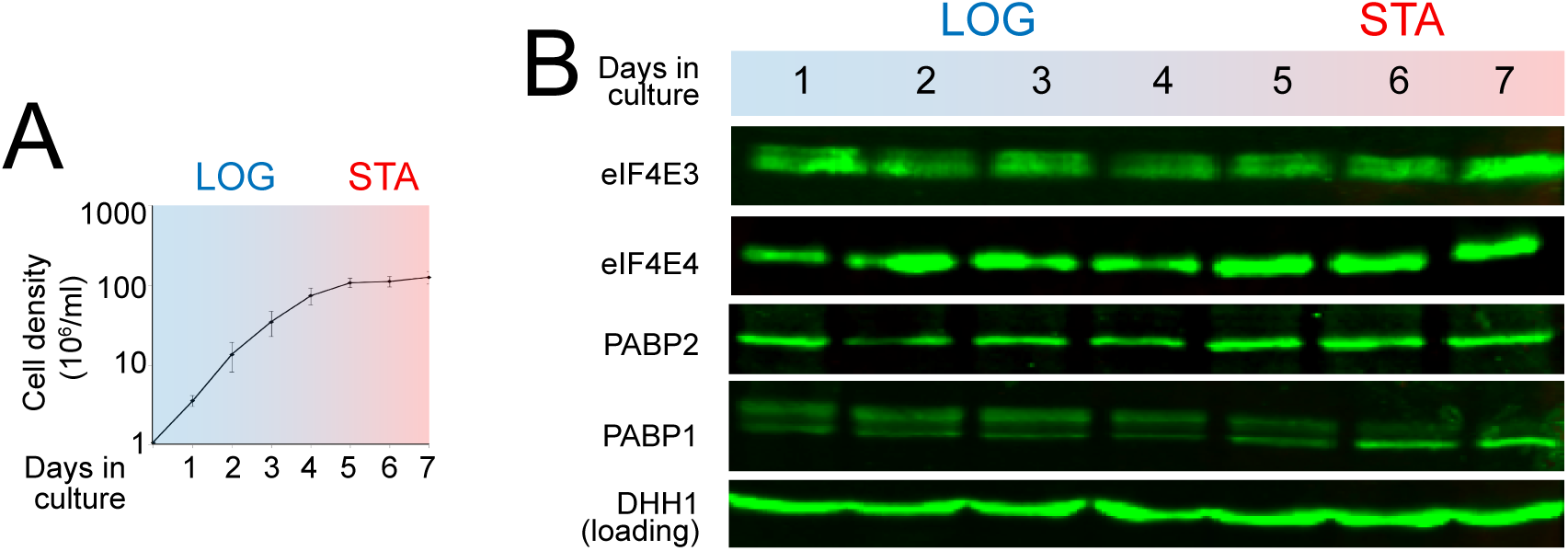
Expression of proteins of interest in a growth curve. (A) Epimastigotes grow in log phase until the 3-4^th^ day and from the 5^th^ day onwards transition to the stationary grow phase. (B) eIF4E3, eIF4E4, PABP1, PABP2 and DHH1 were detected by western blotting. Representative images of one experiment.

### The eIF4E3 amino-terminal extension binds to the MLLE domain of PABP1 and PABP2

There is extensive evidence for interaction of eIF4E4 with PABP1 [24], [31]. No experimental evidence exists for the direct interaction between eIF4E3 and PABP1/2 even though the interaction motifs are conserved. Analysis of several trypanosomatid eIF4E3 sequences (Figure S3A, S3B) shows conservation of the PAM2 motifs “A” and “B” and loss or degeneration of “C” outside of the *Trypanosoma* genus. This shows the potential of eIF4E3 to bind PABP1 and/or PABP2 through a similar mechanism to that used by eIF4E4. Two different methods were used to test the interaction between eIF4E3 and the PABPs, yeast-two hybrid screen and *in vitro* pulldown assay.

The yeast two-hybrid assay was used to test all possible interactions between eIF4E3, eIF4E4, eIF4G3, eIF4G4, PABP1 and PABP2. The proteins were expressed with N-terminal fusions of either the activation domain (AD) or the DNA-binding domain (DBD) of the split GAL4 transcription factor. The fusions were transformed into the haploid strains Y8800 and the DBD into the haploid strains Y8930. Mating between Y8800/Y8930 resulted in the diploid Y strain expressing both fusion proteins. The interactions were tested for the HIS3 reporter in the presence of two concentrations of the inhibitor 3-AT. As expected, we observed growth of the combinations eIF4E3/eIF4G4 and eIF4E4/eIF4G3, as well as of the positive and not of the negative control. We also observed growth of the combinations eIF4E3/PABP1 and eIF4E3/PABP2. We did not observe growth for eIF4E4/PABP1, a possible explanation is the effect of the N-terminal tag, which did not seem to affect eIF4E3 likewise (data not shown). To summarize, these experiments indicated direct interactions between eIF4E3 and both PABPs and confirmed all previously reported interactions (eIF4EG3/eIF4E4, eIF4EG4/eIF4G3 and eIF4E4/PABP1 [23], [24]) (Figure 3B).

**Figure 3.**
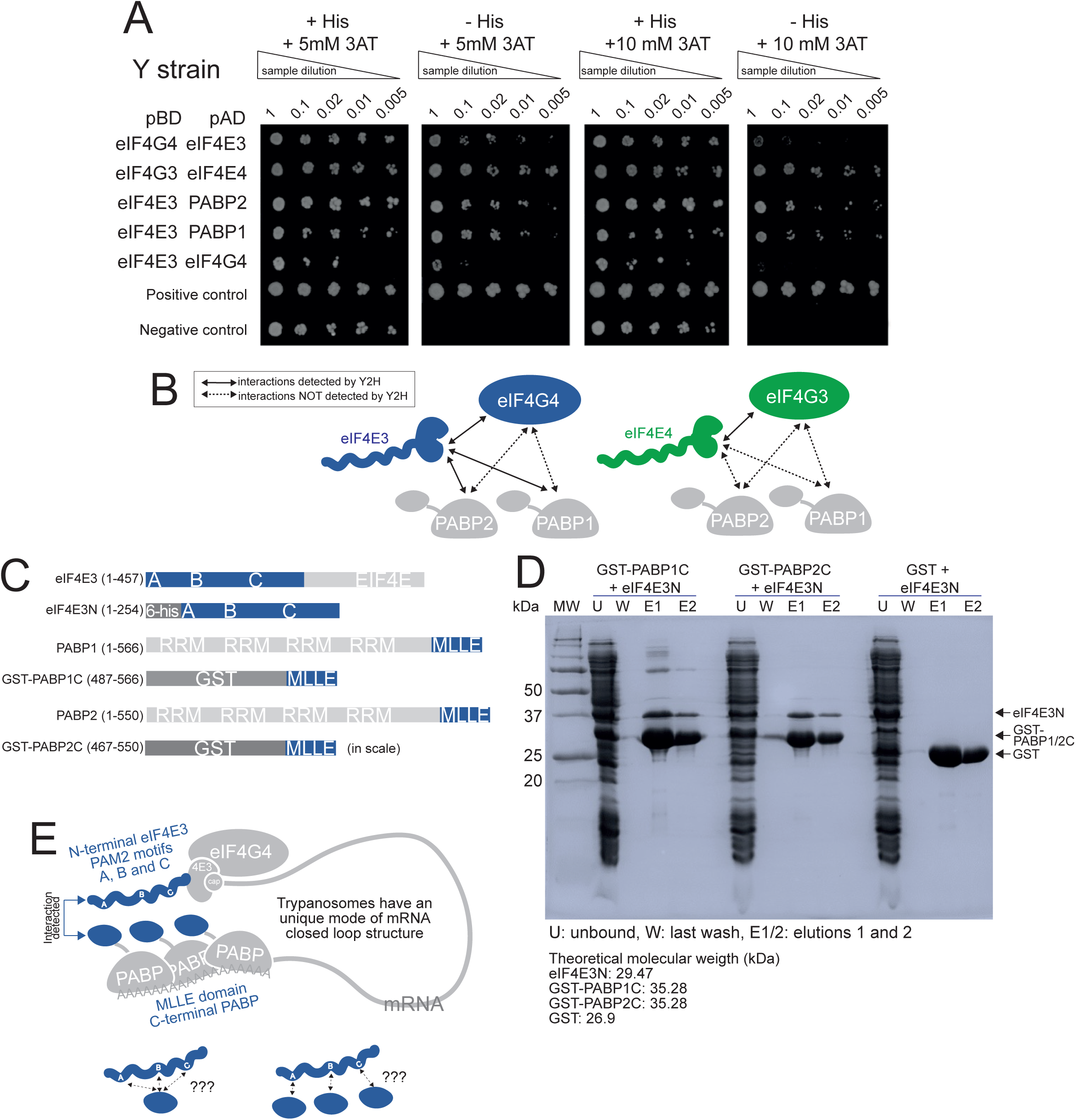
Two different methods support the eIF4E3:PABP1 and eIF4E3:PABP2 interactions. (A) All translation initiation factors were cloned in fusion with both the activation domain (pAD) and the DNA-binding (pBD) of plasmids pDEST^TM^22 and pDEST^TM^23, respectively. Only the positive interactions and the control strains are shown. Cells were cultured under restrictive (-His; –Ade) and non-restrictive (+His; +Ade) growth conditions. When indicated, the medium was supplemented with increasing amounts of 3-AT, increasing the stringency of the assay. Dilutions (0.1, 0.02, 0.01, and 0.005) of saturated yeast cultures containing the indicated plasmids were spotted on the plates, and photographs were taken after 3 days of incubation at 30 °C. Positive and negative control plasmids were supplied by the gateway’s manufacturer. Yeast two hybrid assays are positive for the interactions eIF4E3:eIF4G4, eIF4E3:PABP1, eIF4E3:PABP2 but negative for eIF4G3/4:PABP1/2 and eIF4E4:PABP1/PABP2 interactions. (B) Scheme of interactions detected or not detected by the yeast two hybrid assay. (C) Design of constructions for the pulldown assay. The regions included in the truncated versions of eIF4E3, PABP1 and PABP2 are highlighted in blue. The carboxi-terminal domain (MLLE) of both PABPs were expressed fused to glutathione-S-transferase (GST). (D) Results of the pulldown assays analyzed by SDS-PAGE stained with Coomassie. GST-fused PABPs or GST alone immobilized in glutathione-Sepharose resin were used as baits. After loading *E. coli* extracts containing recombinant eIF4E3N and washing the resin, GST containing samples were eluted with glutathione. Molecular weight (MW) of the markers are indicated in the left (kDa), U: unbound, W: last wash, E1/2: elutions. The theoretical molecular weights of the recombinant proteins are indicated. (E) Proposed model for trypanosomes mRNA closed loop structure mediated by the interaction between the PAM2 motifs in the amino-terminal extension of eIF4E3 and the carboxi-terminal (MLLE) domain in PABP1 or 2.

Next, we narrowed down the regions responsible for the interaction of eIF4E3 with PABP1/2 using *in vitro* pulldown assays. The following recombinant protein variants were produced: the eIF4E3 amino-terminal extension including the three PAM2 motifs (eIF4E3N) and the MLLE C-terminal domains of PABP1 and PABP2 fused to GST in their N-terminal (GST-PABP1C and GST-PABP2C) (Figure 3C). GST alone was also produced and used as a negative control. The GST-PABPs or GST bound to glutathione beads were used as bait. Our results show the interaction between the amino-terminal domain of eIF4E3 and the GST-fused MLLE of PABP1 and PABP2, but not with GST (Figure 3D).

We propose that the unique architecture of closed-loop mRNAs by direct interactions eIF4E4:PABP1 described in *Leishmania* [31] also occurs in *Trypanosoma* eIF4E3:PABP1/PABP2. The stoichiometry of this structure is unknown, but it would be unprecedent a single MLLE binding to three PAM2 [30]. Instead, three PAM2 in a single protein could have a regulatory effect, such as observed for eRF3 [70] and/or a concentration effect resulting in an enhanced eIF4E:PABP interaction.

### Under nutritional stress, eIF4E3 still associates to eIF4G4, but interactions to the 43S PIC are decreased

We purified eIF4E3 and its associated proteins by immunoprecipitation from wild type *T. cruzi* cells using antibodies to the endogenous protein. The immunoprecipitation was done in exponentially growing cells (LIT condition) or in cells under nutritional stress (TAU condition). As negative control, beads bound to a non-reactive IgG were used. The eluates were analyzed by mass spectrometry. A protein was considered associated to eIF4E3 if it had four times higher LFQ intensities in the eIF4E3 sample than in the negative control (*t*-test difference, in log_2_ > 2 and *p*-value < 0.01). We plotted the *t*-test differences of the experiments in LIT and TAU condition in a scatter plot (Figure 4A, Supplemental Figure S4/Table S3). The eIF4E3 and its partner, eIF4G4, are the proteins with the highest enrichment (*t*-test difference) values, indicating that the assay is successful in capturing eIF4E3 and its associated proteins. This was the case in both LIT and TAU conditions, a result which also shows that nutritional stress does not dissociate eIF4G4 from eIF4E3. Of the PABPs, only PABP1 was enriched in both conditions, while PABP2 is detected but below the enrichment cut-off. This was likely caused by a too low stringency in the immunoprecipitation conditions since high levels of PABP2 were found in the control; when we used higher salt and RNAse A in an additional replicate, we readily detected both PABPs in the eIF4E3 immunoprecipitation but not in control. This also supports that the interactions between eIF4E3, eIF4G4 and the PABPs are direct (Figure 4B).

**Figure 4.**
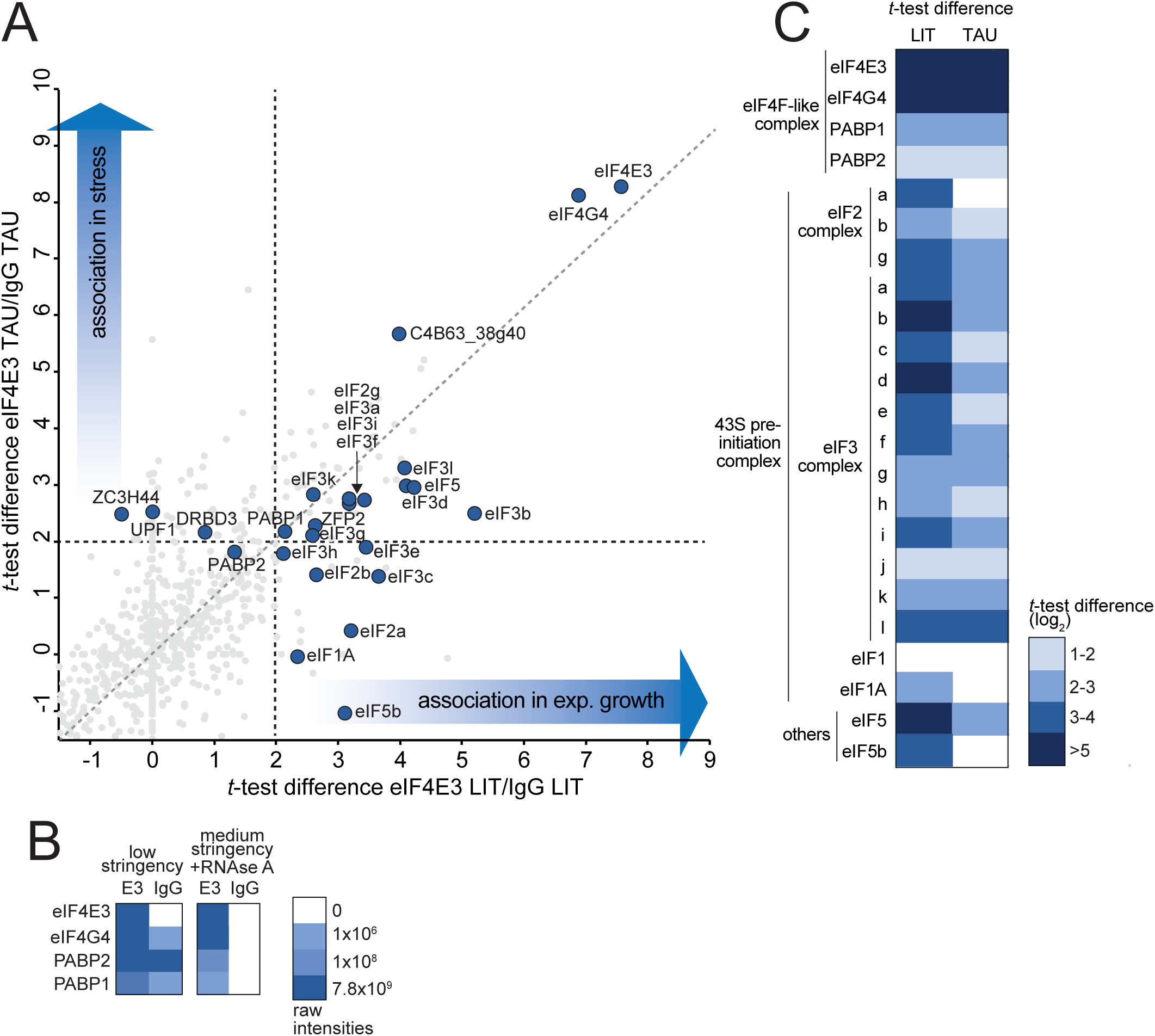
Proteins associated to native eIF4E3 in LIT and TAU stress. (A) Scatter plots generated plotting the *t*-test difference generated by comparing the LFQ-intensities of eIF4E3 and IgG in LIT (associated in exponential growth) and TAU (associated in stress) conditions in a Student’s *t*-test. (B) Heatmaps showing the raw intensities of the indicated proteins between two additional replicates, one with low stringency conditions (repeated with the exact protocol as in 4A) and the other with a stringent buffer and RNAse A. (C) Heatmaps showing the *t*-test differences (in log_2_) for the indicated proteins (eIF4F-like complex, 43S pre-initiation complex and other translation initiation factors) from the experiment in 4A.

Several other proteins were specifically enriched with eIF4E3 in the lower stringency buffer (Figure 4A), of these, 49 in both conditions, with 17 being specific to LIT and 55 specific to TAU. Some proteins are RNA binding proteins (RBPs), possibly bound to eIF4E3-associated mRNAs. Of these, UPF1, ZC3H44 and DRBD3 were specific to the TAU conditions, indicating a possible function of these proteins in stress conditions. The remaining proteins were analyzed based on the localization of their *T. brucei* orthologue that was determined in the genome-wide localization project TrypTag [71]; they mostly localize to the endocytic apparatus, suggesting that eIF4E3 might associate to the ER, possibly the rough ER, where translation of membrane-bound proteins take place.

A large fraction of the eIF4E3 associated proteins were proteins of the 43 pre-initiation complex. In exponential growth, almost all subunits of the eIF3 complex (a-l but j), subunits from the eIF2 complex (a, b and g), eIF1A (but not eIF1), eIF5 and eIF5b were enriched (Figure 4A and B). This is a strong indication that eIF4E3 is a ‘canonical’ eIF4E in trypanosomes. Nutritional stress caused a marked decrease in 43S association: proteins from eIF3 and eIF2 complexes are still enriched but with lower *t*-test difference values than in LIT conditions. eIF2a, b, eIF3c, e, h and eIF5b to eIF4E3 are still detected but below the threshold set for enrichment (Figure 4B). Similar findings were observed with a second immunoprecipitation strategy using GFP-tagged eIF4E3 and anti-GFP antibodies (Supplemental Figure 4/Table S3); the association between eIF4E3 and eIF4G4 was not affected by stress and the 43S PIC complex enrichment decreases in TAU. However, in this strategy, the 43S PIC complex proteins did not meet the enrichment criteria, perhaps as an effect of the GFP tag.

The decrease in association of eIF4E3 with the 43S PIC at nutritional stress correlates with the observed decrease in global translation, pointing to a function of eIF4E3 in this repression. Importantly, the eIF4F complex does not disassemble at starvation stress.

### eIF4E4 interacts with a different cohort of proteins than eIF4E3

We used the same immunoprecipitation strategy to identify proteins associated with eIF4E4 (Figure 5A and Supplemental Figure 5/Table S4). Next to the bait, the expected eIF4F complex proteins eIF4G3, PABP1 and eIF4A1 were among the highest-enriched proteins. PABP2 was not enriched, consistent with the eIF4E4/G3 complex being preferentially associated with PABP1 [24]. Similar to what was observed for eIF4E3, the interactions between the eIF4F subunits were not affected by nutritional stress; these were also not affected by an increased stringency and RNAse A in the additional replicate (Figure 5B).

**Figure 5.**
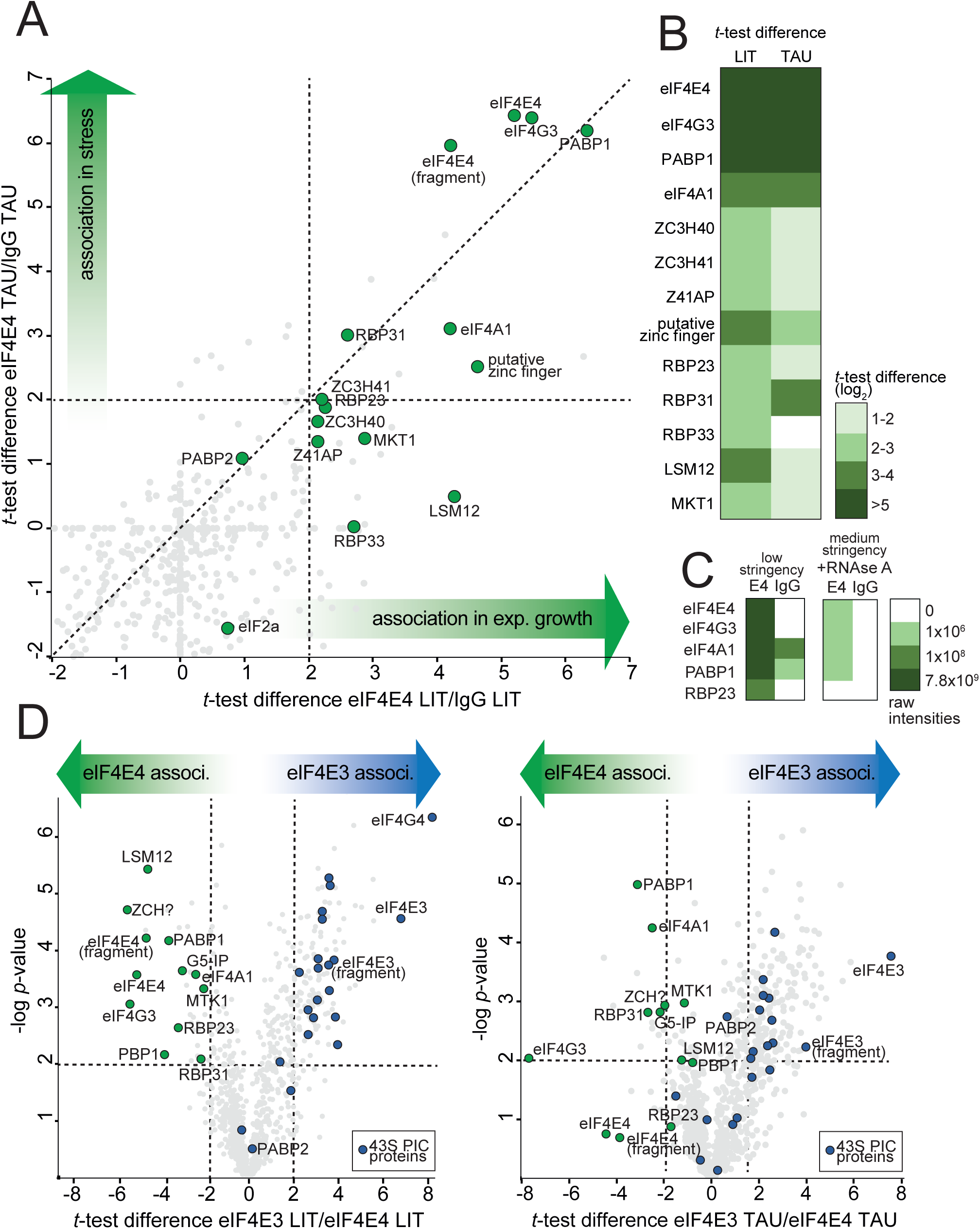
Proteins associated to native eIF4E4 in LIT and TAU stress. (A) Scatter plots generated plotting the *t*-test difference generated by comparing the LFQ-intensities of eIF4E4 and IgG in LIT (associated in exponential growth) and TAU (associated in stress) conditions in a Student’s *t*-test. (B) Heatmaps showing the raw intensities of the indicated proteins between two additional replicates, one with low stringency conditions (repeated with the exact protocol as in 5A) and the other with a stringent buffer and RNAse A. (C) Heatmaps showing the *t*-test differences (in log_2_) for the indicated proteins (eIF4F-like complex, 43S pre-initiation complex and other translation initiation factors) from the experiment in 5A. (D) Volcano plots generated plotting the t-test difference generated by comparing LFQ-intensities of eIF4E3 and eIF4E4 in LIT and TAU conditions in a Student’s *t*-test. 43S PIC proteins are the dots indicated without text.

Several further proteins were enriched with eIF4E4 in the lower stringency buffer (Figure 5A): 21 in both conditions, 13 specific to LIT and 6 specific to TAU. These included several RBPs: RBP23, RBP31, ZC3H40, ZC3H41, Z41AP, a putative zinc finger protein (C4B63_2g319) and two proteins involved in the regulation of RNA stability (MKT1 and LSM12 [72]). In nutritional stress, some RBPs (RBP23, ZC3H40, ZC3H41, Z41AP, RBP31, MKT1 and LSM12) have their *t*-test difference values decreased but just below the statistical cut-off. Only RBP33 is not detected anymore in nutritional stress conditions and likely loses its association with eIF4E4. In contrast to eIF4E3, we found no enrichment of the 43S PIC proteins. Although they were detected, their levels were decreased upon nutritional stress. An alternative immunoprecipitation experiment, using cells expressing eIF4E4-GFP and GFP-antibodies gave similar results (Supplemental Figure 4/Table S4).

Our data show that eIF4E3 and eIF4E4 associate with distinct protein cohorts. For further confirmation, we directly compared the protein associations between eIF4E3 and eIF4E4 in the *t*-test (Figure 5D, Supplemental Table S5). Proteins preferentially associated to each eIF4E will have large positive/negative *t*-test difference values while proteins with similar association (with interactions to both) or no association (contaminants) will have *t*-test difference values closer to zero. We found many proteins differentially associated to either eIF4E: 117/135 to eIF4E3 and 29/12 to eIF4E4 (LIT/TAU). eIF4E3, eIF4G4 and most of the 43S PIC proteins preferentially associate to eIF4E3. eIF4A1, eIF4G3, PABP1, RBP33, RBP23, C4B63_2g319 were specific to eIF4E4. The proteins MKT1, PBP1, LSM12, G5-IP, which form a recently described complex [73], were all preferentially associated to eIF4E4. In TAU, similar findings were observed, except that eIF4E4 itself did not meet the statistical threshold.

Combining all data from the immunoprecipitations of the two eIF4E, with two different capture strategies and in two experimental conditions, we describe a scenario where two different eIF4F-like complexes exist, showing preferential interactions to distinct cohorts of proteins, and thus likely have distinct functions.

### eIF4E3 and eIF4E4 associate to different cohorts of mRNAs and this association is retained in stress

The differences in associated proteins between eIF4E3 and eIF4E4 suggest that each could also have differential associations to mRNAs. To test this, we performed the same immunoprecipitation assays (GFP antibodies to GFP-fusions or specific antibodies to the wild type proteins) but analyzed the co-precipitated mRNAs. Sequencing data quality check, mapping statistics and read counts are available in Supplemental File 1. We considered an mRNA to be associated to an eIF4E if the mRNA was up-regulated in the eIF4E immunoprecipitation condition relative to the IgG or GFP-FLAG control more than two-fold (log_2_(FC) > 1) with a False Discovery Rate < 0.05 (adjusted *p-value*). The complete dataset can be found in Supplemental Figure S5/Table S6. Overall, a larger number of mRNAs was identified with the GFP-tagged proteins: 447/586 mRNAs were associated with eIF4E3, and 95/175 with eIF4E4 under LIT/TAU condition (Figure 6A, Supplemental Figure S4/Table S6). This could be explained by the increased expression levels of the bait due to the episomal expression and/or differences between the resins.

**Figure 6.**
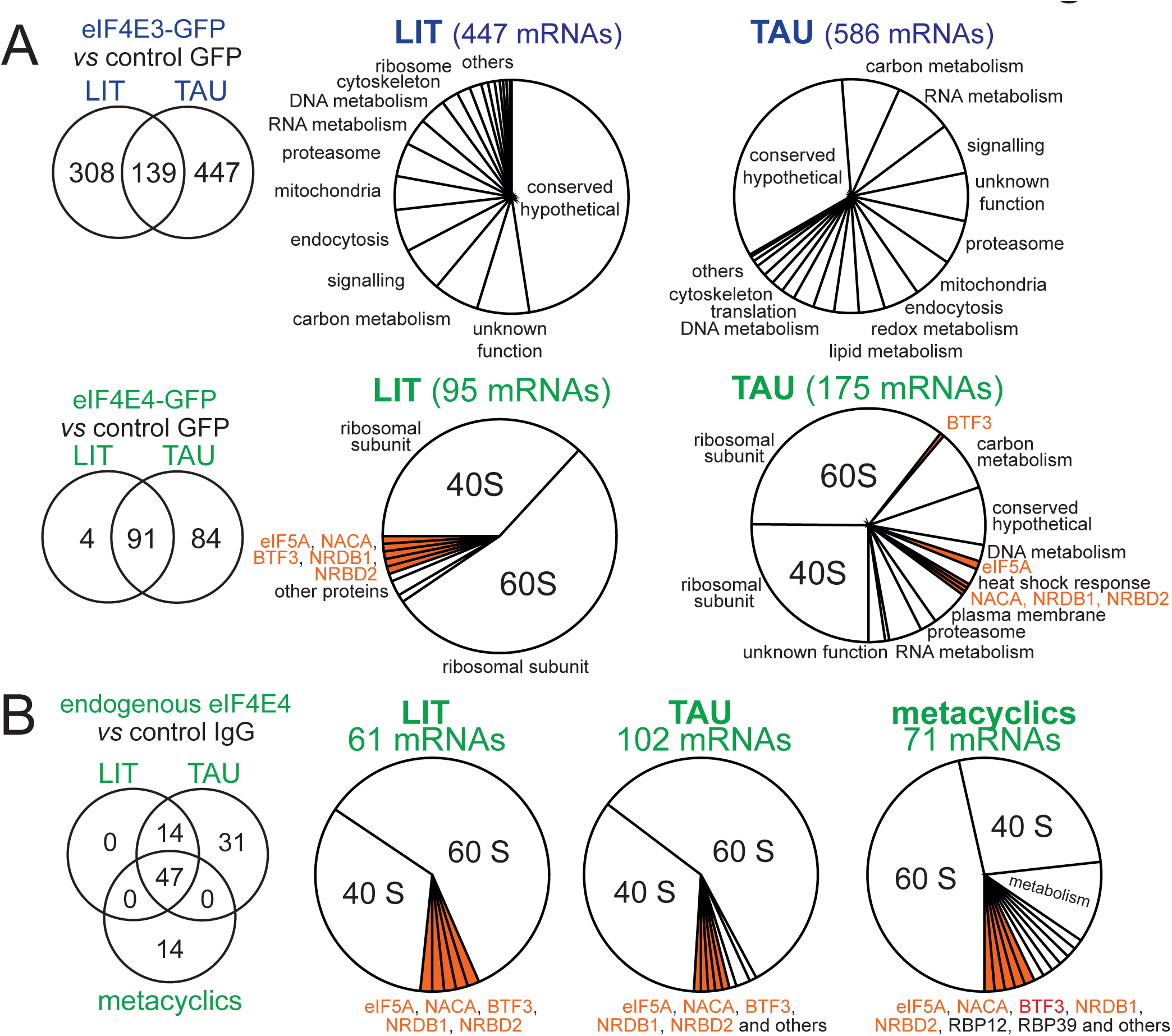
Description of eIF4E3 and eIF4E4 differentially associated mRNAs in LIT (associated in exponential growth) and TAU (associated in stress) conditions in a DESeq2 comparison. (A) A list of mRNAs differentially associated to eIF4E3/4-GFP relative to the GFP-FLAG control in LIT and TAU (log_2_ (FC) > 1, FDR < 0.05) was manually categorized based on the expected function of the mRNA using its “product description” annotation (Table S6). A pie chart shows the distribution of these categories for eIF4E3-GFP and eIF4E4-GFP in LIT and TAU. Venn diagram shows mRNAs common between these experiments. The mRNAs of ribosome-associated proteins, discussed in the text, are coloured orange. (B) A list of mRNAs associated to native eIF4E4 relative to the IgG control in LIT, TAU and metacyclics (log_2_ (FC) > 1, FDR < 0.05) was manually categorized based on the expected function of the mRNA using its “product description” annotation. A pie chart shows the distribution of these categories for native eIF4E4 in LIT, TAU and metacyclics. Venn diagram shows mRNAs common between these experiments. The mRNAs of ribosome-associated proteins, discussed in the text, are coloured orange.

For the eIF4E3 and eIF4E4 GFP-fusions strategy we manually categorized the associated mRNAs according to the (predicted) function of the encoded protein. About half of the eIF4E3 associated transcripts encode hypothetical proteins, and the other half proteins with diverse functions, with no obvious enrichment in a specific category. The data agree with a function of eIF4E3 as a regulator of bulk mRNA translation. The detection of only ∼500 mRNAs of the ∼15000 mRNAs encoded in the *T. cruzi* genome, is likely caused by the detection limit of the experiment. In fact, we found a positive correlation between eIF4E3-association and mRNA abundance, by a comparison with total poly(A)-enriched RNA seq data (Supplemental Figure S7). In contrast to eIF4E3, the eIF4E4-associated mRNAs were lower in number and encoded almost exclusively proteins of both ribosomal subunits and ribosome-associated proteins, indicating that eIF4E4 has specialized for translation of this specific group of mRNAs.

To our surprise, nutritional stress did not cause a decrease in the number of associated mRNAs, neither for eIF4E3 nor for eIF4E4, despite of the global translational arrest. In fact, the opposite was observed: the number of detected mRNAs increased 1.3/1.8 fold for eIF4E3/eIF4E4. There was no obvious change in the functional categories of mRNAs associated to eIF4E3. Likewise, eIF4E4 remained associated with ribosomal protein encoding mRNAs, but some other mRNAs became associated too. The data suggest that both eIF4Es remain associated with their mRNAs during the translational repression caused by nutritional stress, with eIF4E4 possibly gaining some mRNA targets.

We have repeated all experiments with antibodies against the endogenous protein, this time including the metacyclic life cycle stage, even though this stage has greatly reduced eIF4E4 levels (compare Fig. 1C). The assay is reproductible, mRNAs associated to eIF4E4 between the two independent experiments do overlap (Supplemental Figure S8 and Table S7). Focusing on the second experiment, we observed the preference for mRNAs encoding ribosomal proteins even in metacyclic trypomastigotes. In addition, five further mRNAs were shared for the three conditions, all encoding ribosome-associated proteins: eIF5A, the nascent associated complex subunit alpha (NACA), the basal transcription factor 3 (BTF3 also known as nascent associated complex subunit beta) and two nuclear RNA binding proteins 1 and 2 (NRDB1/2) (Figure 6D). While not *bona fide* ribosomal proteins, these can be implicated in ribosome function and structure. Their association to eIF4E4 indicates a preference beyond ribosomal mRNAs but also proteins implicated in ribosome structure and function. How this preference occurs is not clear.

### Stress-translated mRNAs are not associated with eIF4E3 and eIF4E4

Additionally, we analyzed how the association of a mRNA to eIF4E and to ribosomes changes in stress. We were able to calculate this with available poly(A)-enriched total RNA (RNA-seq) and ribosome footprint (Ribo-seq) data (Figure 7A, Supplemental Files 3 and 4, respectively). Changes in ribosomal association of mRNAs, which are not influenced by changes in abundance, are indicated (Figure 7B, Supplemental Table S8) (log_2_FC (TAU/LIT) >1 or <-1 is the enrichment cut-off controlled by *p-adj* < 0.05). mRNAs with greater ribosomal association in TAU than in LIT localize to the right-side of the plot and mRNAs with greater association in LIT than TAU to the left side. We removed all mRNAs except the subset that is associated to eIF4E4 (Figure 7C) and eIF4E3 (Figure 7D) to visualize their specific ribosome associations. In stress, eIF4E4-associated mRNAs showed overall less ribosomal association, suggesting their translation efficiency is decreased even when bound to eIF4E4.

**Figure 7.**
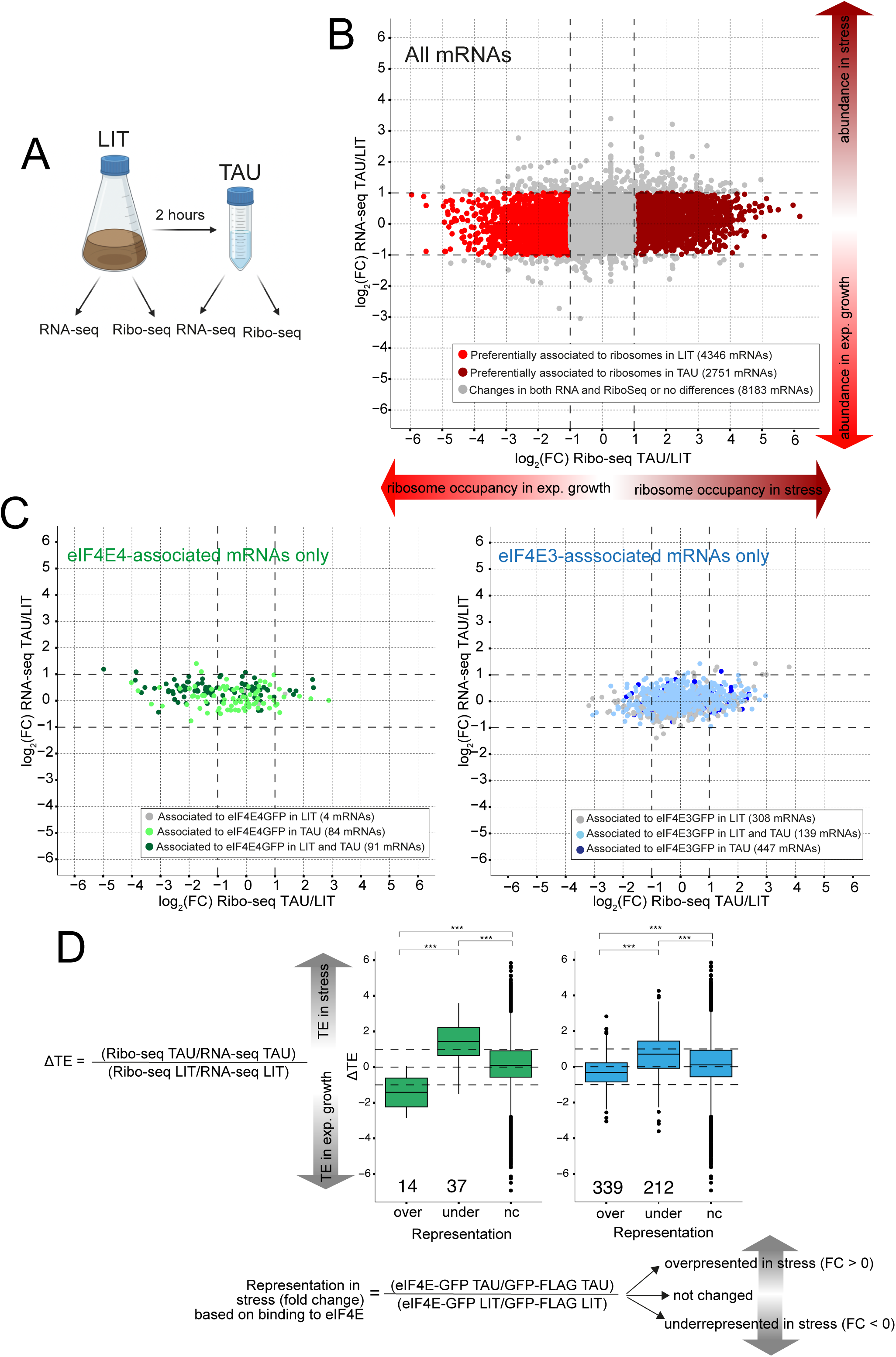
(A) RNA-seq and Ribo-seq were carried out in two different conditions: LIT (epimastigotes in exponential growth and carbon-rich media) and TAU (epimastigotes previously in LIT were incubated in carbon-poor media for two hours). (B) Scatter plot generated by plotting the log_2_ (Fold Change, FC) (DESeq2 output) of the RNA-seq and Ribo-seq experiments comparing TAU and LIT. The cut-offs for the FC values were log_2_ > 1 or < – 1 and are indicated by the black dashed lines. (B) mRNAs which are differentially associated in the immunoprecipitation of eIF4E4-GFP (in LIT, TAU or both) and to eIF4E3-GFP (in LIT, TAU or both) were plotted in the scatterplot of 7B. (D) The variation in translational efficiency (ΔTE) between TAU and LIT is calculated using the formula shown. Then, the variation in representation (association to eIF4E) between TAU and LIT is calculated using the formula. mRNAs are grouped in overrepresented (over, more association to eIF4E in TAU compared to LIT), underrepresented (under, loss of association to eIF4E in TAU compared to LIT) or not changed (nc). A bar plot compares the ΔTE values for each group. A statistical *Wilcoxon*-test comparing the averages between groups confirms that representation in stress anti-correlates to ribosome association.

We also decided to correlate variation in translation efficiency (ΔTE) and association to eIF4E in stress. We first calculated the ΔTE of a mRNA [(Ribo-Seq TAU/RNA-seq TAU)/(Ribo-Seq LIT/RNA-seq LIT)] (the DESeq2 likelihood ratio test (LRT) controlled with a statistic *Wald* test) (Figure 7D). A mRNA with a large ΔTE value has more ribosomal association in TAU; by consequence, a mRNA with a small ΔTE value would have more ribosome association in LIT. We then used a similar equation to calculate the variation in the eIF4E:mRNA association, which we called representation [(E3-GFP TAU/GFP-FLAG TAU)/(E3-GFP LIT/GFP-FLAG LIT)]. A mRNA with representation values > 0 (overrepresented) is more associated to the eIF4E in TAU than LIT; an underrepresented mRNA (values < 0) would be more associated to the eIF4E in LIT than TAU. This resulted in overrepresentation of 339 and underrepresentation of 212 eIF4E3-associated mRNAs. Fewer eIF4E4-associated mRNAs had changes between TAU and LIT: 14 are overrepresented, 11 of which code for ribosomal proteins, and 37 are underrepresented. We plotted the ΔTE values of mRNAs overrepresented, underrepresented and not changed and observed that mRNA representation in the eIF4E immunoprecipitation anti-correlates to ribosome association in stress (*p* < 0.01, *Wilcoxon*-test). In conclusion, eIF4E association is not sufficient for ribosome association: in stress, the more associated to the eIF4E3/4 a mRNA is, the less ribosomal association it has. This implies that mRNAs with translation inhibited by stress would still bind to the eIF4E but downstream steps in translation would be regulated.

## Discussion

Our experiments address functional aspects of two *T. cruzi* eIF4E paralogues and how their association to proteins and mRNA respond to translation arrest during nutritional stress. eIF4E3 and eIF4E4 associate to different proteins and mRNAs. eIF4E3 behaves as a canonical eIF4E and associates to mRNAs encoding proteins from many functional categories. eIF4E4 is specialized to mRNAs codifying ribosomal proteins. In stress, the core eIF4F-like complexes remain stable, suggesting they would still be able to bind mRNA. Indeed, the same mRNAs or mRNAs from the same functional categories, but not stress-responsive mRNAs, associate to eIF4E3 and eIF4E4 and have low ribosome association in stress. We can implicate eIF4E3 to function majorly in optimal growth conditions, but not in stress. The data support a model where eIF4E3 associated mRNAs might not be translated in stress due to a decrease in association with the 43S PIC, which is likely an effect of eIF2 phosphorylation by stress-activated kinases [40]. There is not enough evidence supporting a similar model for eIF4E4, as we did not detect major and reproductible changes in association of 43S PIC proteins with eIF4E4. The best evidence of regulation for eIF4E4 in stress is the PABP1 dephosphorylation. Indeed, phosphorylation is known to affect *T. brucei* eIF4E4:PABP1 interaction [74]. Therefore, stress does not affect the association of eIF4E to mRNAs but rather events downstream of cap mRNA binding, probably either by impeding ribosomes to be recruited or through a more sophisticated effect on how mRNA ends communicate.

The finding that nutritional stress does not dissociate mRNAs from the eIF4E3/4 is counterintuitive, as the eIF4F connects the 43S PIC to mRNAs, but similar findings were reported in yeast [75], [76]. Costello *et al*., 2017 interpreted their results based on the observation that the eIF4F releases the mRNA during scanning phase (or recently shown in elongation [77]): mRNAs with high translational efficiency would cycle between more successive rounds of initiation than mRNAs with low translation efficiency and would have a more dynamic eIF4F association, cycling between 40S joining/AUG scanning and 60S joining/eIF4F release. By consequence, mRNAs with low translation efficiencies would have the eIF4F waiting for the 40S subunit. Their interpretation is that more stable eIF4F:mRNA interactions would be more likely captured in the immunoprecipitation. This model fits our data and possibly explain our results.

We can add to this model how the communication between mRNA ends would be regulated in stress. A preference for the closed loop conformation in non-translating mRNAs can be seen using smFISH. Importantly, this can be associated to ribosome release but not stress granule localization [78]. It is worth mentioning that communication is not limited by eIF4G:PABP only, and not even by poly(A) itself, as poly(A)-tail-less histone mRNAs [79] and m^6^A modified mRNAs [80] have associations bridging the 5’ and 3’ ends. A diversity of protein:protein interactions can communicate mRNA ends: here, in *T. cruzi* we describe for the first time the direct interaction between eIF4E3 and the two poly(A) binding proteins. This interaction is likely mediated by the presence of PAM2 motifs in the amino-terminal extension of eIF4E3 and the MLLE domain in the carboxy-terminal domains of PABP1 and PABP2, which would be the same motifs mediating the eIF4E4 and PABP1 interaction [24], [31].

The PAM2 motif is a short 12-amino acid motif found in a variety of RNA binding proteins [32]. Experimental data from four PAM2-containing proteins was used to propose a model where phosphorylation near the PAM2 motifs has a negative effect on the MLLE interaction. The effect of PABP phosphorylation was not demonstrated, but it was speculated that the phosphorylation at the linker between the last RRM and the MLLE could have a similar effect, in which the phosphate negative charges of the PAM2 and the MLLE would repel themselves [81]. We and others see PABP1 dephosphorylated when global translation is arrested in kinetoplastids [37], suggesting this effect would be conserved in the PAM2:MLLE of eIF4E3/4:PABP1/2. Importantly, *T. brucei* and *Leishmania* PABP1 are phosphorylated at the RRM-MLLE linker [37], [74]. Based on what we discussed so far, we speculate that efficiently translated mRNAs would have a more dynamic eIF4F:mRNA interaction but also more distant 5’ to 3’ ends. A recent study described in detail how the phosphorylation of *T. brucei* eIF4E4 and PABP1 affect their interaction [74]. Their model contradicts what we speculate, as it proposes that mRNAs more efficiently translated would have eIF4E4 and PABP1 phosphorylated. More experimental evidence would be necessary to support one model or the other.

Is there evidence of a similar regulatory mechanism between eIF4E3:PABP1 and eIF4E3:PABP2 *in vivo*? Our data does not provide this, because we could not detect phosphorylation of eIF4E3 and PABP2, but eIF4E3 is a phosphoprotein [69]. We thus first analyzed whether eIF4E3 and PABP1 would associate at all. In our immunoprecipitation, we found a clear preference of eIF4E4 for PABP1, while eIF4E3 could potentially interact with both PABP, which is supported by yeast two hybrid data. Another way to answer this would be to look at the two *Trypanosoma* PABP. Different studies described *Leishmania* and *T. brucei* eIF4E4, but also PABP1, to bind to mRNAs encoding ribosomal proteins [29], [82]. *Leishmania* PABP2 behaves similar to *T. cruzi* eIF4E3 and binds bulk mRNAs [82]. Another important evidence parting eIF4E3 and PABP1 functions is the localization of their mRNAs and proteins associated in starvation stress granules (SG). mRNAs encoding ribosomal proteins are underrepresented from SG [83], [84]. *T. brucei* PABP2, eIF4E3, eIF4G4 and most of PABP2 interaction partners localize to SG, where PABP1, eIF4E4, eIF4A1 and the other PABP1 partners are largely absent [36], [85]. PABP1 can localise to starvation stress granules, but only when ectopically expressed [85], [86]. Taken together, data from several independent studies do fit in a model where two possible regulations for mRNAs happen during stress: eIF4E4/PABP1/associated proteins regulate the translation of ribosomal-protein encoding mRNAs and are prevented from entry to SG during stress. eIF4E3/PABP2/associated proteins regulate the translation of bulk mRNAs and localize to SG upon stress together with other translation factors. It is curious that there is no evidence of ribosomal subunits in trypanosomatid SGs [83]. Ultimately, the *in vivo* eIF4E3:PABP1 interaction is not fully supported.

Ribosomal-protein encoding mRNAs are the mRNAs with more drastic changes with ribosome association between *T. cruzi* replicative epimastigotes and non-replicative metacyclic trypomastigotes [44]. We found that this change can be detected as fast as two hours after TAU stress (manuscript in preparation). In *T. brucei*, a similar finding was reported involving differentiation to a non-dividing stage. Ribosomal-protein encoding mRNAs are stable and abundant but do have a low ribosome association and curiously mRNAs encoding eIF5A, NACA and BTF3 behave similarly [45]. A cautious interpretation is necessary here, as they have very good codon optimality scores [87] and possibly shorter translation elongation times could create artifacts on the TE calculation [18]. Similar findings are described in mammals, where the TE of mRNAs encoding ribosomal proteins is regulated by TOP motifs adjacent to the cap. Upon starvation or mTOR inhibition, drastic changes in the TE of these mRNAs occur. Little is known on how this signal is relayed, possibly by 4E-BP1 [5] and/or LARP1 [88]. This mechanism is unlikely to exist in trypanosomatids because all mRNAs have a SL and ribosomal-protein encoding mRNAs have very short 5’ UTRs lacking TOP motifs [45], [83]. Recently, ZC3H41 and its associated protein (Z41AP) have been shown to co-precipitate ribosomal-protein encoding mRNAs in a manner sensitive to nutritional stress [89]. We do see association of ZC3H41 and Z41AP with eIF4E4 (native and tagged). However, our data do not fully support their findings, because these are still associated in stress. Another missing link is the prevention of ribosomal-protein encoding mRNAs to localize to starvation stress granules [83]. Taken together, this indicates that a novel mechanism, maybe Kinetoplastida-specific, regulates the translational efficiency of these special mRNAs. We speculate that the dephosphorylation of eIF4E4:PABP1, as presumed for PAM2:MLLE [81], would make these mRNAs less efficiently translated. A somehow convergent evolution example would be the negative effect of LARP1 on mammalian TOP mRNAs. LARP1 is able to bind to both the TOP and to PABP by PAM2 motifs and compete with eIF4G [88]. This was proposed to close the mRNA ends in another closed loop structure associated to a lower translational efficiency.

The emergence of PAM2 motifs in eIF4E3 and eIF4E4 unique amino-terminal sequences is intriguing and could explain why these are the two factors with active roles in translation. A phylogenetic analysis of PAM2 motifs could trace their origin in trypanosomes, but this is challenging due to their short size. It is tempting to speculate a *de novo* origin of a single PAM2 which was duplicated in a common eIF4E3/4 ancestor. This motivated us for a proteome-wide search of PAM2 motifs using HMMER (http://hmmer.org). We found all PAM2-containing proteins to be involved in mRNA metabolism (G1-IP1, G5-IP, CE1, 4E-IP2, RBP35) [90], [91], [92] and the PAM2 motif was conserved in all available trypanosomatid sequences (Supplemental Figure 7 and Supplemental Table 9). Curiously, G1-IP1 and G5-IP directly interact with eIF4G1 and eIF4G5 and are part of the eIF4E5 and eIF4E6 complex, respectively [90], [91]. Their PAM2 motifs are putative and the direct G1-IP1:PABP and G5-IP:PABP interaction need to be demonstrated, but we have evidence of a strong association with PABP2 [36], [93]. This conservation again underlies the importance of the communication between the mRNA ends.

In summary, we describe the co-existence of two distinct eIF4F complexes in *T. cruzi*, by showing different association of eIF4E3 and eIF4E4 to proteins and to mRNAs. While eIF4E3 appears to be the canonical eIF4E, eIF4E4 associates in ribosomal protein-encoding mRNAs. Both eIF4E3 and eIF4E4 associate to mRNAs generally important for growth, but not to mRNAs specially translated under nutritional stress. This implies that a third eIF4F complex would perform the translation of stress-responsive mRNAs (Figure 8).

**Figure 8.**
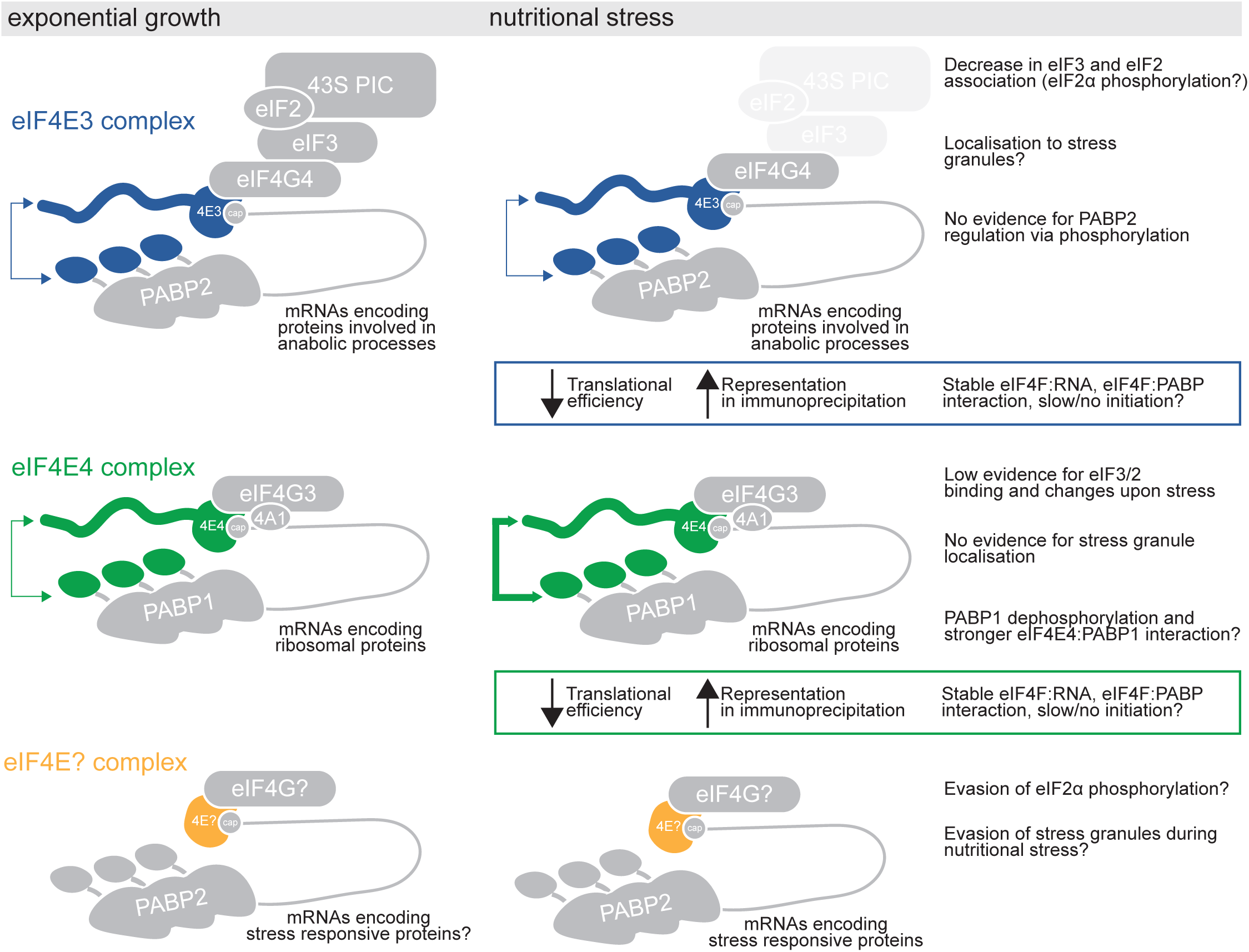
Current model of the co-existence of eIF4E complexes and their regulation during nutritional stress in trypanosomatids. eIF4E3 and eIF4E4 have eIF4F complexes described conserved in three trypanosomatid species. eIF4E3 and eIF4E4 associate to mRNAs important for exponential growth (anabolic functions and ribosomal proteins) and this does not change upon stress when stress-responsive mRNAs are translated. Indeed, representation on the immunoprecipitation in stress anti-correlates to ribosomal association. A possible explanation would be that a stable eIF4E:mRNA/eIF4E:PABP association would lead to slow/no translation re-initiation. A third eIF4E must bind to stress responsive mRNAs and binding could be exclusive to stress or remains in exponential growth. This third eIF4E does not have amino-terminal extensions (exclusive to eIF4E3/eIF4E4) but could bind to PABP2 through other proteins (e.g., G1-IP and G5-IP). It is unlikely that would preferentially bind to PABP1, as it is known to prefer eIF4E4. During nutritional stress, the translation of eIF4E3 and eIF4E4 associated mRNAs are inhibited at several steps of translation. eIF4E3 complexes could be inhibited by a decreased eIF3/eIF2 association, likely an effect of eIF2α phosphorylation, and localisation in stress granules. eIF4E4 could be inhibited by an effect of PABP1 phosphorylation on PAM2:MLLE interaction and therefore in the closed loop structure of their mRNAs and decreased re-initiation rates. The third eIF4E must evade the effect of eIF2α phosphorylation, likely by using upstream open reading frames (uORFs), and the localization in stress granules.

## 6. Data Availability

All proteomics data have been deposited to the ProteomeXchange Consortium through the PRIDE partner repository [94] with the dataset identifier PXD051440. All RNA-seq data have been deposited at NCBI’s Gene Expression Omnibus [95] and are accessible through GEO Series accession number GSE263777.

## 7. Acknowledgements

We express our gratitude to Vanessa Martin Santos and Sibelli Tanjoni for their assistance in *T. cruzi* culture, medium preparation, differentiation, and purification of metacyclic trypomastigotes. Special thanks to Leticia da Silva Pereira for cloning PABP1491-570. We also extend our appreciation to Michel Batista, Kelly Cavalcanti Machado, and Rodrigo Soares Caldeira Brant for their efforts in sample preparation and mass spectrometry analysis. We are indebted to the FIOCRUZ Program of Technological Platforms for granting us access to the Mass Spectrometry (RPT-02H), High Performance Sequencing Platform and Integrated Structural Biology (RPT15A) facilities.

## 9. Supplemental materials

**Supplemental Table 1.** List of primers used in this study.

**Supplemental Table 2.** Full dataset of the immunoprecipitations done in different buffers and RNAse A.

**Supplemental Table 3.** Full dataset of the immunoprecipitations done for eIF4E3 in all strategies and conditions and analysed by mass spectrometry.

**Supplemental Table 4.** Full dataset of the immunoprecipitations done for eIF4E4 in all strategies and conditions and analysed by mass spectrometry.

**Supplemental Table 5.** Full dataset of the comparisons of the proteins associated between eIF4E3 and eIF4E4 shown in Figure 5.

**Supplemental Table 6.** Full dataset of the RNA analysis of the immunoprecipitations done for eIF4E3 and eIF4E4 in the two capture strategies (data in Figure 6 and in Figure S6).

**Supplemental Table 7.** Full dataset of the RNA analysis of the immunoprecipitations done for eIF4E4 for the second time, capturing the native protein using antibodies, but including trypomastigote metacyclics (data in Figure 6 and Figure S8).

**Supplemental Table 8.** Full dataset of the analysis shown in Figure 7. Scripts used to calculate translation efficiency and representation of mRNAs in the immunoprecipitation.

**Supplemental Table 9.** PAM2 sequences, either experimentally validated, manually curated or predicted by the HMMER.

## 10. Figures and legends

**Supplemental Figure 1**. Validation of PABP1 and PABP2 antibodies with recombinant proteins. Validation of PABP1 and PABP2 antibodies with recombinant proteins. (A) Volumes of purified full length PABP1 and PABP2 were run in a gel alongside known BSA concentrations and stained with Coomassie. Bands were quantified and a standard curve used to quantify full length PABP1 and PABP2. (B) Two western blottings were done with PABP1 and PABP2 antibodies. In PABP1 western blotting, 5 times more recombinant PABP2 was added. In PABP2 western blotting, 5 times more recombinant PABP1 was added. Still, no cross-reaction happened.

**Supplemental Figure 2**. All uncropped western blottings presented in this study.

**Supplemental Figure 3**. Conservation of PAM2 motifs “A”, “B” and “C” of the amino-terminal sequences of eIF4E3 in trypanosomatids. (A) Distribution of PAM2 motifs and the position of aminoacids in the amino-terminal of eIF4E3 in *Trypanosoma cruzi*, *Trypanosoma brucei* and *Leishmania major*. (B) Alignment of the amino-terminal sequences of eIF4E3 in trypanosomatids, PAM2 “A”, “B” and “C” are conserved and indicated in blue boxes (Cfa: *Crithidia fasciculata*, Lpy: *Leptomonas pyrrhocoris*, Lse: *Leptomonas seymouri*, Emo: *Endotrypanum monterogeii*, Lbr: *Leishmania braziliensis*, Ldo: *Leishmania donovani*, Ltr: *Leishmania tropica*, Lma: *Leishmania major*, Lme: *Leishmania mexicana*, Lta: *Leishmania tarentolae,* Ocu: *Obscuromonas modryi*, Ade: *Angomonas deanei*, Tgr: *Trypanosoma grayi*, Tth: *Trypanosoma theileri*, Tcr: *Trypanosoma cruzi,* Tbr*: Trypanosoma brucei*, Pha: *Phytomonas sp*. isolate Hart1, Pem: *Phytomonas sp.* isolate EM1). (C) All PAM2 motifs known in the literature curated in [32] or 27 predicted and experimentally validated (See Table S9 for the sequences and references from their studies). (D) PAM2 motifs from Kinetoplastids eIF4E3 amino-terminal sequences were aligned, HHM and logos generated (http://skylign.org).

**Supplemental Figure 4**. (A) Volcano plots generated by plotting the –log_10_ p-value versus the *t*-test difference, comparing the bait (endogenous eIF4E3) to a negative control (IgG coupled beads) in LIT and TAU conditions. The black line is the statistical cut-off from Perseus, but we used *t*-test difference > 2 and –log_10_*p*-value > 2. Selected hits are indicated in blue. (B) Volcano plot generated by plotting the –log_10_ p-value versus the *t*-test difference, comparing the bait (eIF4E3-GFP) to a negative control (GFP-FLAG cells) in LIT and TAU conditions. The black line is the statistical cut-off from Perseus, but we used *t*-test difference 2 and –log_10_*p*-value > 2. Selected hits are indicated in blue. (C) Venn diagram of the enriched proteins between LIT and TAU for the two capture strategies. The predicted protein localization of the majority of the proteins is indicated.

**Supplemental Figure 5**. (A) Volcano plots generated by plotting the –log_10_ p-value versus the *t*-test difference, comparing the bait (endogenous eIF4E4) to a negative control (IgG coupled beads) in LIT and TAU conditions. The black line is the statistical cut-off from Perseus, but we used *t*-test difference > 2 and –log_10_*p*-value > 2. Selected hits are indicated in blue. (B) Volcano plot generated by plotting the –log_10_ p-value versus the *t*-test difference, comparing the bait (eIF4E4-GFP) to a negative control (GFP-FLAG cells) in LIT and TAU conditions. The black line is the statistical cut-off from Perseus, but we used *t*-test difference 2 and –log_10_*p*-value > 2. Selected hits are indicated in green. (D) Venn diagram of the enriched proteins between LIT and TAU for the two capture strategies.

**Supplemental Figure 6**. Venn diagram with all comparisons between mRNAs differentially associated with eIF4E3 and/or eIF4E4 in different capture strategies and conditions. Only mRNAs that met the criteria (log_2_(Fold Change, FC) > 1 and False Discovery Rate (FDR or p-value adjusted) < 0.05) were used for comparisons.

**Supplemental Figure 7**. Violin plots comparing mRNAs differentially associated with (A) eIF4E3/(B) eIF4E4 and mRNAs not associated in abundance (log_2_ of transcripts per million, TPM, using data from RNA-seq in LIT and TAU from Figure 7), ribosomal occupancy (log_2_ of transcripts per million, TPM, using data from Ribo-seq in LIT and TAU from Figure 7), length (log_2_ of gene length) and translational efficiency (direct subtraction of the values log_2_TPM Ribo-seq – log_2_TPM RNAseq represent the equation log_2_ (TPM Ribo-seq/TPM RNA-seq)). All comparisons were done with the data obtained in LIT and TAU. Comparisons between associated/not associated mRNAs were done using a statistical *Wilcoxon*-test and the *p*-value is indicated.

**Supplemental Figure 8**. Venn diagram comparing mRNAs differentially associated with endogenous eIF4E4 in two independent experiments. Only mRNAs that met the criteria (log_2_(Fold Change, FC) > 1 and False Discovery Rate (FDR or p-value adjusted) < 0.05) were used for comparisons.

**Supplemental Figure 9**. Alignment of putative PAM2 *T. cruzi* proteins predicted in this study.

