## Supplementary material for "Two distinct *Trypanosoma* eIF4F complexes co-exist, bind different mRNAs and are regulated during nutritional stress": Figure S1

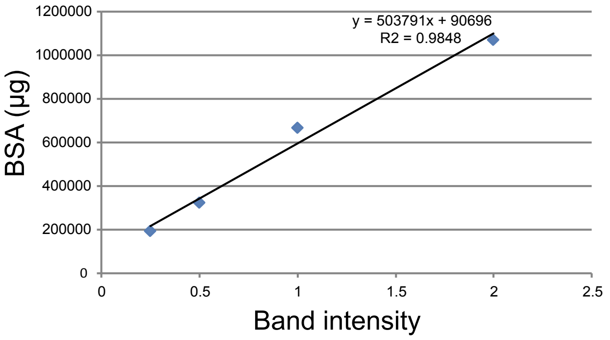

A

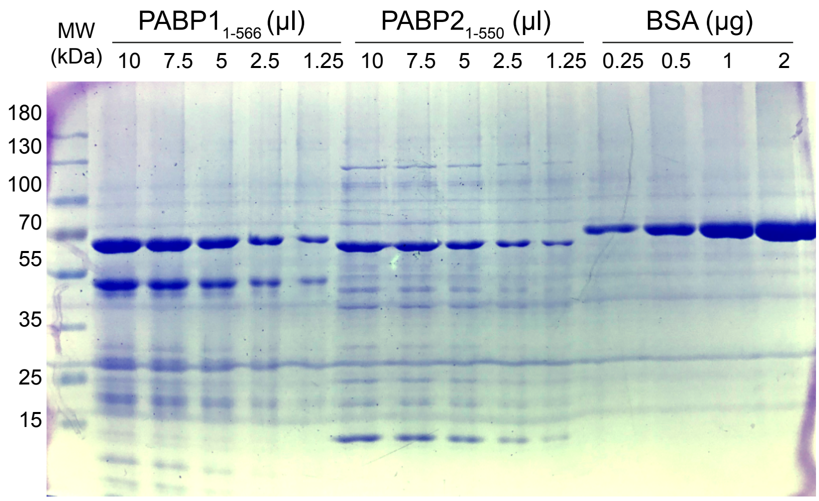

| BSA | 0.25 | 0.5 | 1 | 2 |
|---|---|---|---|---|
| Band intensity | 192000 | 323000 | 667000 | 1070000 |
| | Volume | Intensity | ug/uL | Concentration |
| PABP1 | 10 | 526000 | 0.08641 | |
| | 7.5 | 479000 | 0.10277 | |
| | 5 | 330000 | 0.095 | |
| | | | | ng/uL |
| | | Average | 0.09473 | 94.7251715 |
| | Volume | Intensity | ug/uL | |
| PABP2 | 10 | 389000 | 0.05921 | |
| | 7.5 | 288000 | 0.05222 | |
| | 5 | 150000 | 0.02354 | |
| | | | | ng/uL |
| | | Average | 0.03374 | 33.7433579 |

B

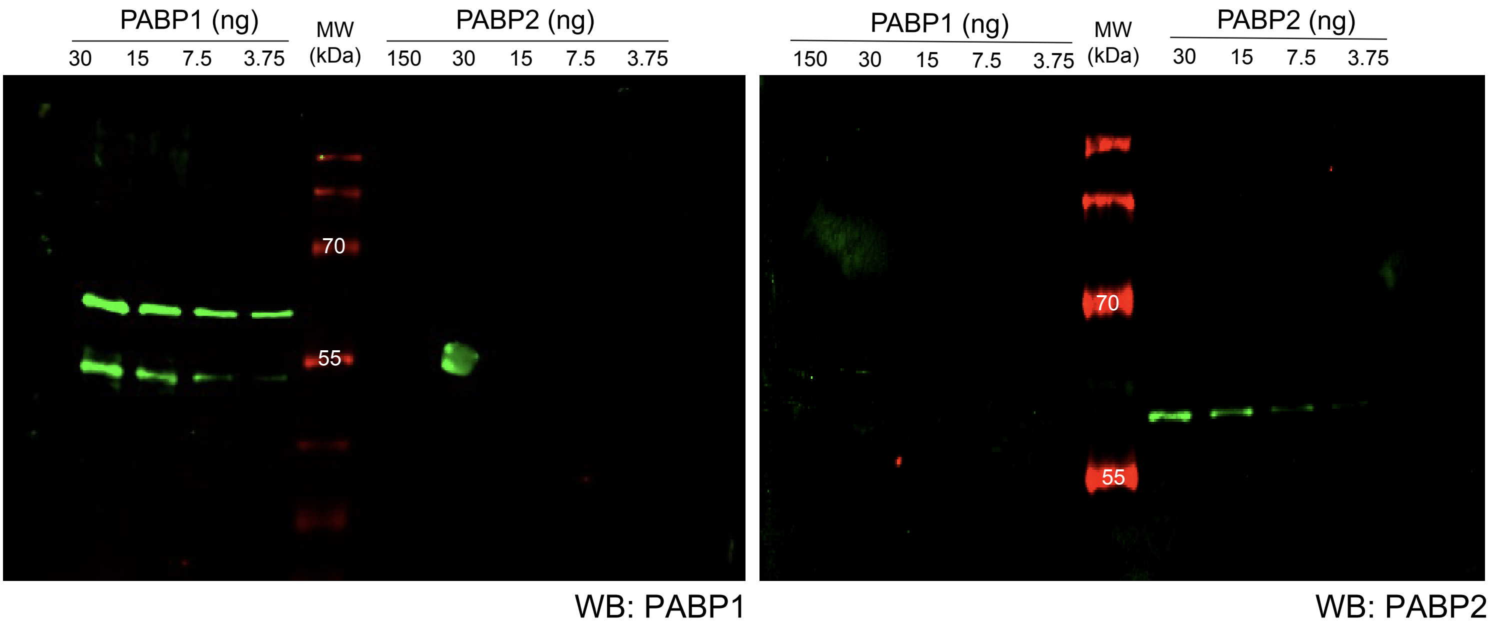
