## Supplementary material for "Two distinct *Trypanosoma* eIF4F complexes co-exist, bind different mRNAs and are regulated during nutritional stress": Figure S2

#### Uncropped western blotting of Figure 1C

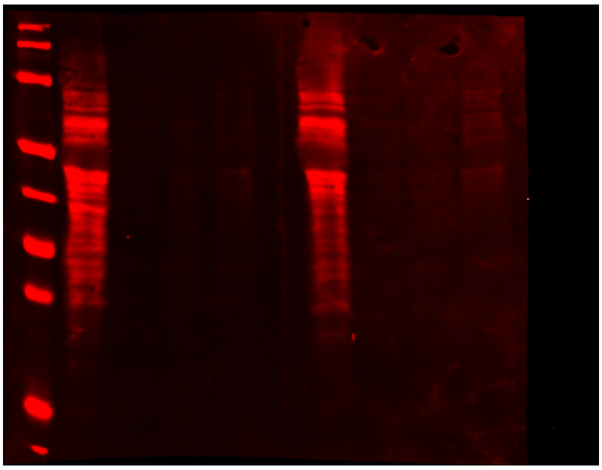


WB: puromycin

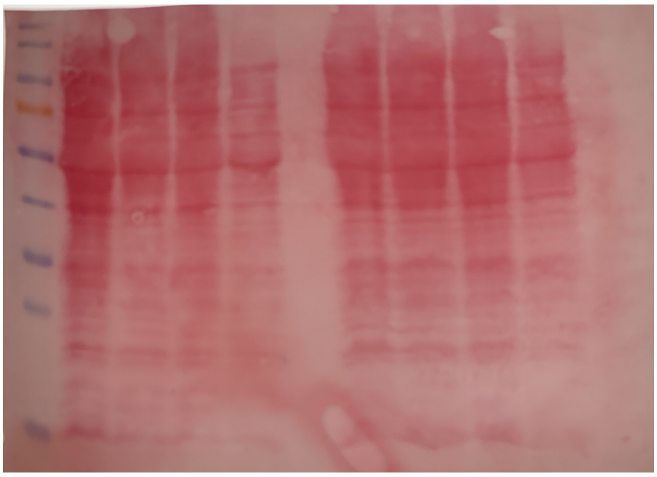

Ponceau

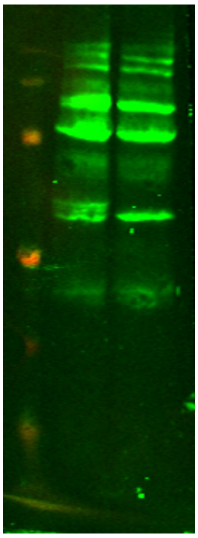


PABP1

Theoretical molecular weight (kDa)
eIF4E3: 49.7
eIF4E4: 45.3
PABP1: 63.8
PABP2: 61.4

### Figure S2

Uncropped western blotting of Figure 1D

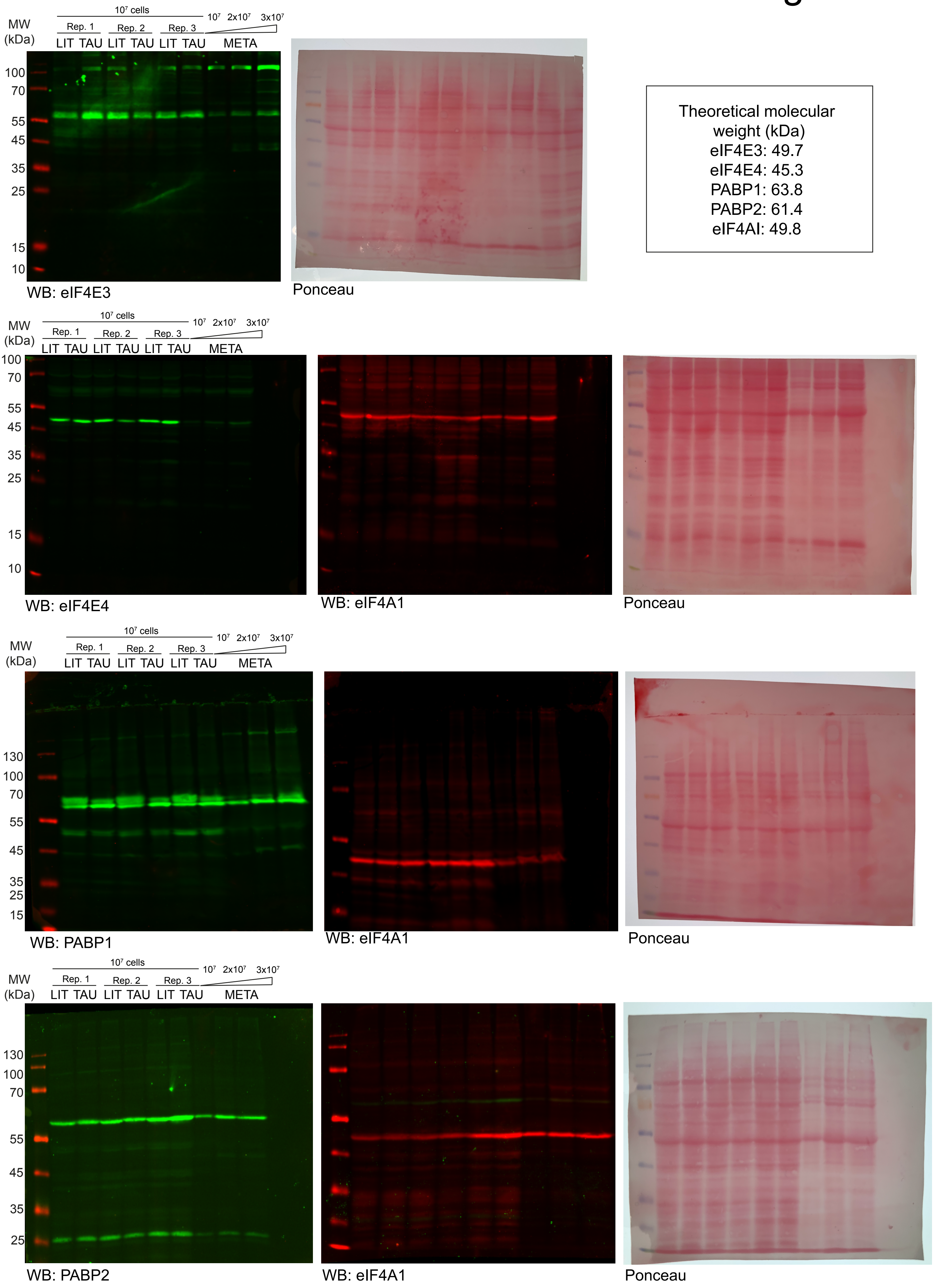
