## Supplementary material for "Two distinct *Trypanosoma* eIF4F complexes co-exist, bind different mRNAs and are regulated during nutritional stress": Figure S3

### A Distribution of PAM2 motifs in eIF4E3 sequences

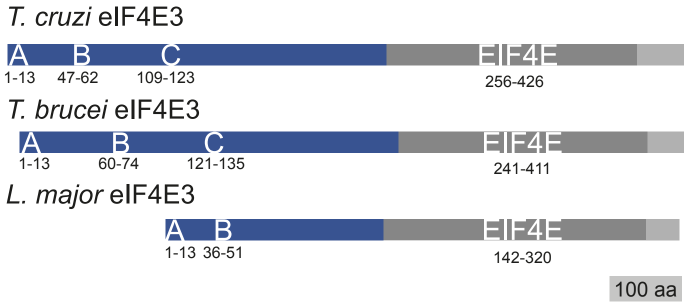


### B Alignment of curated trypanosomatid amino-terminal eIF4E3 sequences and conservation of the PAM2 motifs

```
CfaE3           --MNPSAAAFVPQKGDKT--SSKAKGTATA-----------------TAPHPP-------
LpyE3           --MNPSAAAFVPKKGEKS--PAKGKTNAAA--------------------HPP-------
LseE3           --MNPSAAAFVPKKGDKL--PAKGKASAAA--------------------HPP-------
EmoE3           --MNPSAAAFVPQQRMGKGSPISPPVAAVV---------------PPPSLSPP-------
LbrE3           --MNPSAAAFIPQKSNAKGEPTPSSVAAVT--------------------KPP-------
LdoE3           --MNPSAAAYIPQKSDAKGEPKSSSAAAVV--------------------KPP-------
LtrE3           --MNPSAAAYIPQKSDAKGDPKSSSATAVA--------------------KPP-------
LmaE3           --MNPSAAAYIPQQSDAKGDPKSSSAAAVA--------------------KPP-------
LmeE3           --MNPSAAAYIPPKSDAKGEPKSSSAAAVA--------------------KPP-------
LtaE3           --MNPSAAAYIPQKSDAQREPEPSSVAAVT--------------------KPP-------
OcuE3           MSMNPSAMAFVPHTKRSVKEP----------------------------PQPP-------
AdeE3           --MDPSARNFVPHPSHHHHQH-QQPVAPLG--------------------APP-------
TgrE3           --MNPAAQEFVPRKKRTADPSAVHPANRGP--LPP----------PPPPPPPPQQQQQPP
TthE3           --MNPAAQEFIPQKKRSLDPA-VAPADSRAGQQPPLLHHHHHHQSPPPPPPPP-------
TcrE3           --MNPAAQEFVPRKKCAMDSNAIHMMG----RNPP----QQQQQKQPPQISPP-------
TbrE3           --MNPEAEEFVPKGNRTPGSGGRRGNGGGS-RQIPV---RGERQQPPPPPPPPPQQQQQQV
PhaE3           --MNPAAQEFVLNRTQPNMNY-KTPGNDRT--------------------HTP-------
Pem3            --MNPAAQEFVVNRTQPNMNY-KTPVSDRT--------------------HAP-------
                  *:*  *   ::                                       .*
```

**PAM2 A**

```
CfaE3           SPQLSTKLSAAAQPFVPGA-----------------PKNDAPTPV-------SAAADP--
LpyE3           SPPLATKLSAAAQPFVPVA-----------------PKTDGP------------------
LseE3           SPQLATKLSAAAQPFVPGA-----------------PKTDGPA---------AANADP--
EmoE3           SSQLATRLSATAEPFVPGG-----------------PLQKLEA---------SAHADP--
LbrE3           SAQLVTKLSAAAQPFVPGG-----------------PKQMSAT---------PMVADT--
LdoE3           STQLVTKLSAAAEPFVPGG-----------------PKQASAT---------STHVDP--
LtrE3           STQLATKLSAAAEPFVPGG-----------------PKQMPAT---------STHVDP--
LmaE3           STQPATKLSAAAEPFVPGG-----------------PKQMSAT---------STHVDP--
LmeE3           SIQLATKLSAAAEPFVPGG-----------------PKQMPAT---------STHVDP--
LtaE3           STQLATKLSAAAEPFVPGG-----------------PRHKPAT---------STYVDP--
OcuE3           APVASTKLNATAMPFVPGVSGG-----PGPAGFAAPPPGYAAP---------AQHSDP--
AdeE3           ----GSKLSAQAPPFVPGGQY-------------------------------GNYADP--
TgrE3           LQQQQTKLRASAPAFVPKASTVSAAP-PPPPPLLTQPTAVAASSV-PAPPQAALHVDP-G
TthE3           -QPQPTKLRASAPAFVPKSSAMMAGPLPLDAPTTTTTTTMISSSA-SAVLPAPQPVEPAA
TcrE3           -PATTTKLRATAPAFVPKGSLFPMAT-PPPTSFLQPPVSIPPPAL--APPTVPQYADPNT
TbrE3           APSSQTTLRASAEPFQPKGSQGFGLV-GQVLPYVAGPPGGVPQAW-----ESVEPVPP-L
PhaE3           -INQPTKLSADATPFIPHV-----------------SNKSASNLTIPNADSKLQ-IDPCS
Pem3            -GNQPTKLSADATPFVPHA-----------------PNKPASNQAIAKGDAKPQHVDPSP
                      : * * * .* *
```

**PAM2 B**

```
CfaE3           -------------KVAVDESKAEVPPME-LPQHTSAPPAAA-------------------
LpyE3           --------------VPAEELKPEVRPME-VPKQATAKTAAA-------------------
LseE3           -------------KVPVEEPKPEVPPMK-LPAEATAKDATA-------------------
EmoE3           -------------VVPTEGINTTTHPTTDVLASSLPDAAAA-------------------
LbrE3           -------------KVVTEDGKTTVP----LPTDSLPSSAAA-------------------
LdoE3           -------------KSTTEDGKTTAPPLMERPASSLPDSAAA-------------------
LtrE3           -------------KAATEDGKTTAPSLMERPASSLPDSAAA-------------------
LmaE3           -------------KATTEDEKTTAPLLMECPASSLPDSAAA-------------------
LmeE3           -------------KTATEDGKTTVSPLTERPASSLPNSAAA-------------------
LtaE3           -------------KVTTEDKQSTAPPLMEHPTSSLPDSSAA-------------------
OcuE3           -------------VDEAFGRASGYAPGEGAPAETDA------------------------
AdeE3           -----------QQDSPFSKGATPYTPTNPKLASKFP------------------------
TgrE3           AFGGSDSQPSPGSKTLSLNAVPYTPTNPKIAAKFPSTPQVQRNAS---------------
TthE3           AVPGSDGRPSPGSKALSLDARPYTPTNPKVAAKFPATPQAQQSVAAVDPAEASPTP----
TcrE3           AV----GSGLPGPKSFSPSAPPYTPANPKVVAKFSSPPKTSGSPNP-DPKKTTR------
TbrE3           SVWKSDSVQQAGAKPLSLSATPYTPTNPKISSKFPATPEAQRGPST-EFKDTTP------
PhaE3           SSATDTPTHKNDPNERSESLPQLSVAEPSSSEASPSSVVFNKSGSQ-SFNQISSTEIPPA
Pem3            SPTTDMPAQKNDSVEPSEALPQLSMAESSSNETSPSSPANPKSTSQ-PNNPITSTEVSVS
                                                  .
```

**PAM2 C**

```
CfaE3           ------------DAREKSAEEVAE-EEDS-------------------------------
LpyE3           ------------ESPEGPAKETKDTNDDS-------------------------------
LseE3           ------------ESHEKPTEDAKDTNEDS-------------------------------
EmoE3           -----------------DAGNEADKSEDS-------------------------------
LbrE3           ---------------AGAAKKEVDENEDS-------------------------------
LdoE3           ---------------AGAAKKETDENDDS-------------------------------
LtrE3           ---------------AGAAKKEADENDDS-------------------------------
LmaE3           ---------------AGAAKKEADENDDS-------------------------------
LmeE3           ---------------AGAAKREADENEDS-------------------------------
LtaE3           ---------------AGTANKETDENEDS-------------------------------
OcuE3           ---------------AAASGTSASAERSI-------------------------------
AdeE3           ---------------ATPTEQQQEEQEED-------------------------------
TgrE3           TPDAQSAAQQQQQPPPPQQQQQEHSEESHKSQPL-------------GLANRADSVPLEG
TthE3           SPTATAAQQQDQQQQKEQEKQAEEKEQGAKQEPARDPPLQAEKQE--EPQERGAEPAGEE
TcrE3           TPSPHPEQQQEEEQPNEEKQPREEQEPKPDQSPL------AQKAQTLGVSNRIDSARQED
TbrE3           ---------------ASLQPQEKEVVEVGEKAVAI------------------ASPPEED
```

### C Logos of hidden Markov models for PAM2 sequences

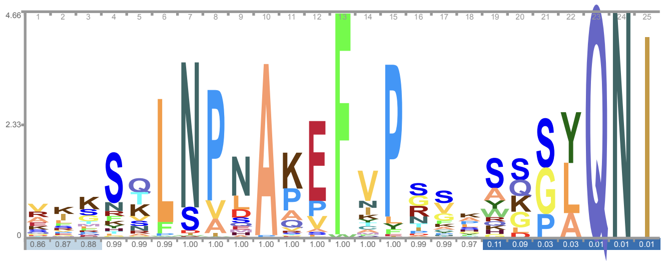

163 manually curated PAM2 sequences were taken from Albretch and Lengauer BBA 2004 and other 27 experimentally validated PAM2 sequences were curated from the literature. These were aligned, a hidden Markov model made and a logo generated.

### D Logos of hidden Markov models for the PAM2 sequences of curated eIF4E3 sequences

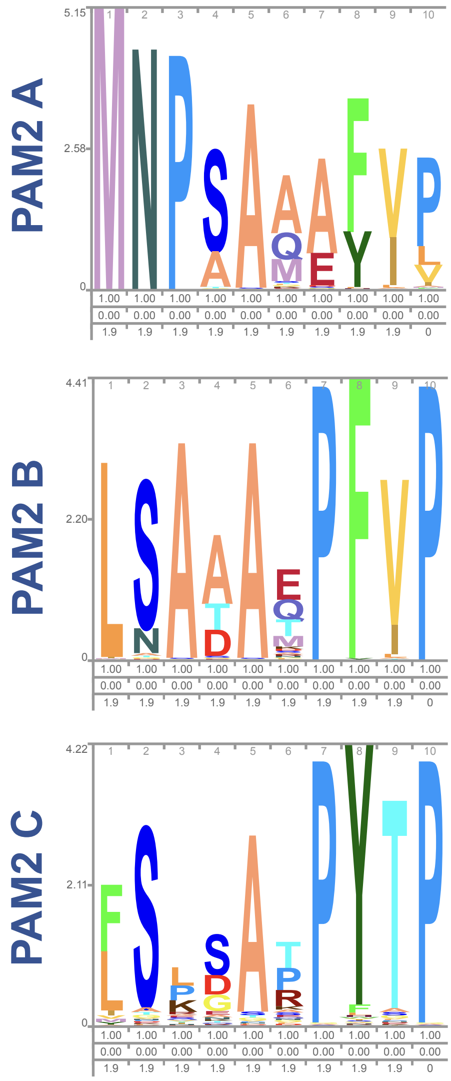


Logos made with http://skylign.org
