## Supplementary figures and images for "Two distinct *Trypanosoma* eIF4F complexes co-exist, bind different mRNAs and are regulated during nutritional stress"

### barplotNull.png

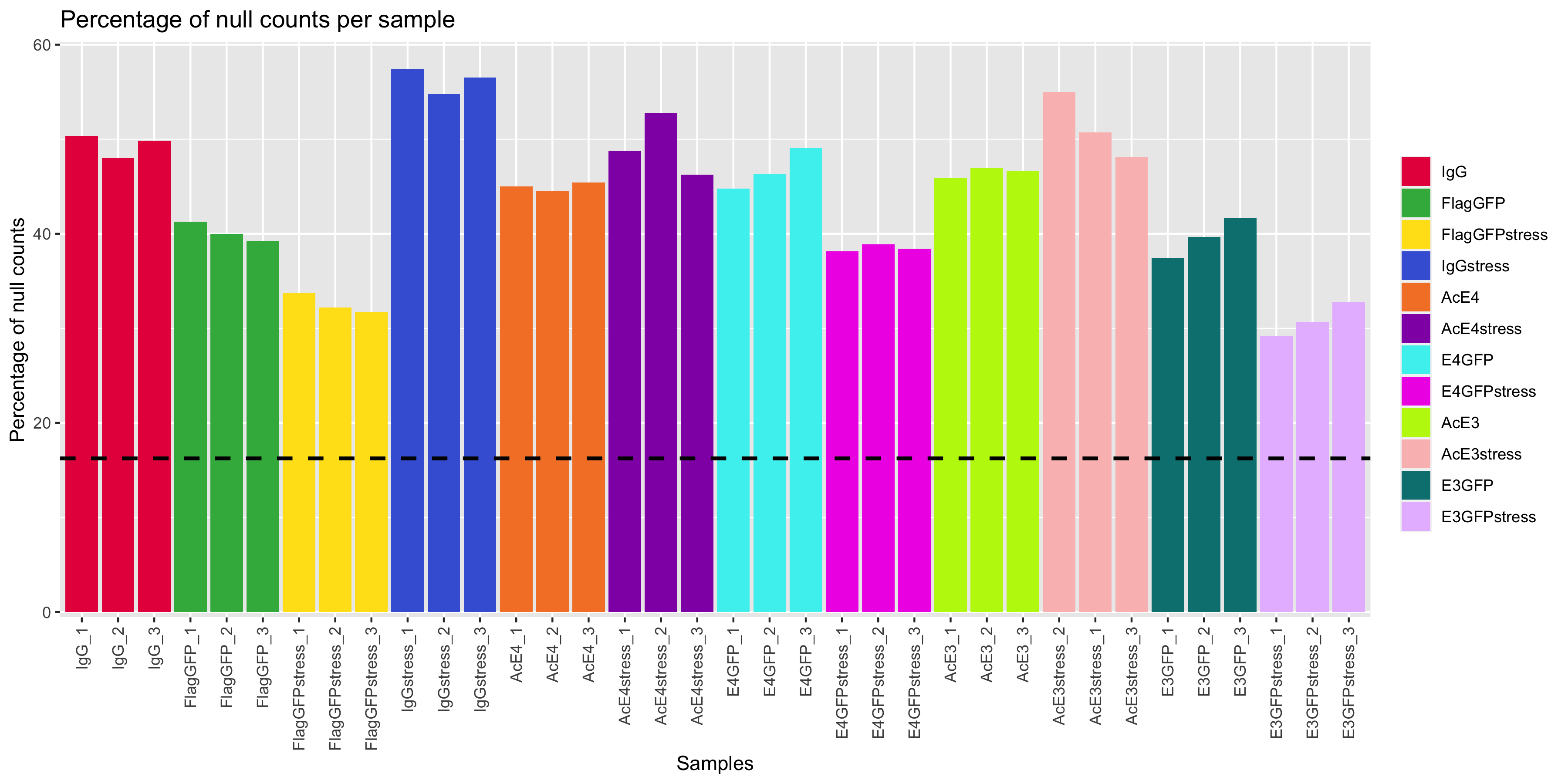

### barplotNull.png

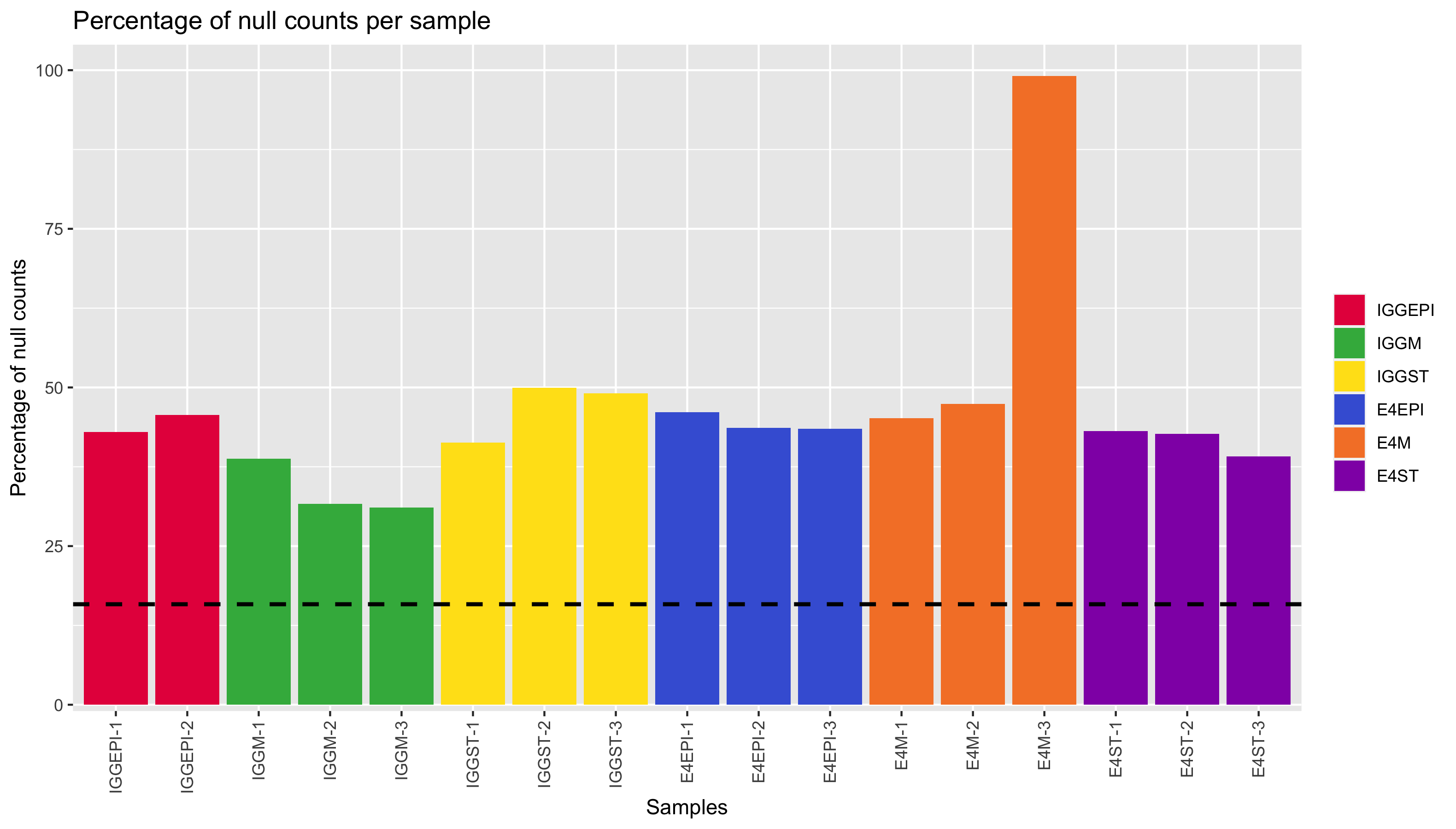

### barplotNull.png

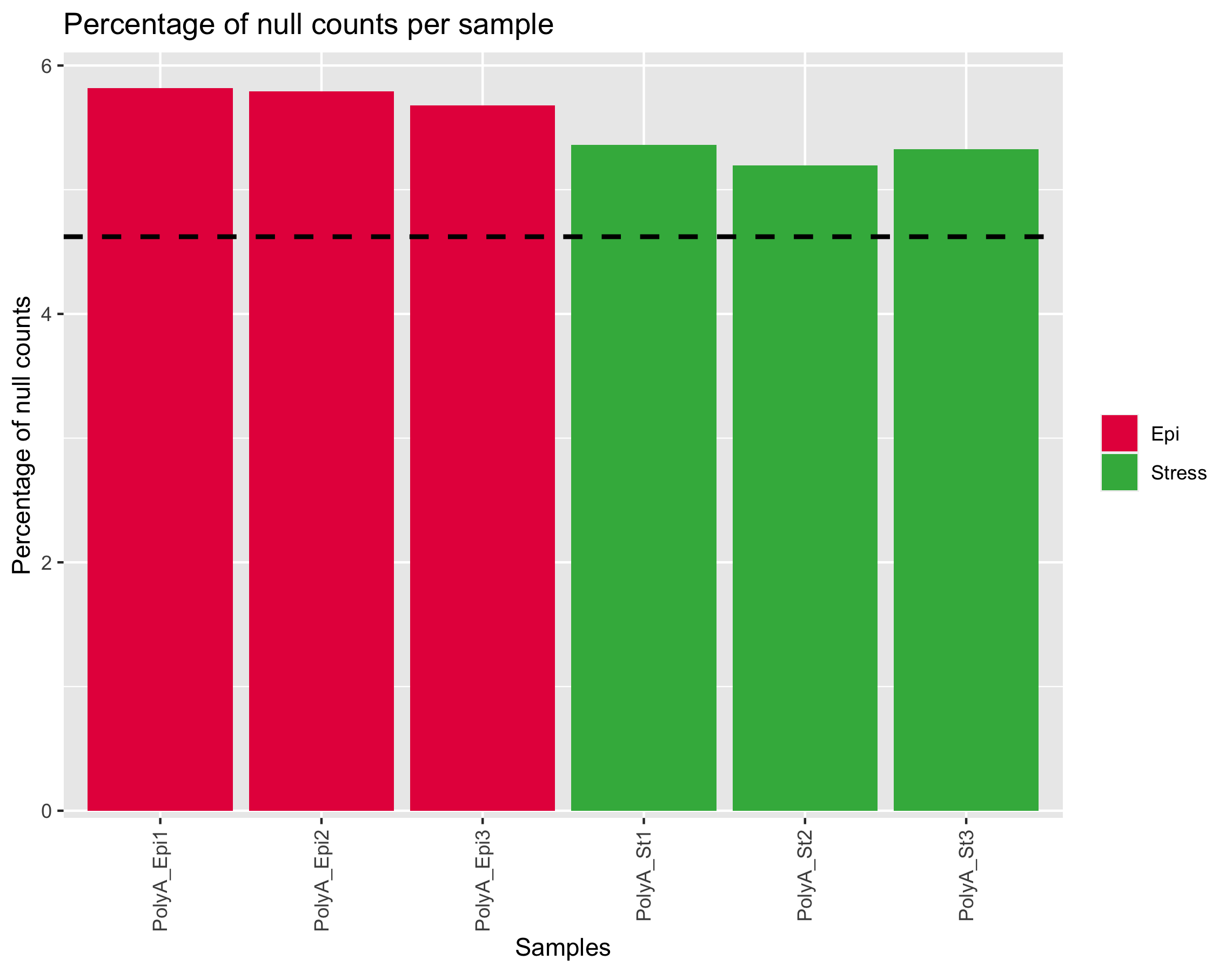

### barplotNull.png

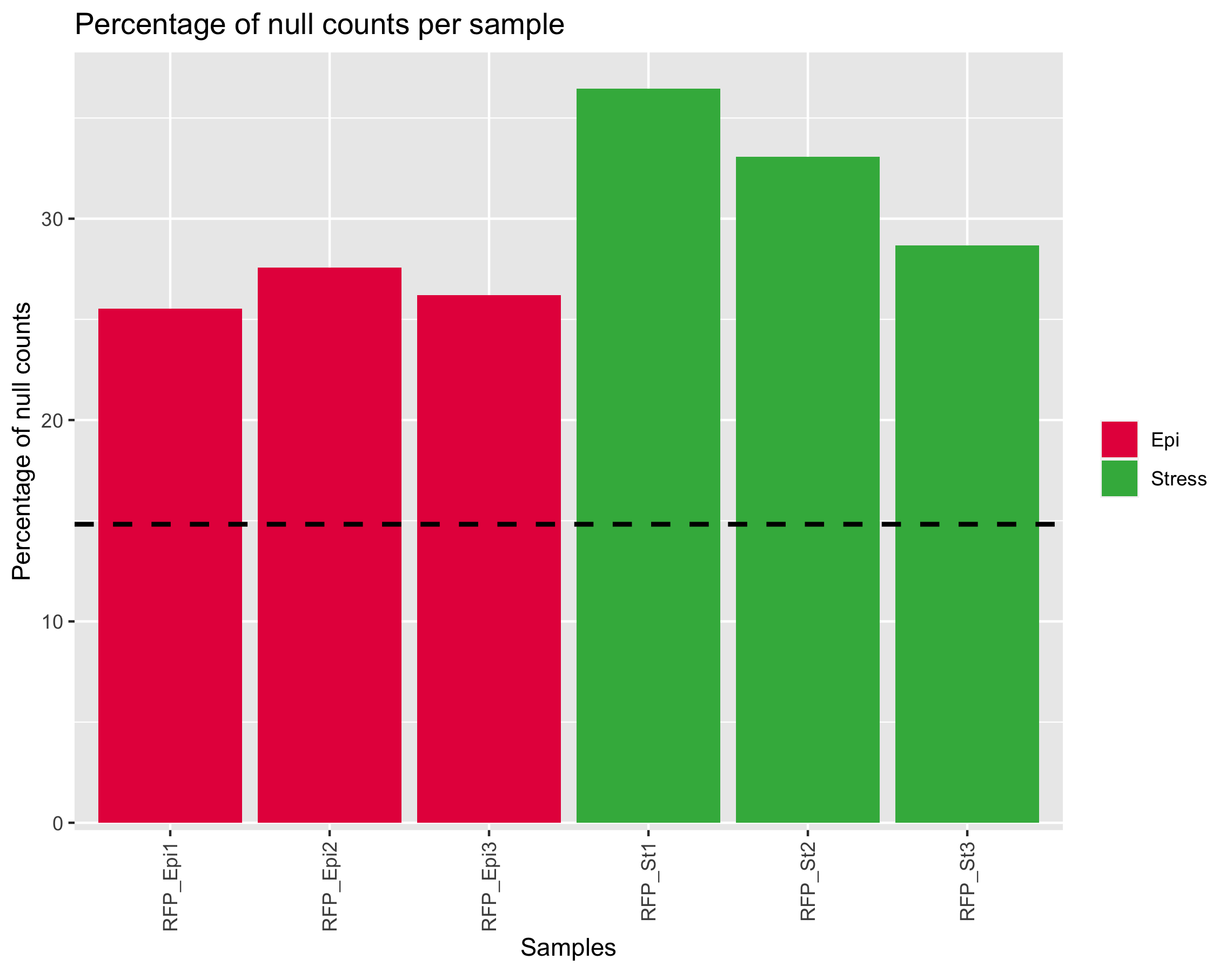

### barplotTotal.png

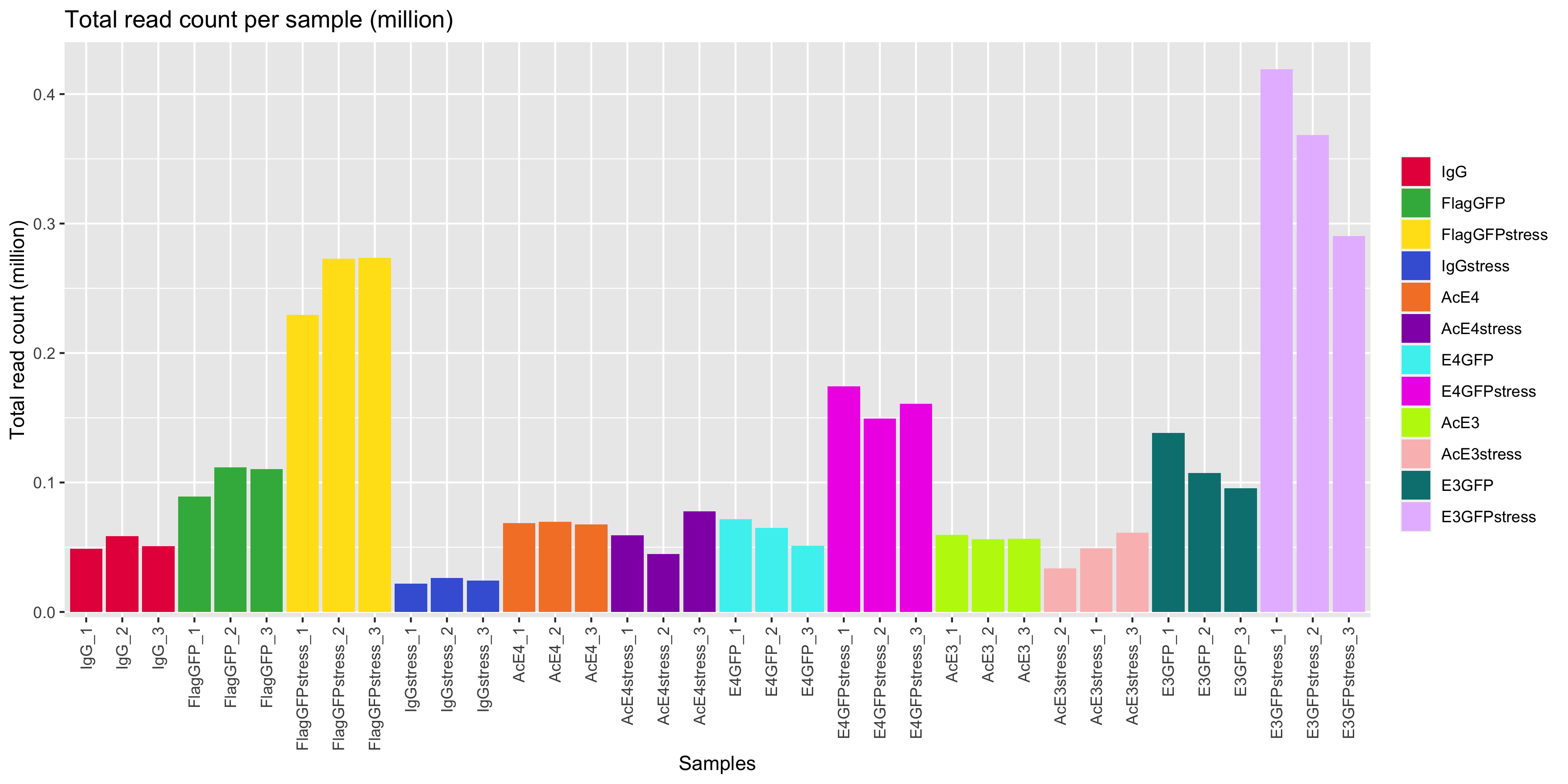

### barplotTotal.png

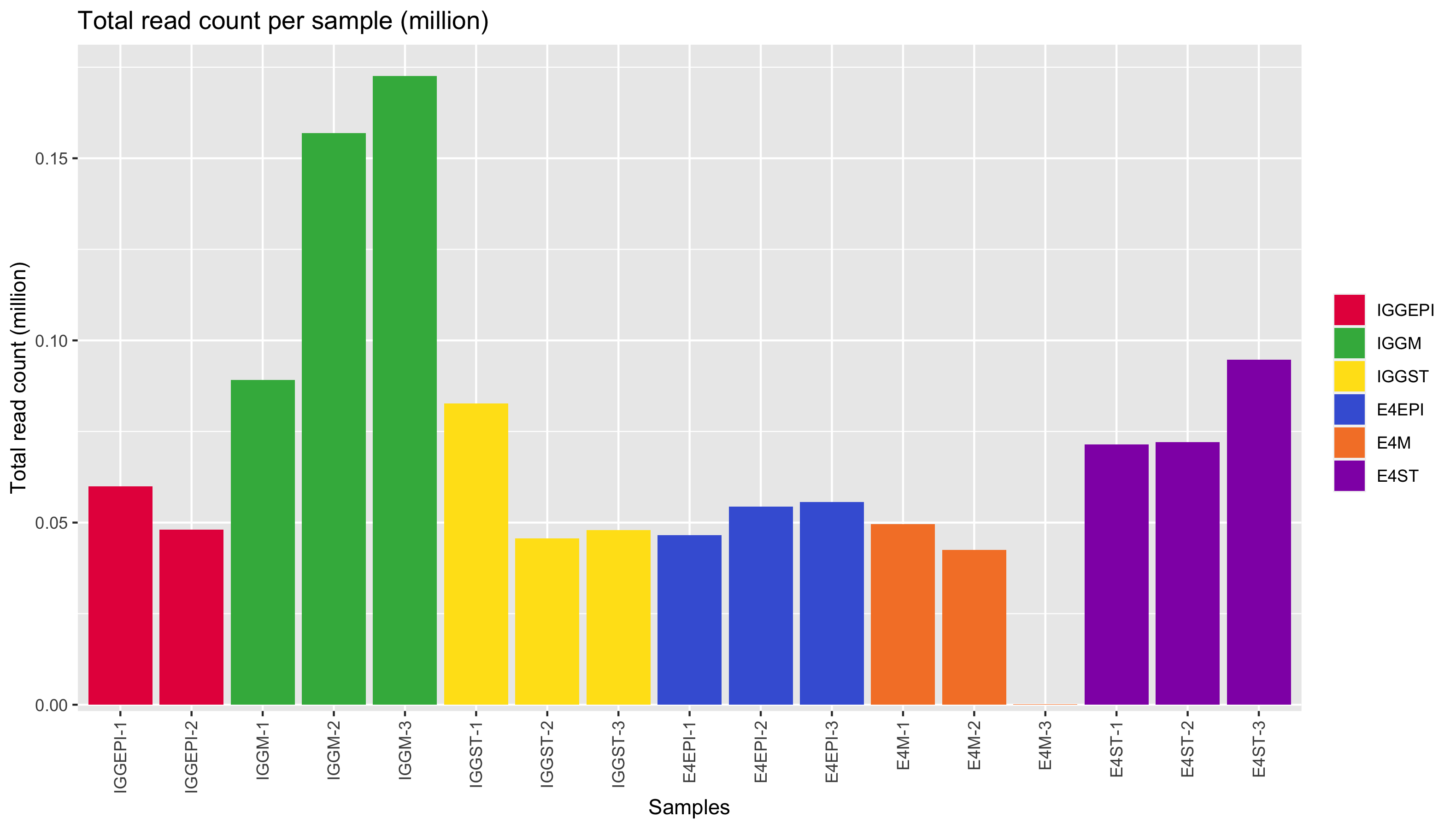

### barplotTotal.png

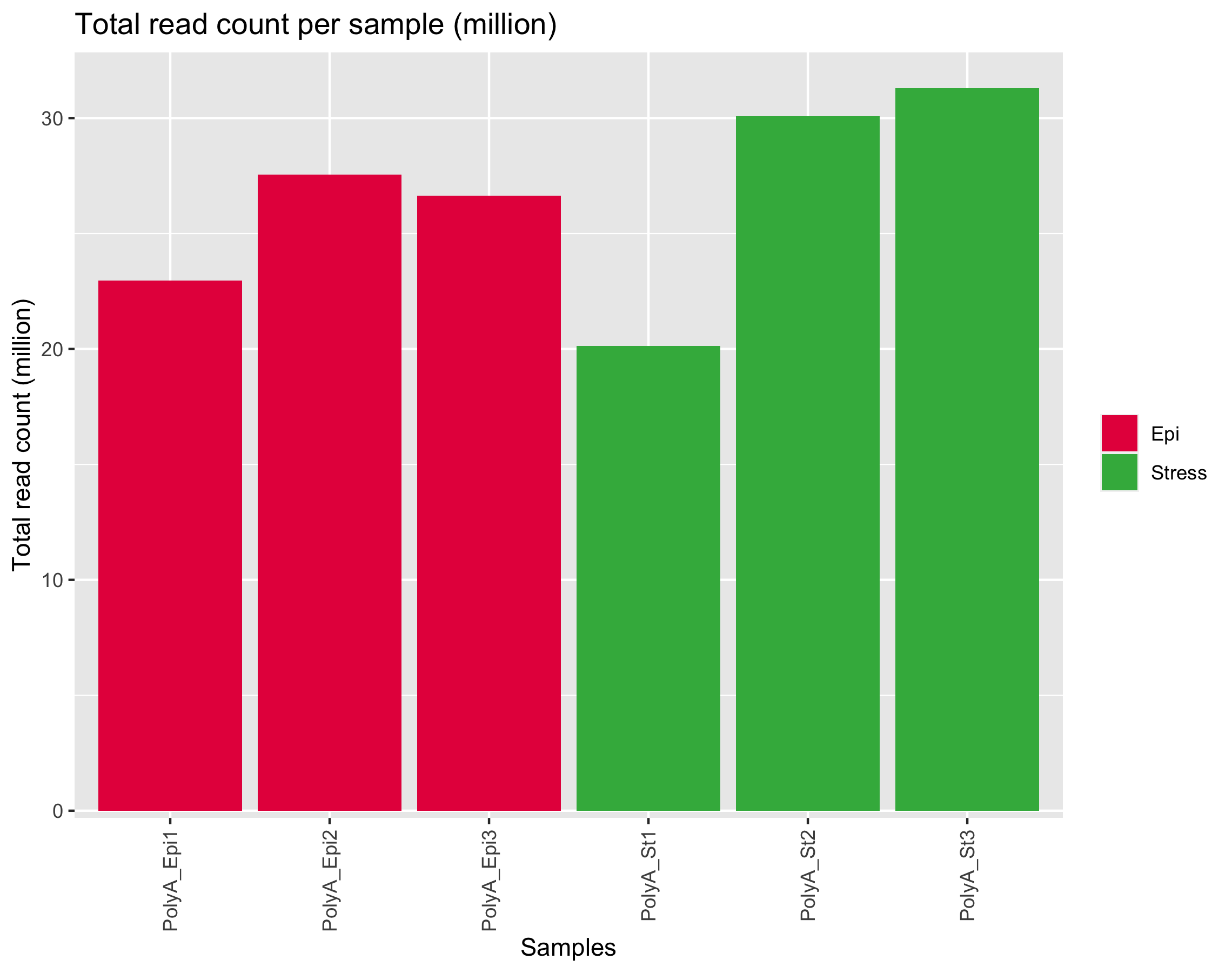

### barplotTotal.png

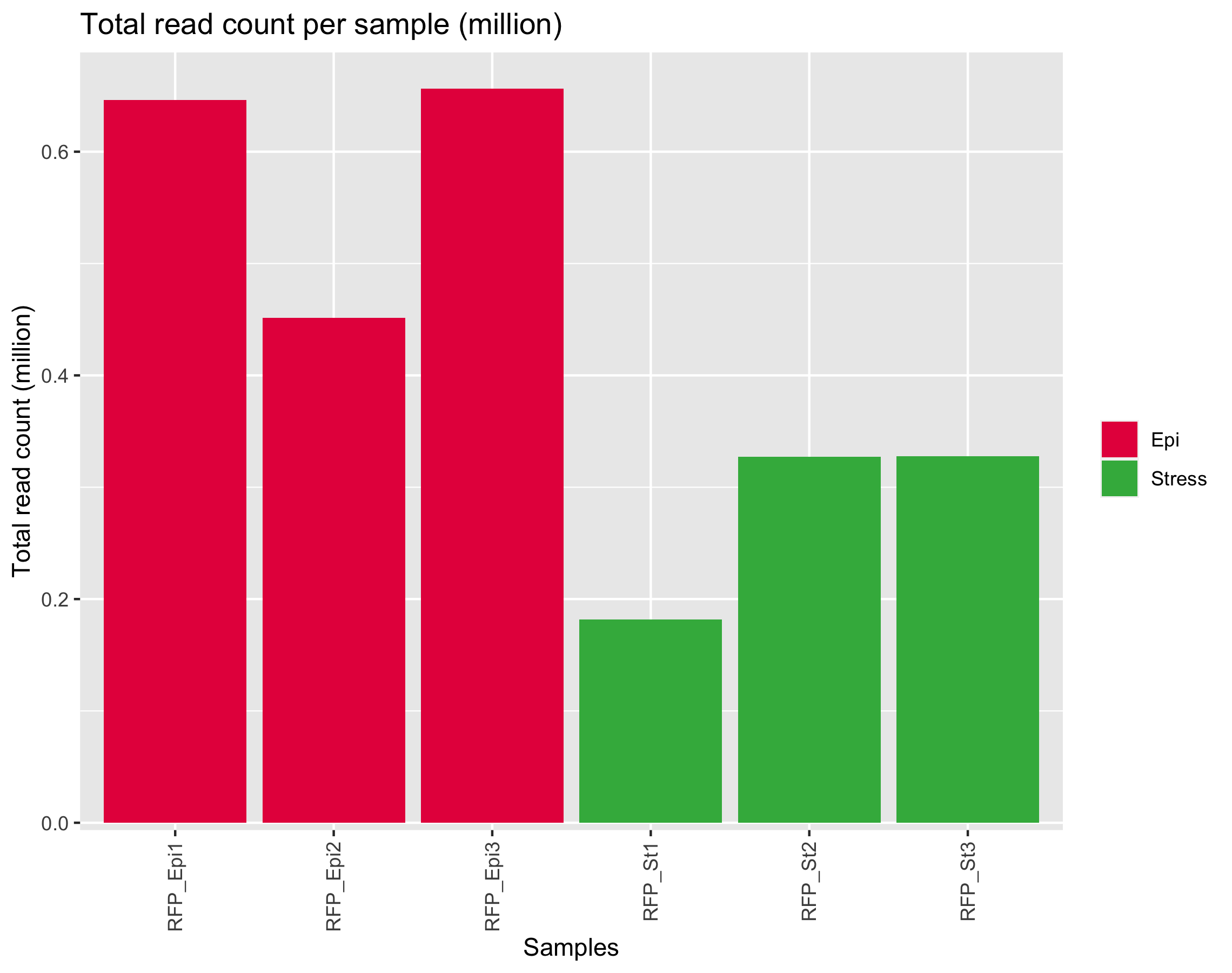

### cluster.png

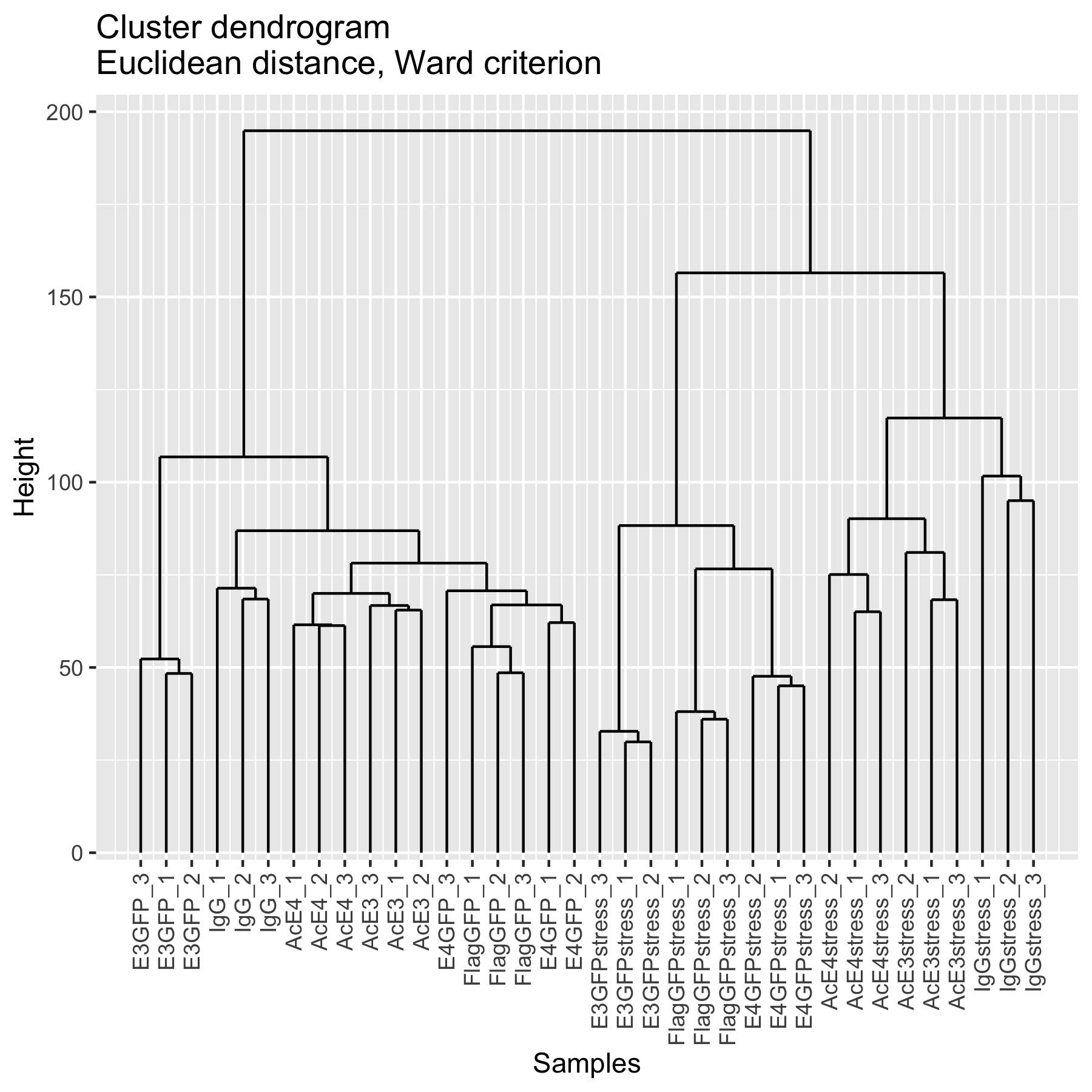

### cluster.png

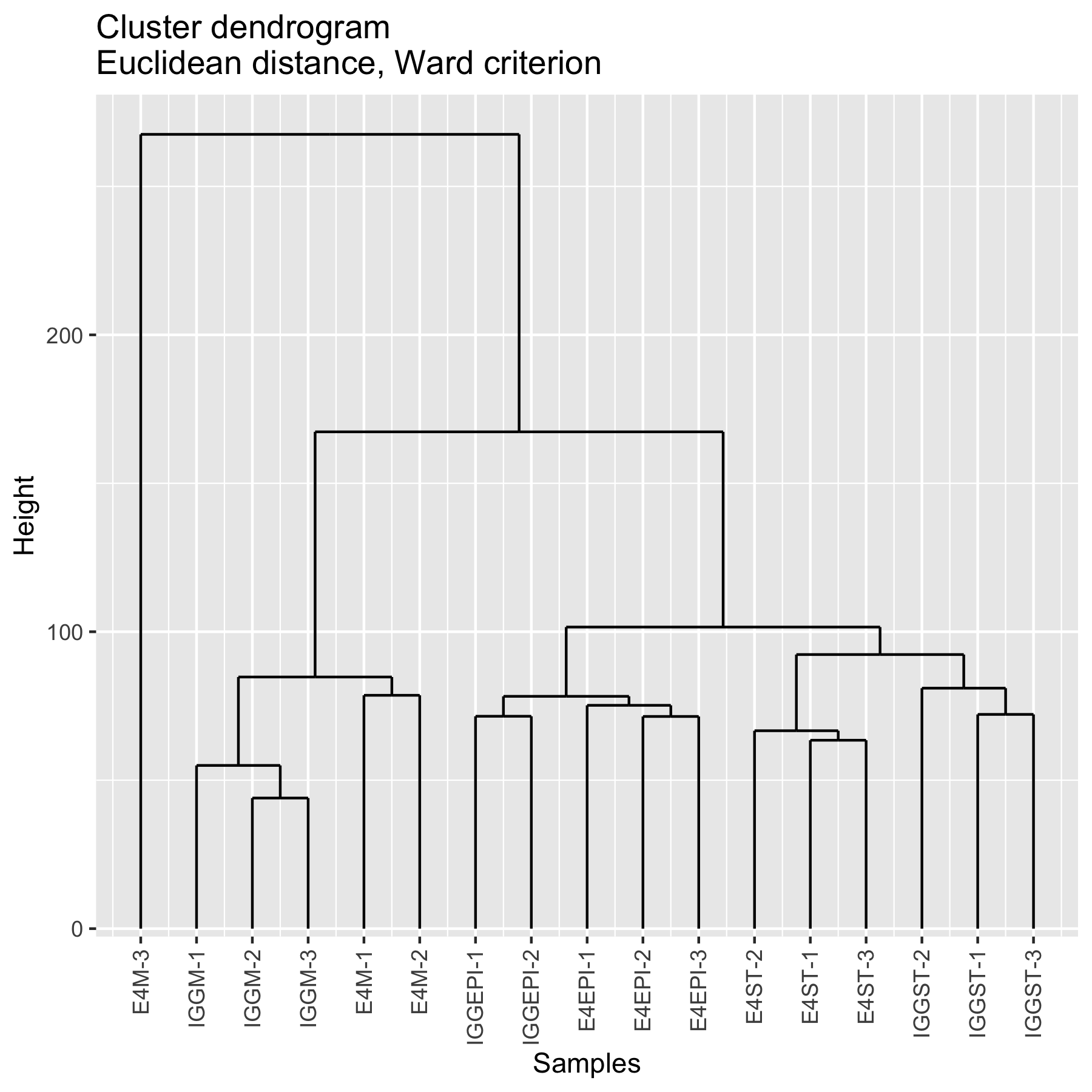

### cluster.png

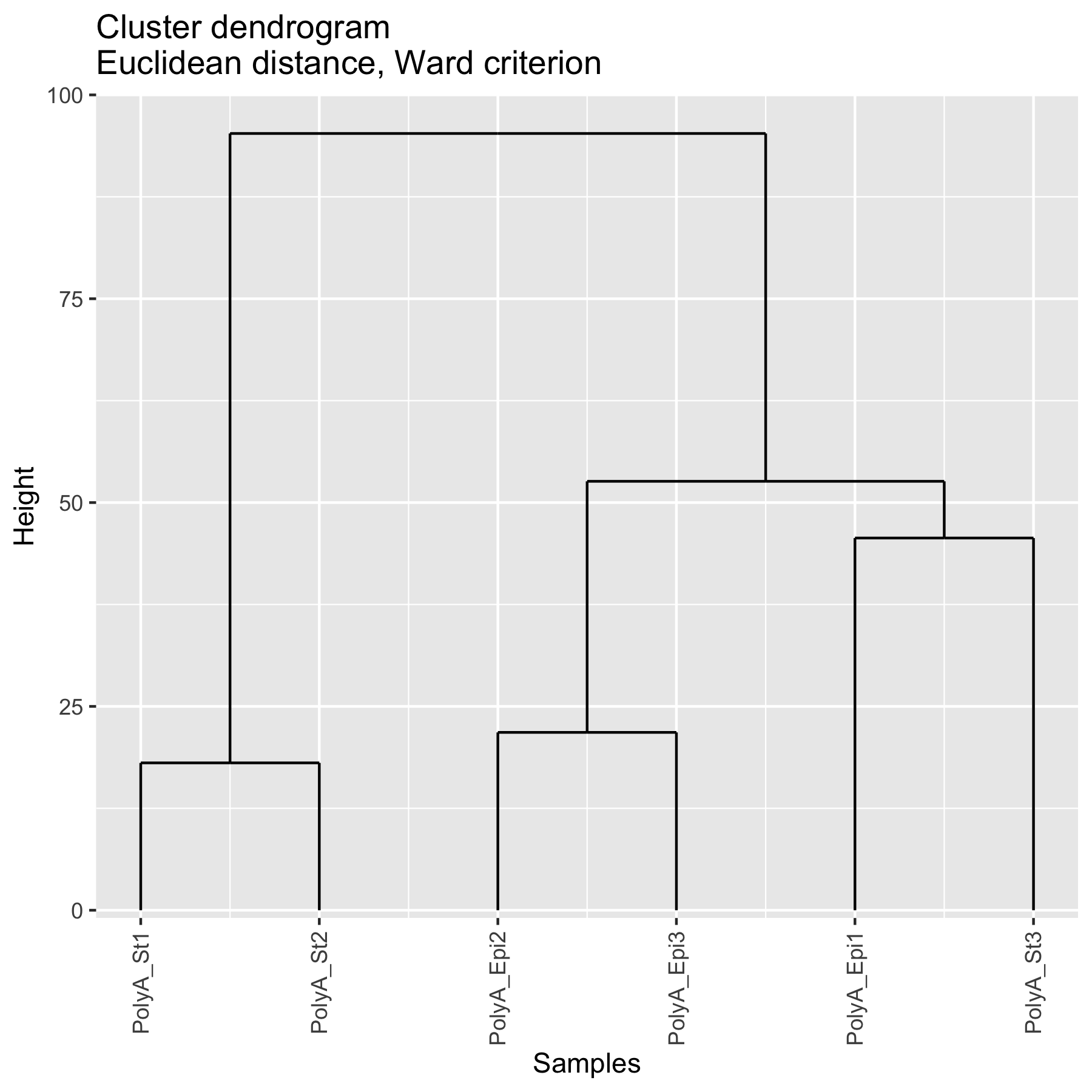

### cluster.png

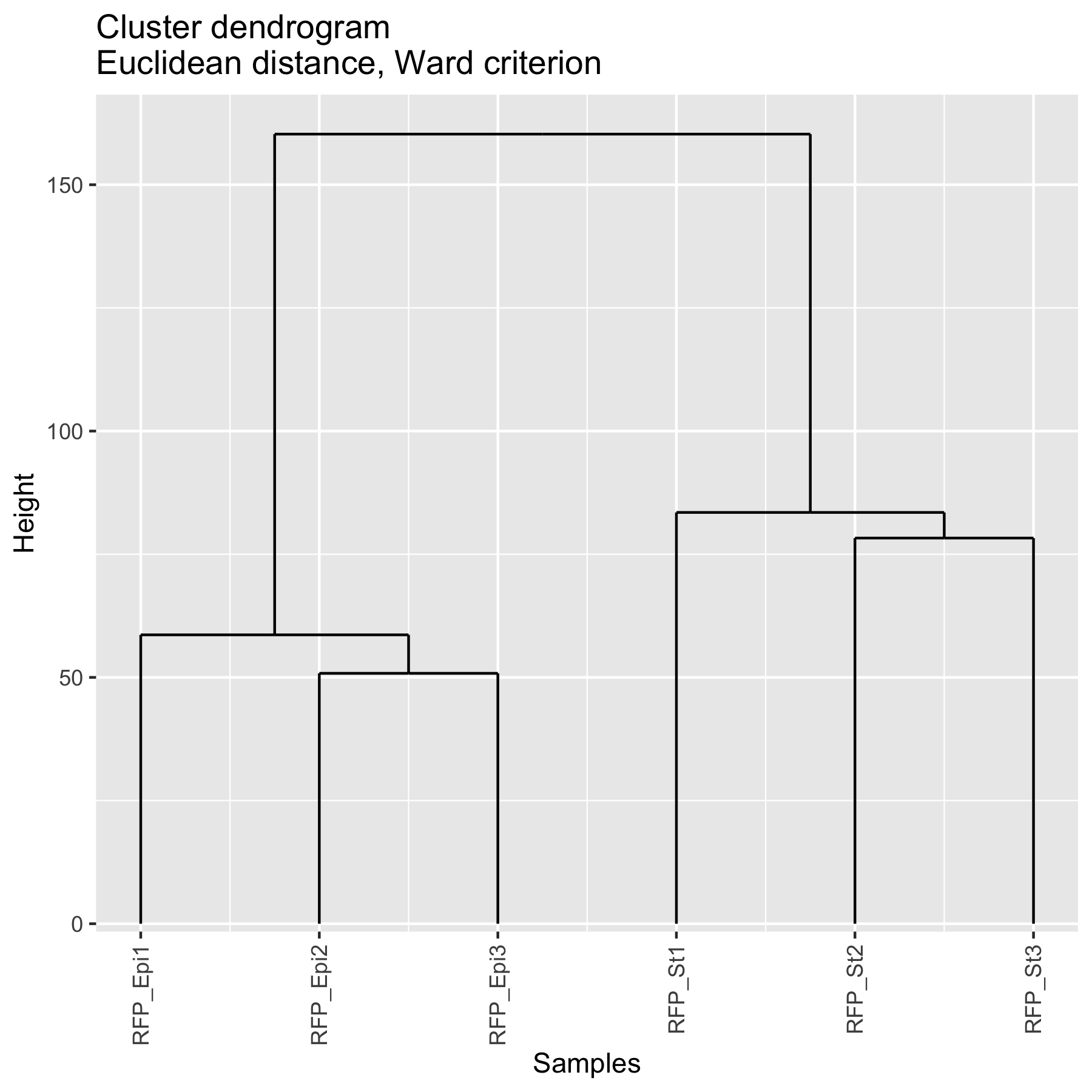

### countsBoxplots.png

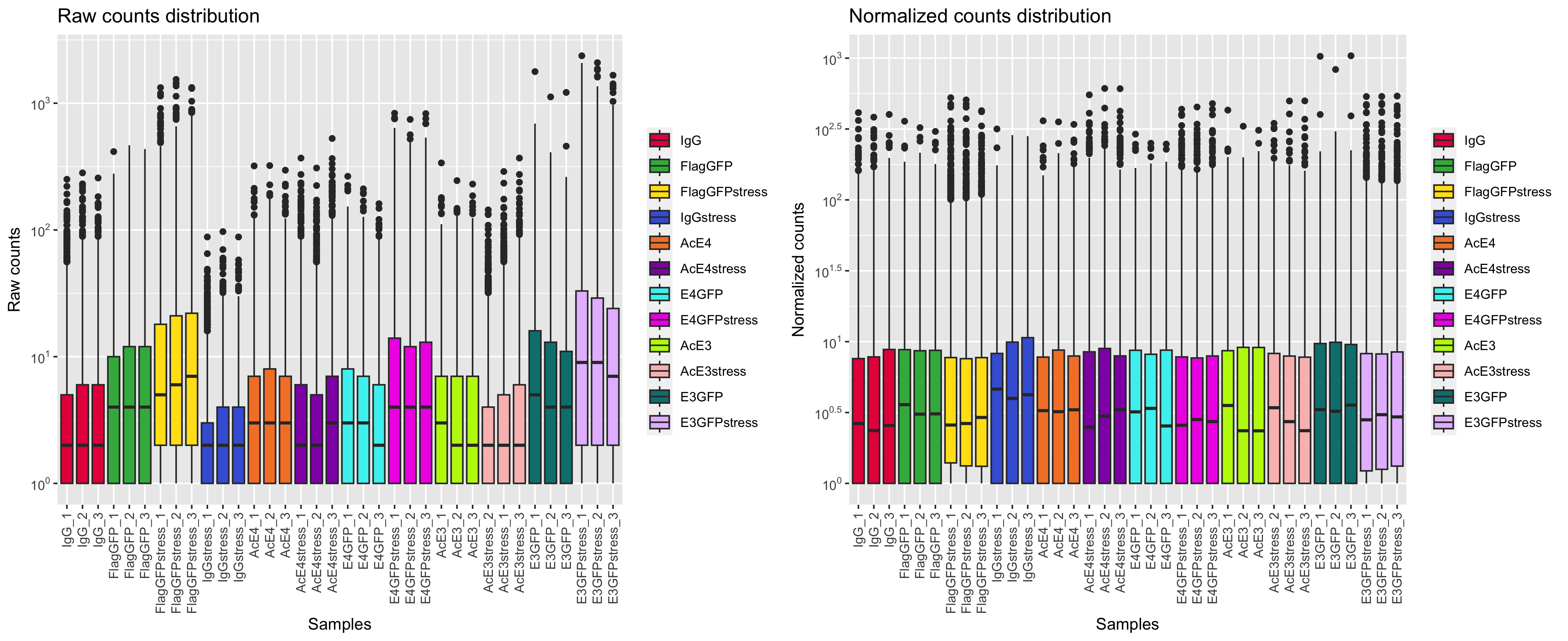

### countsBoxplots.png

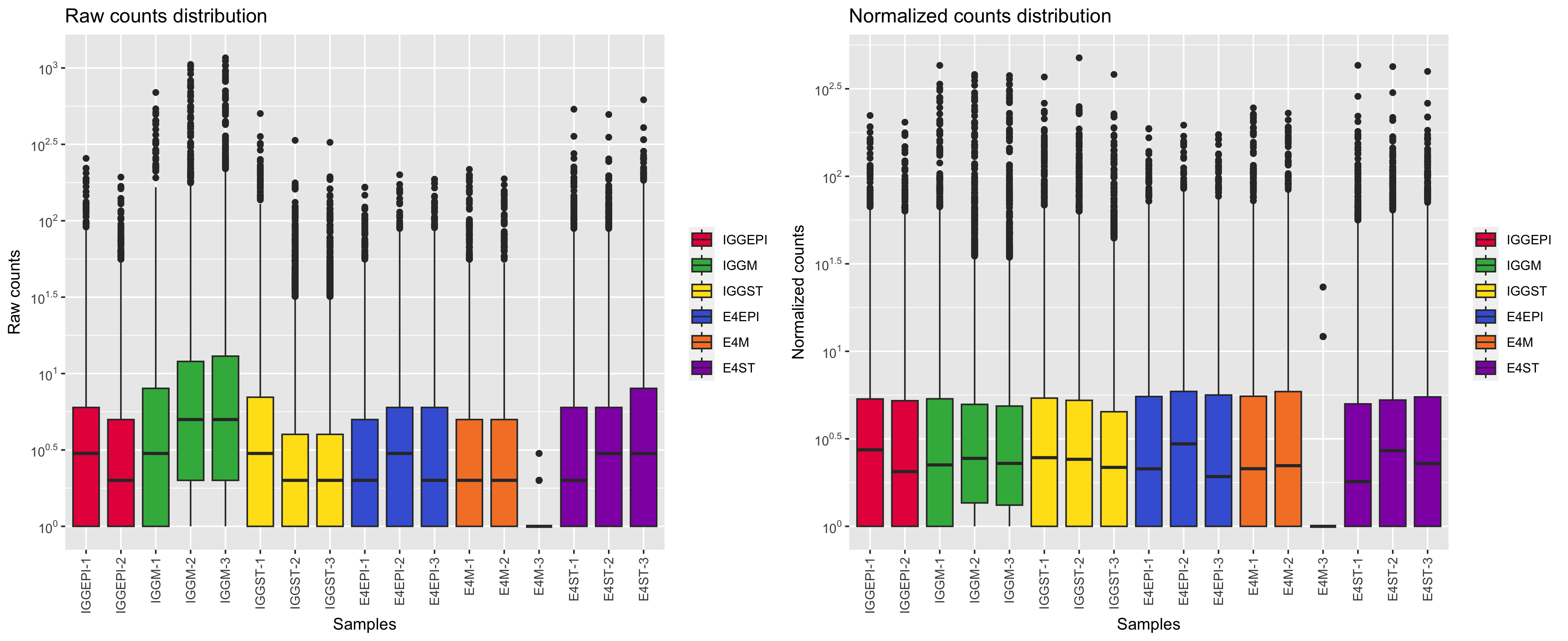

### countsBoxplots.png

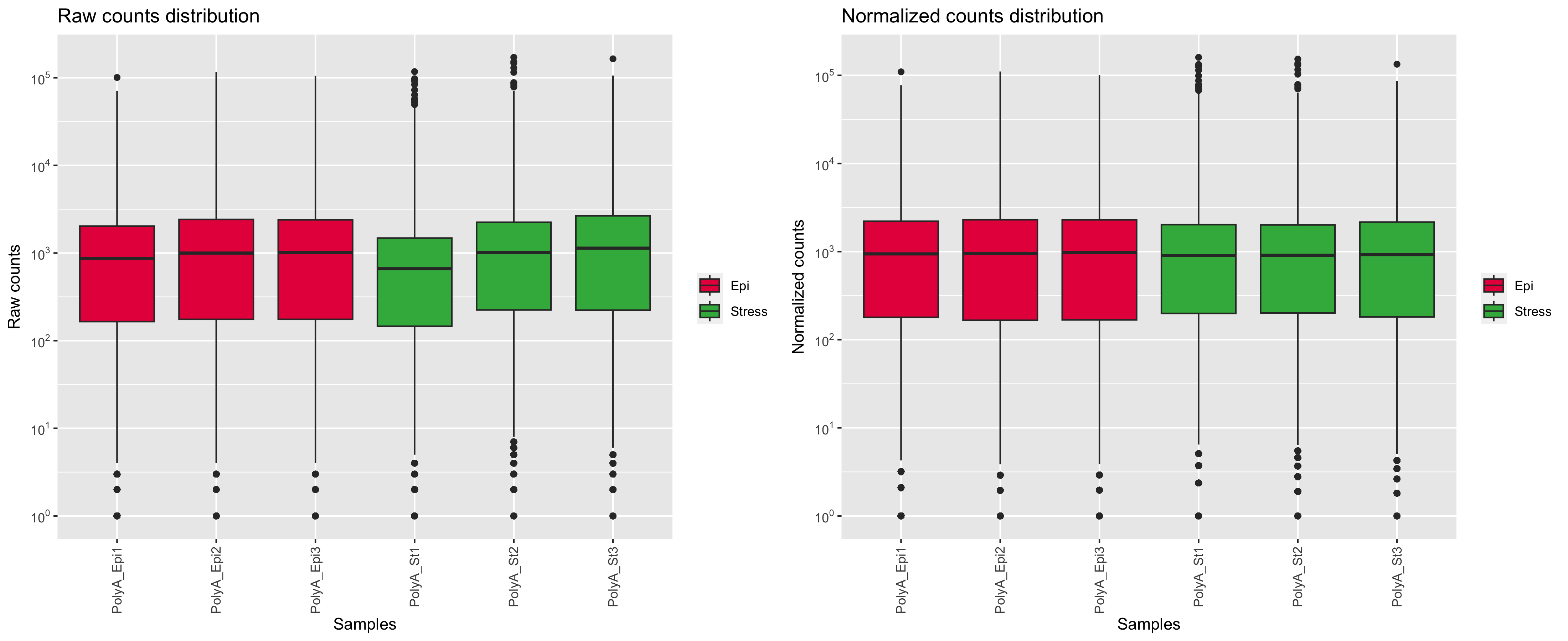

### countsBoxplots.png

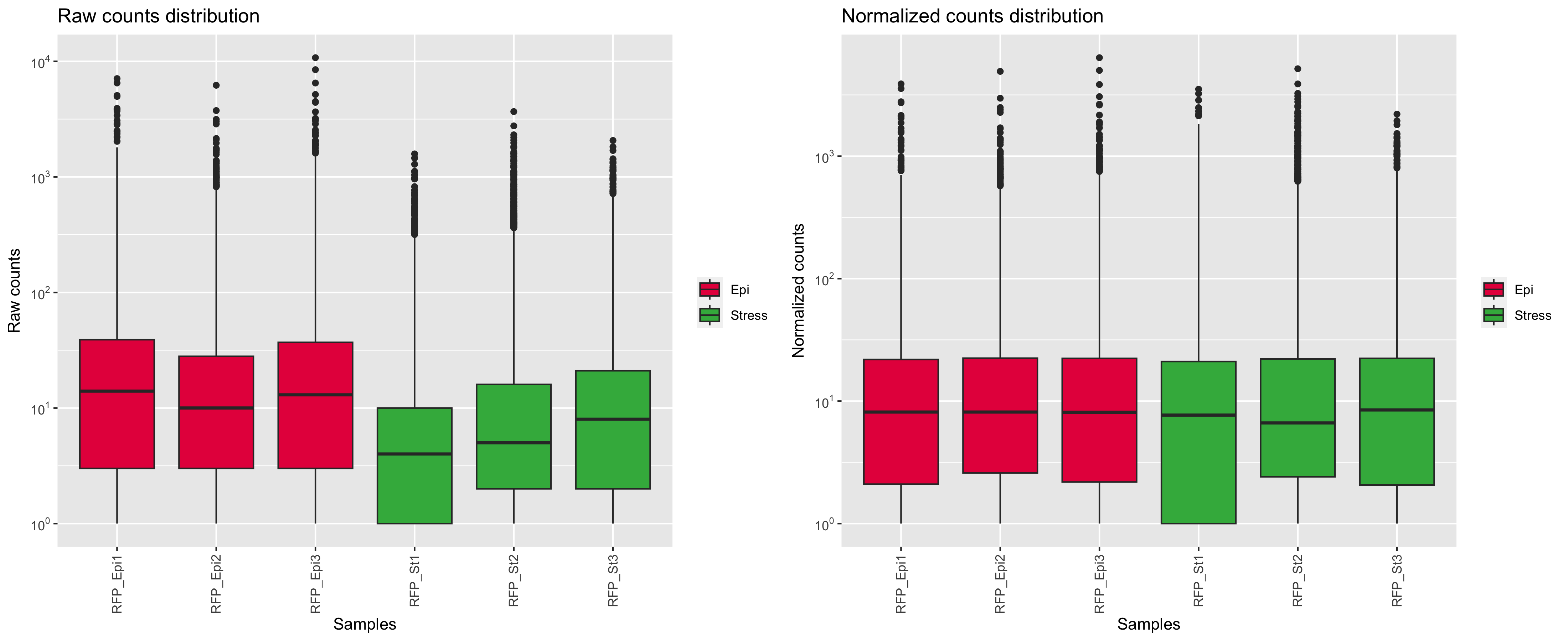

### densplot.png

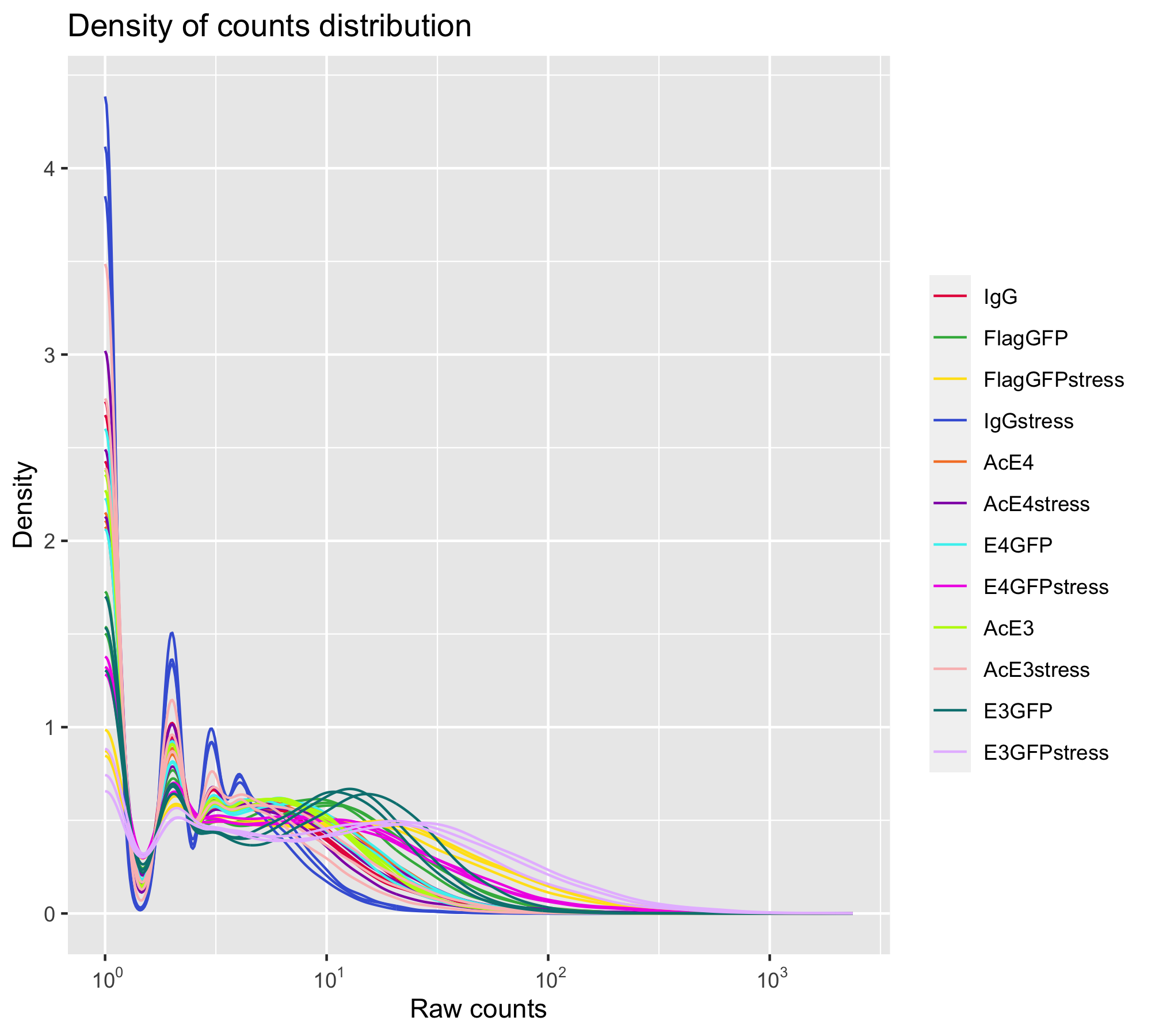

### densplot.png

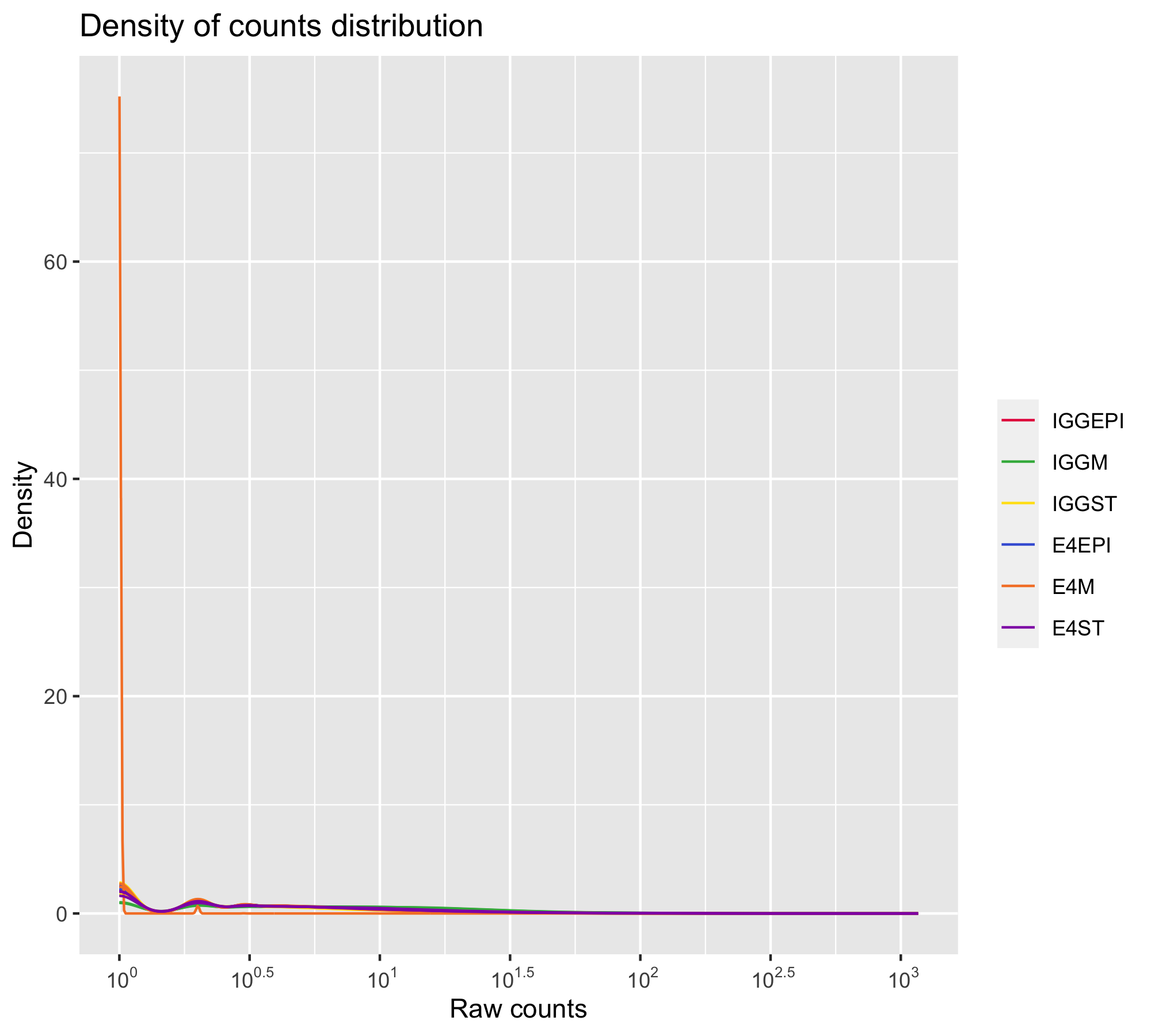

### densplot.png

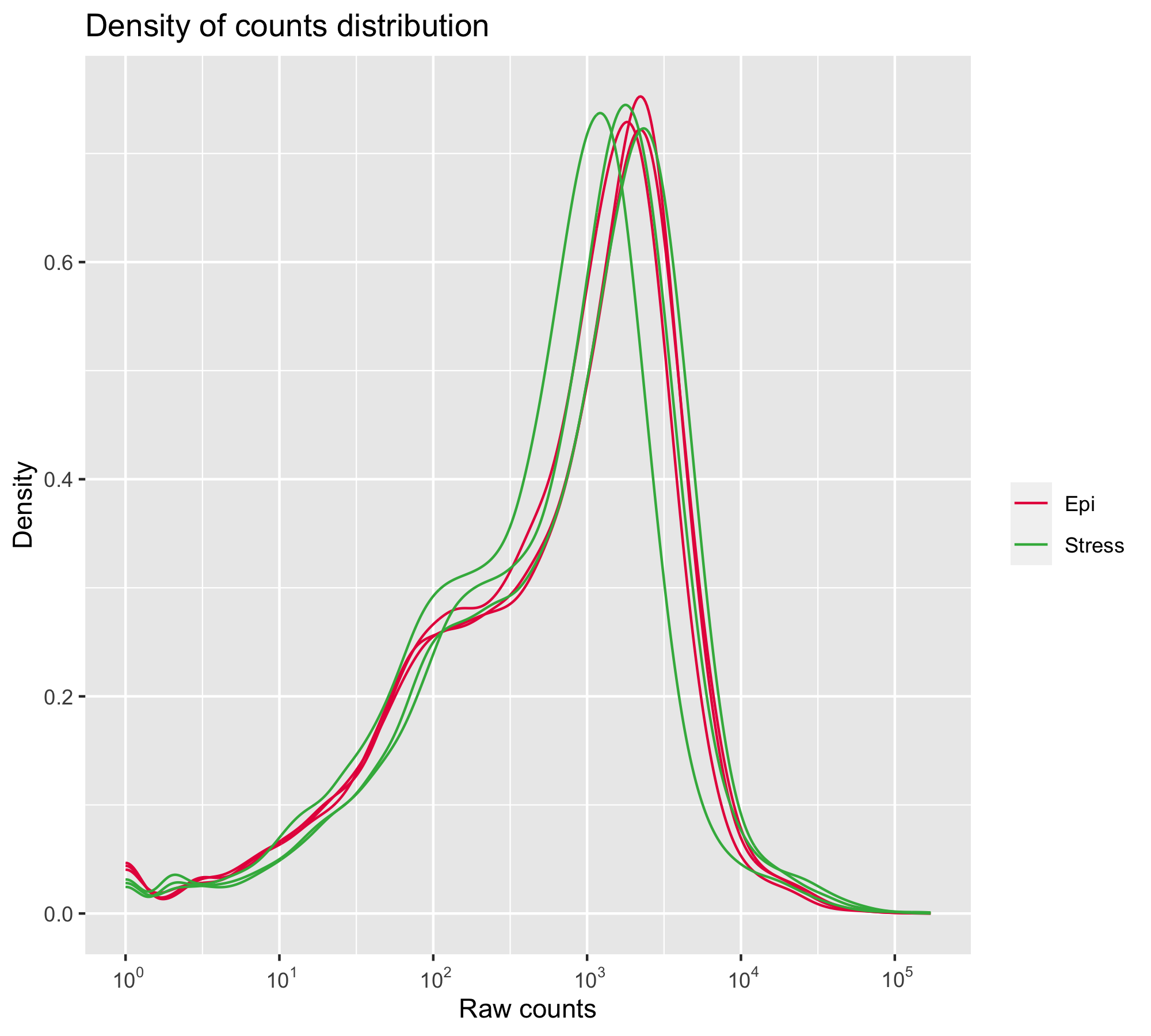

### densplot.png

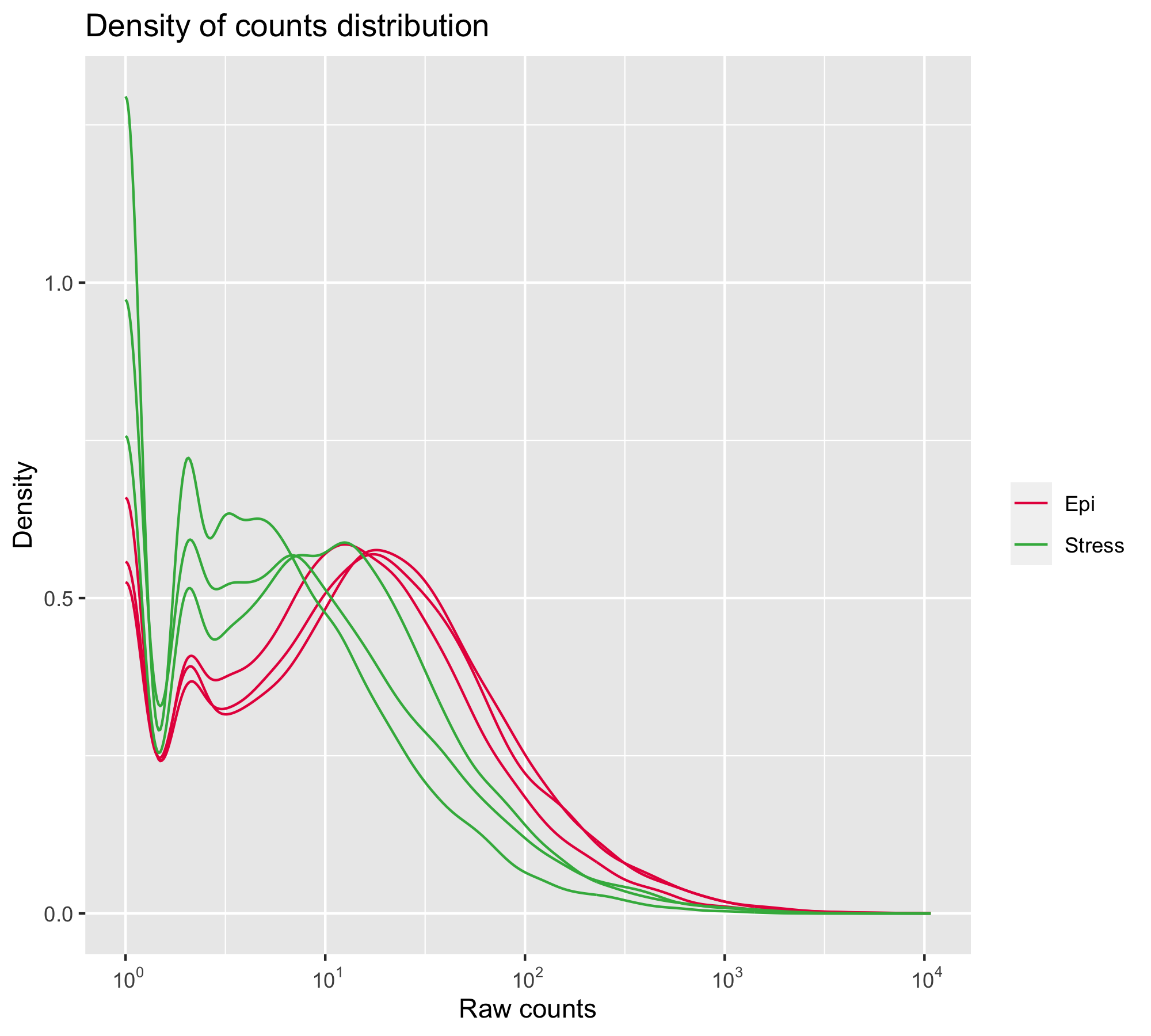

### diagSizeFactorsHist.png

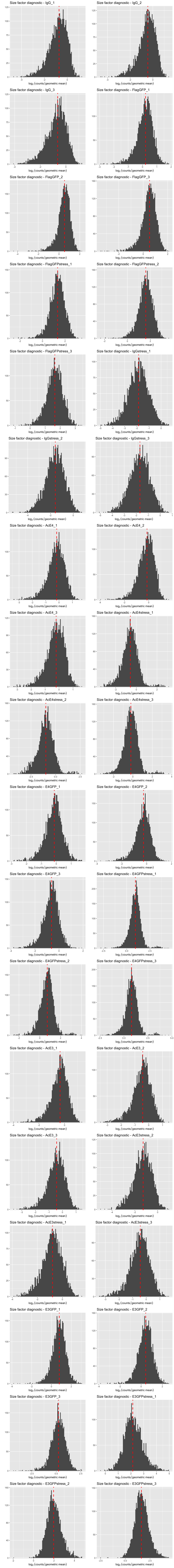

### diagSizeFactorsHist.png

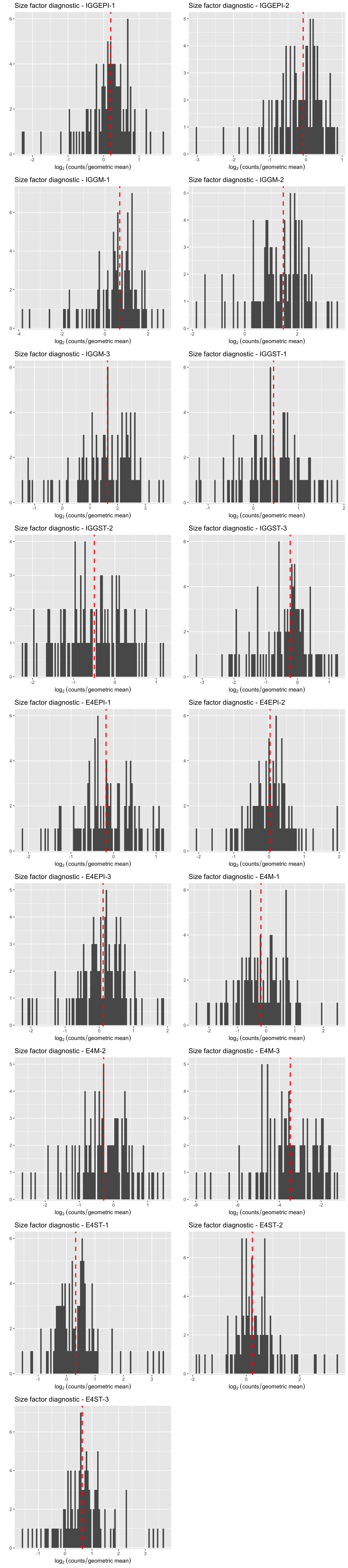

### diagSizeFactorsHist.png

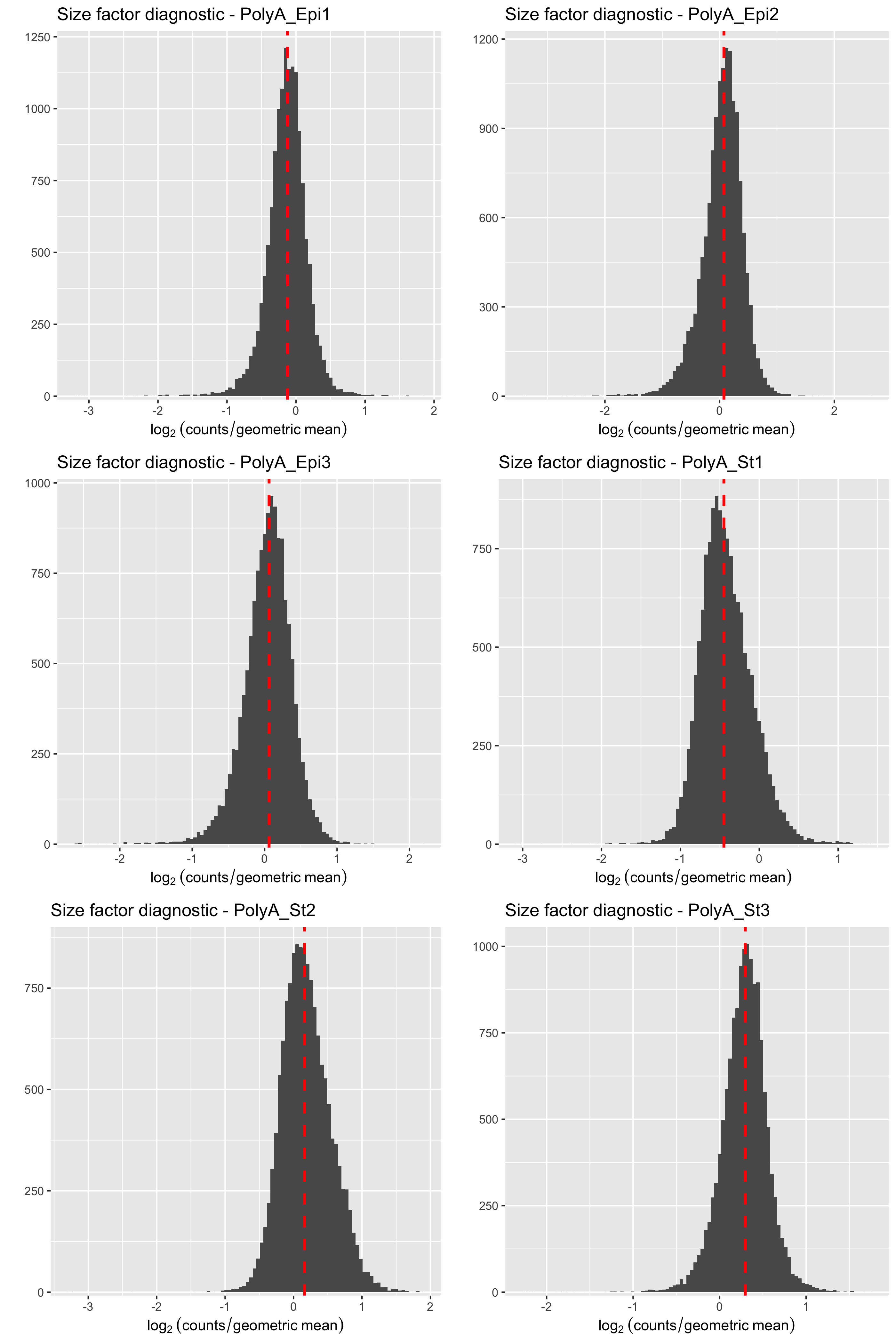

### diagSizeFactorsHist.png

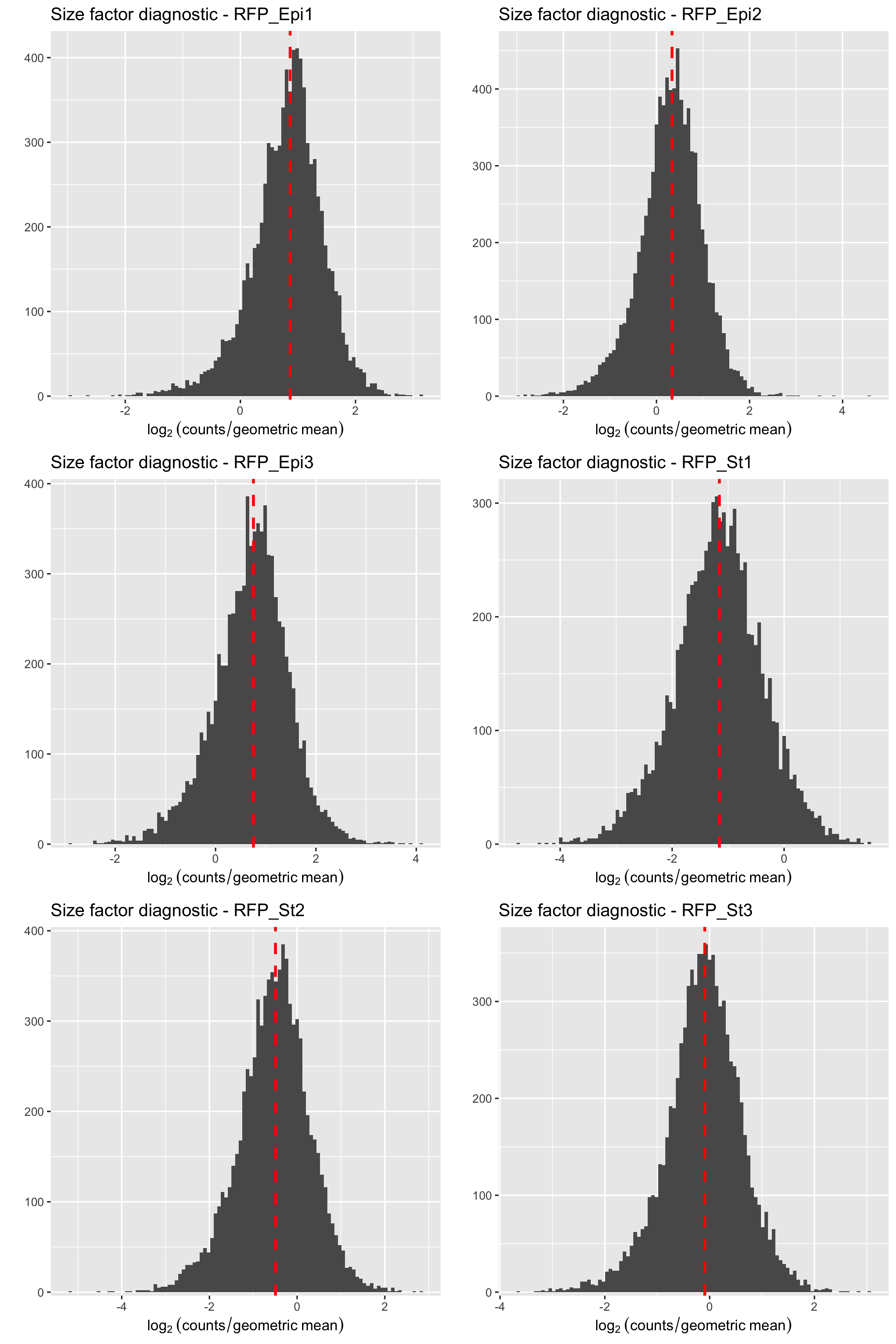

### diagSizeFactorsTC.png

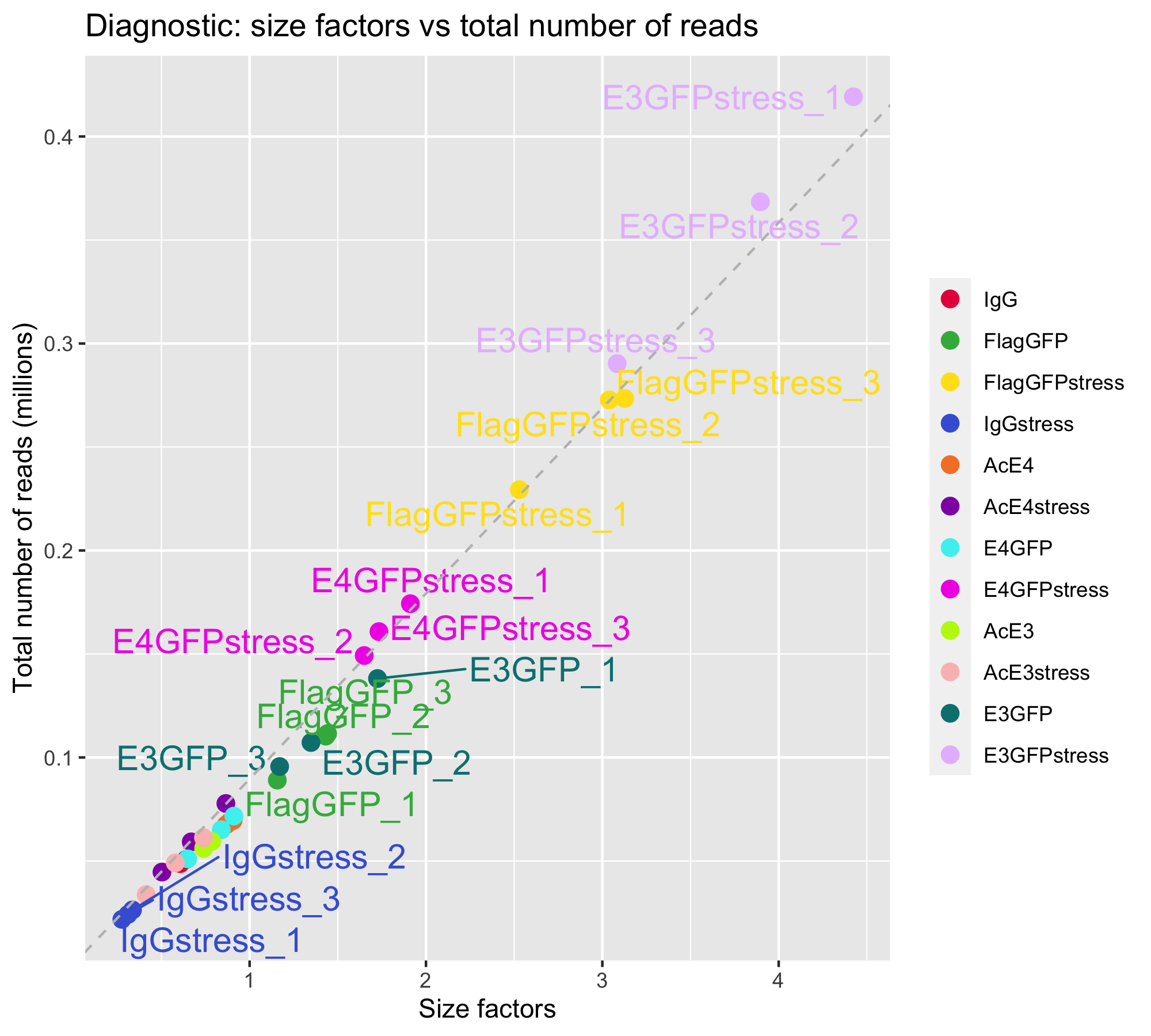

### diagSizeFactorsTC.png

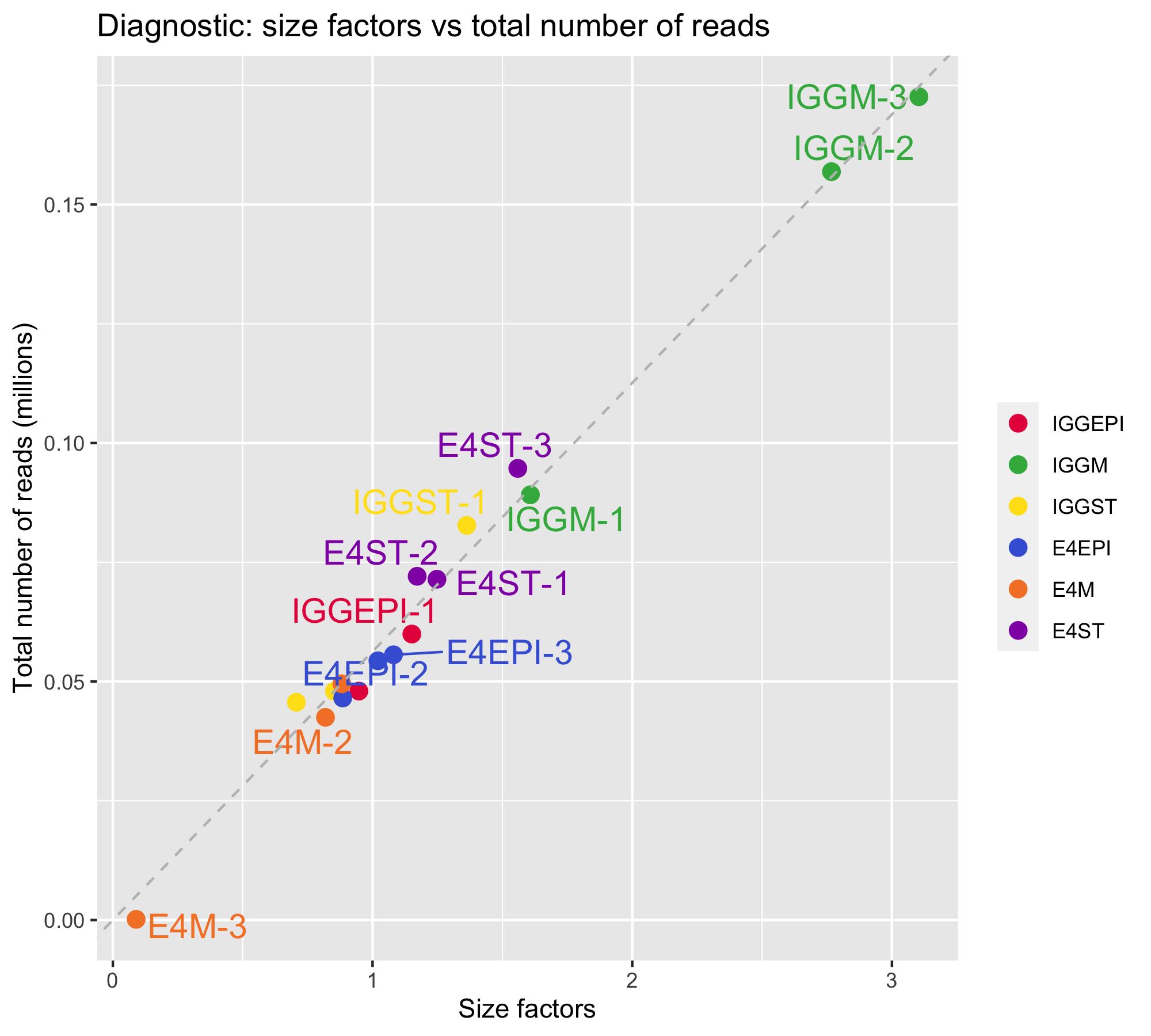

### diagSizeFactorsTC.png

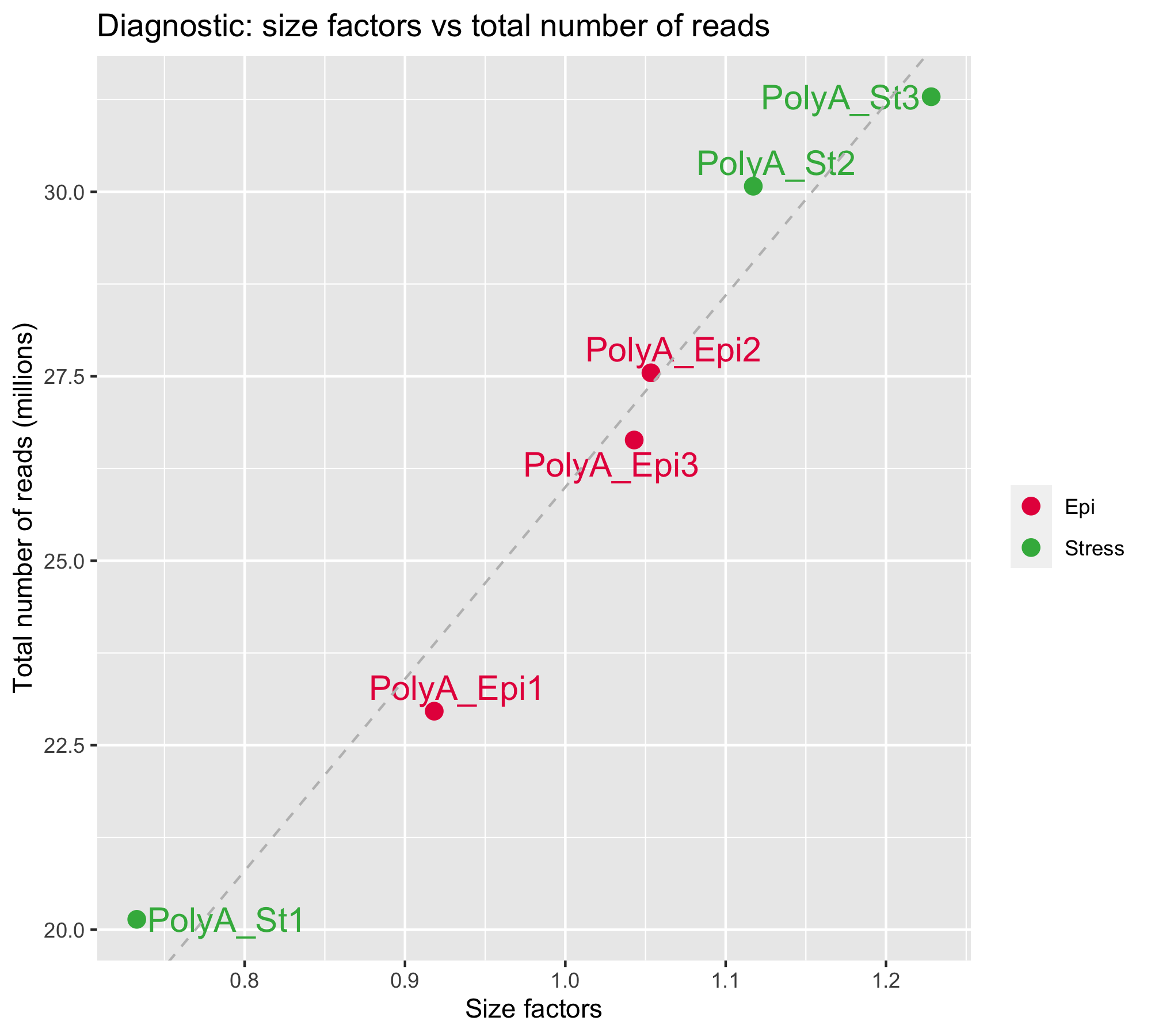

### diagSizeFactorsTC.png

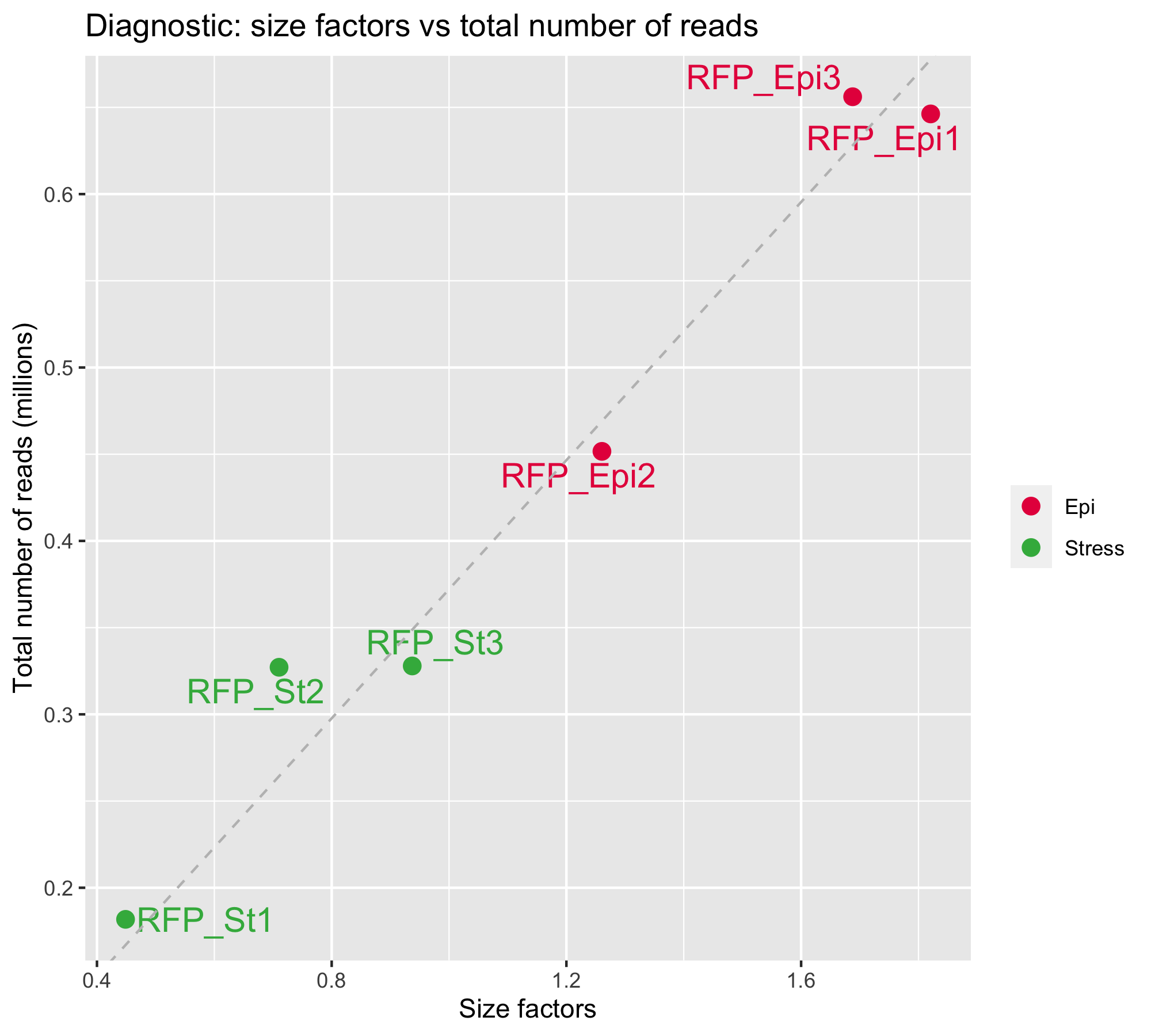

### dispersionsPlot.png

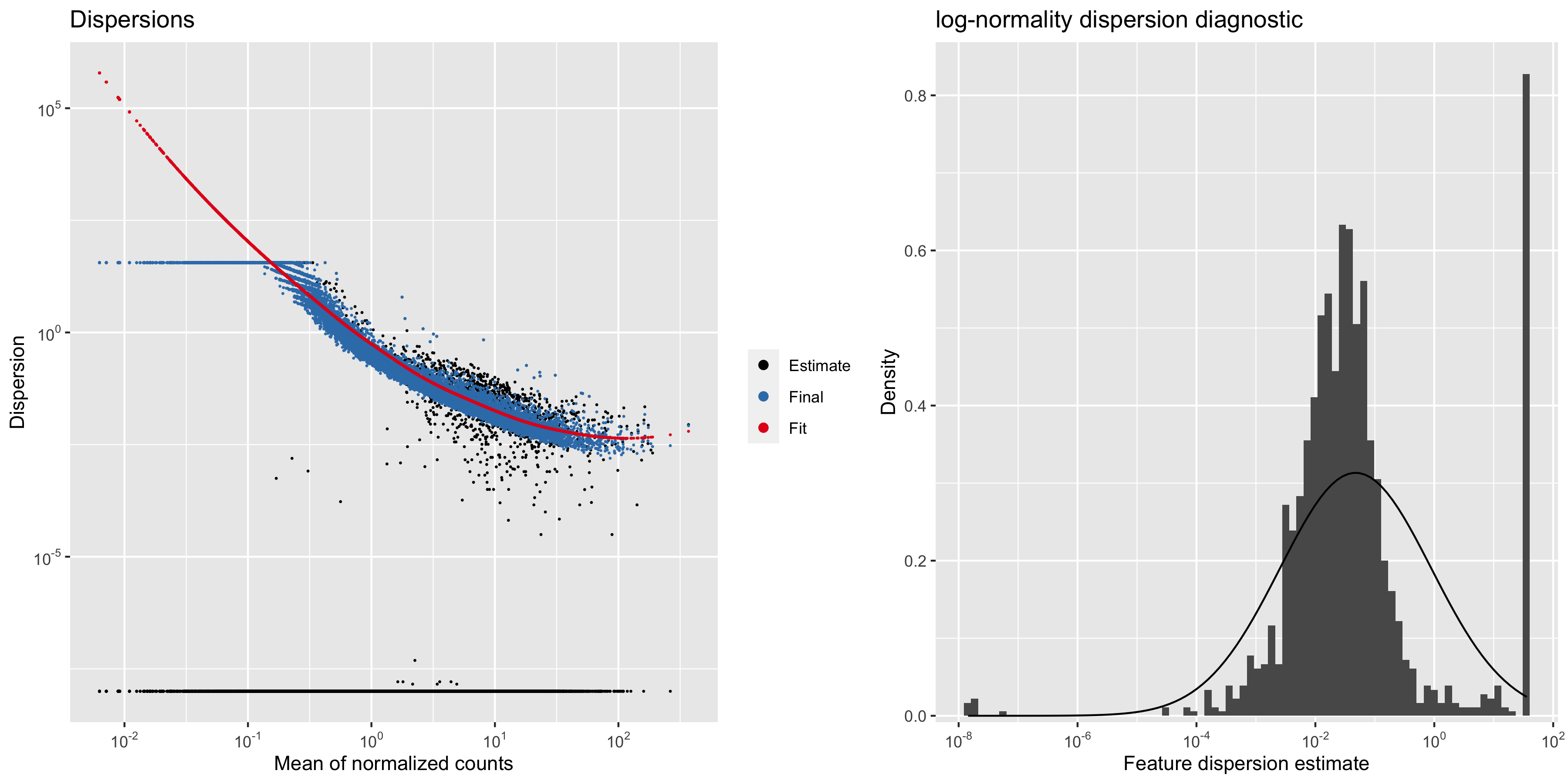

### dispersionsPlot.png

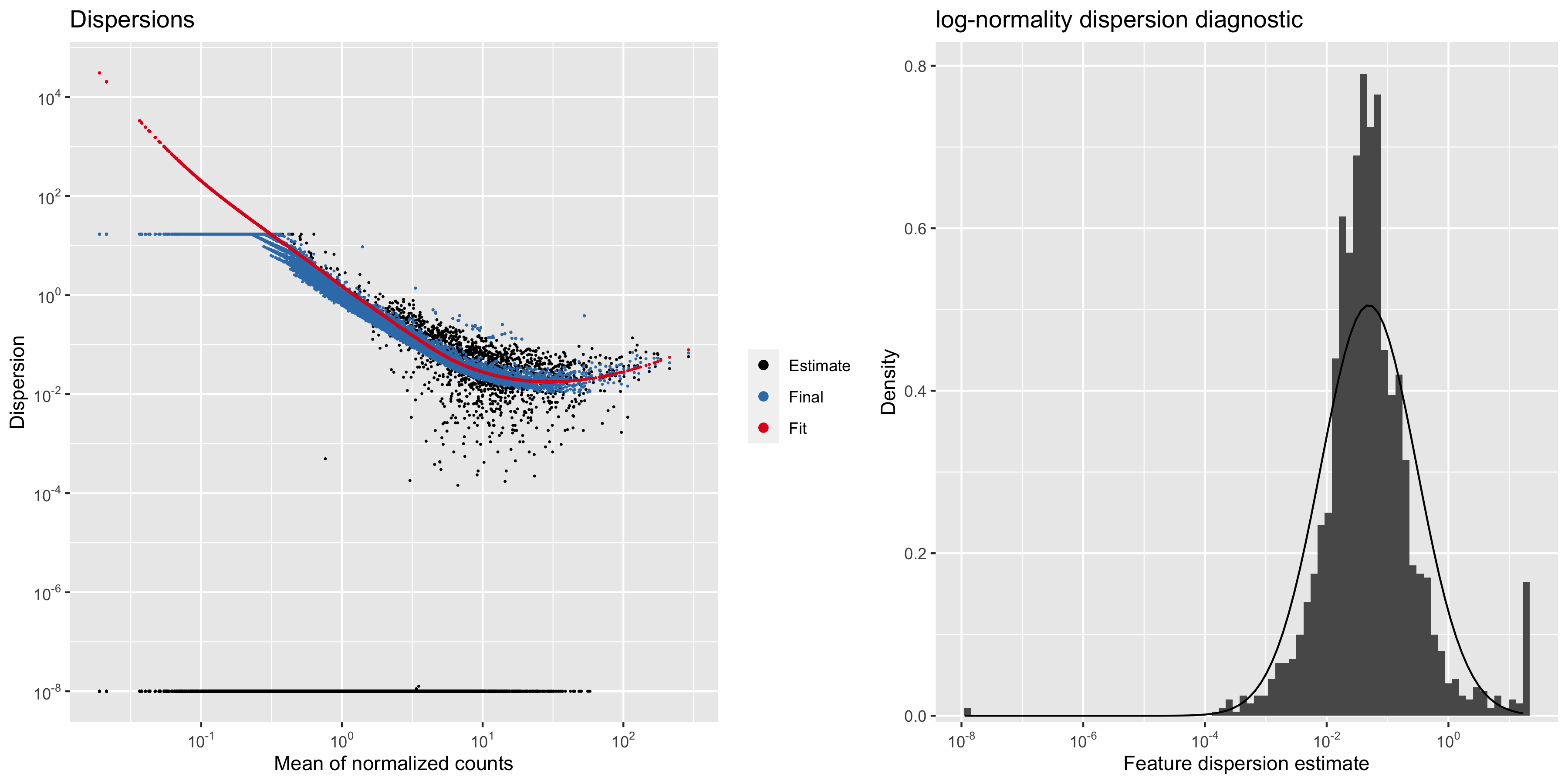

### dispersionsPlot.png

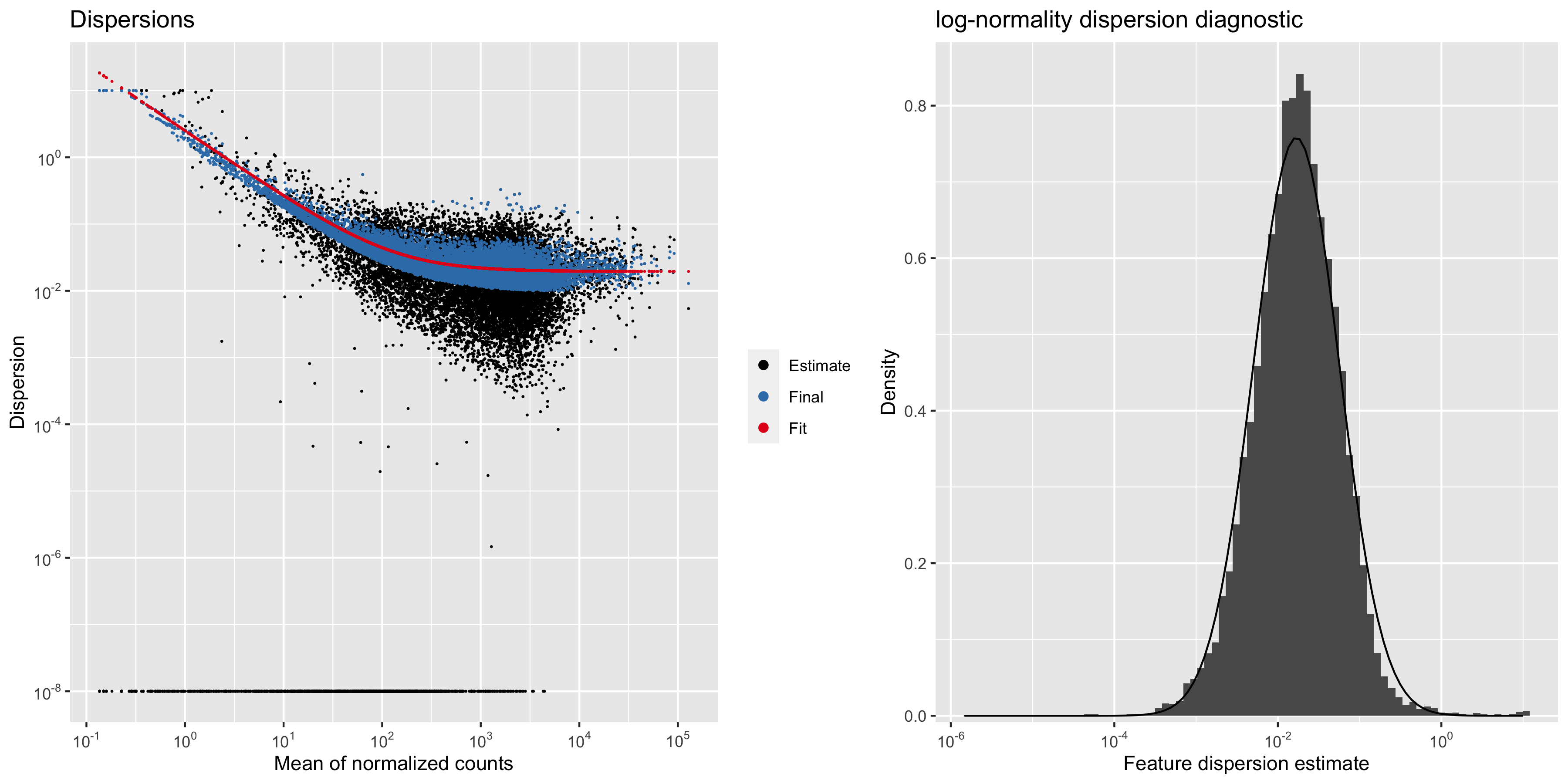

### dispersionsPlot.png

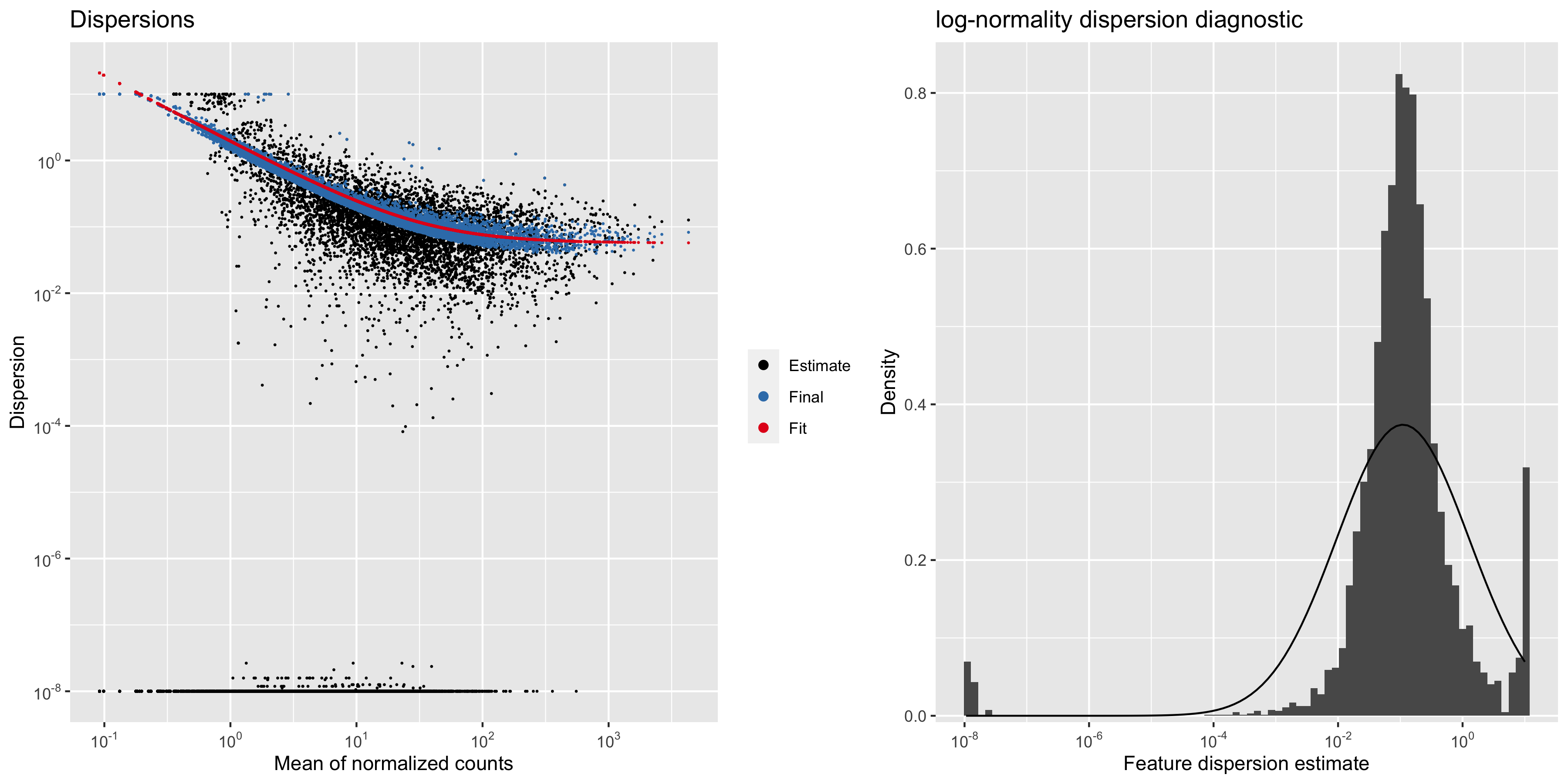

### Figure S4

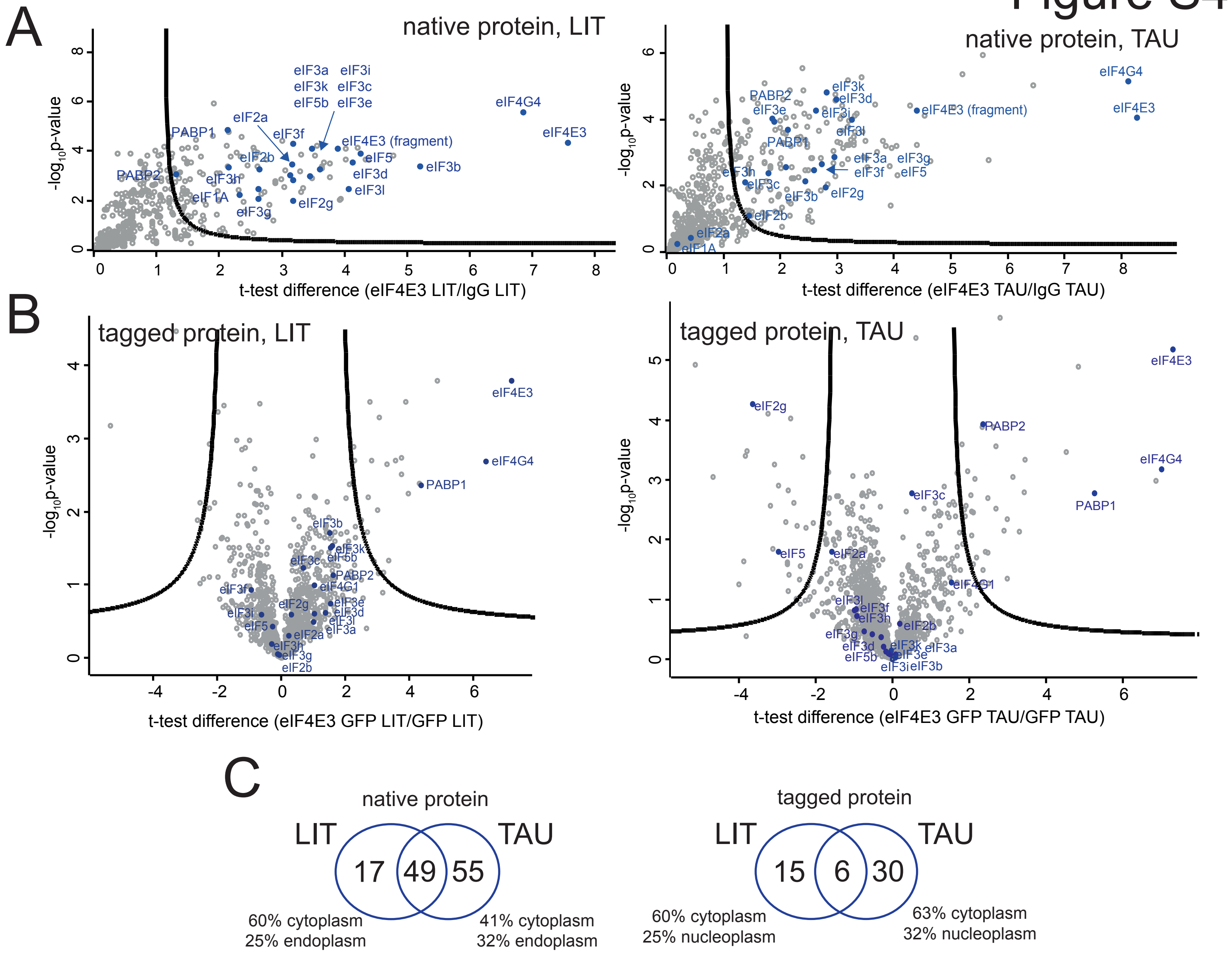

### Figure S5

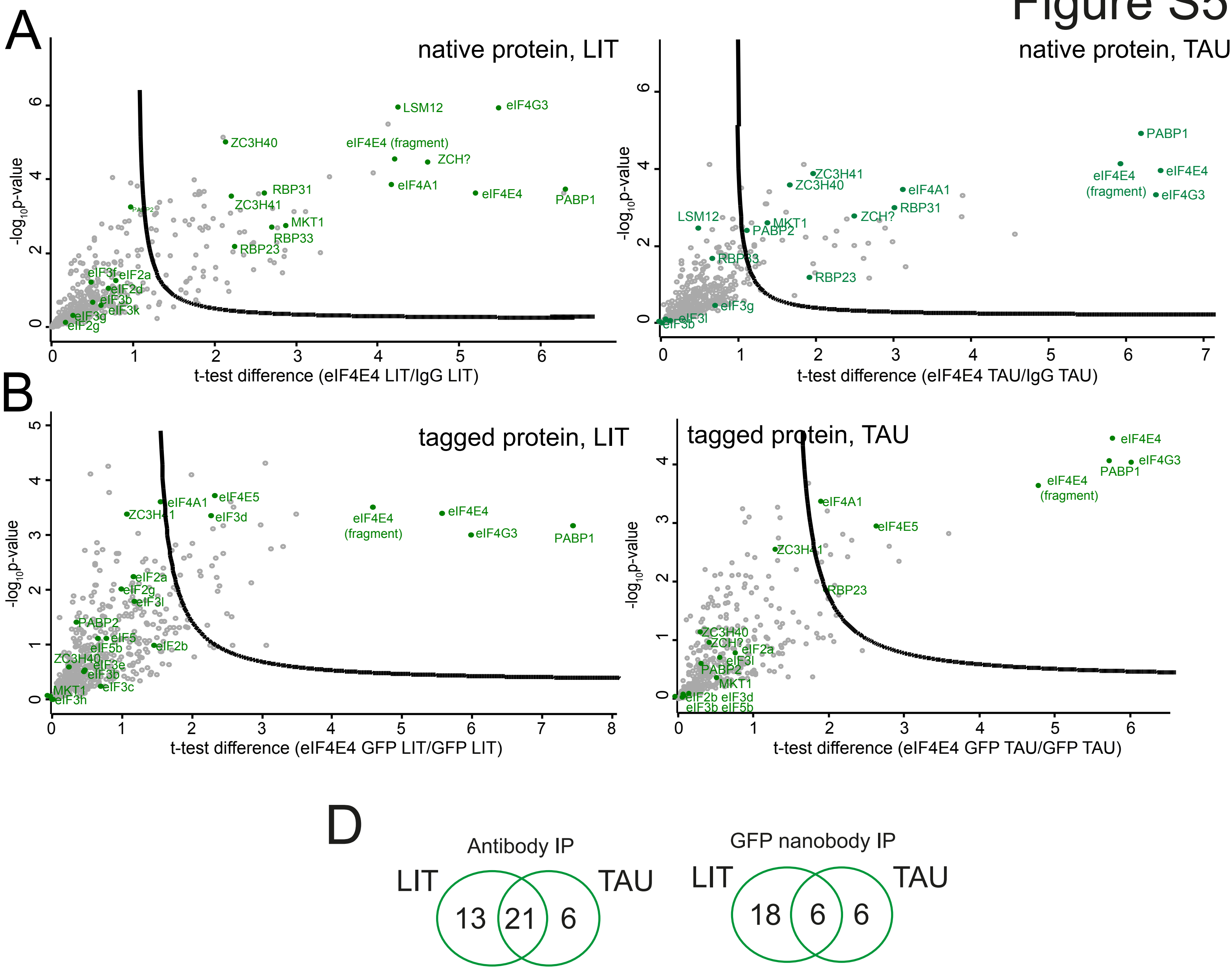

### Figure S6

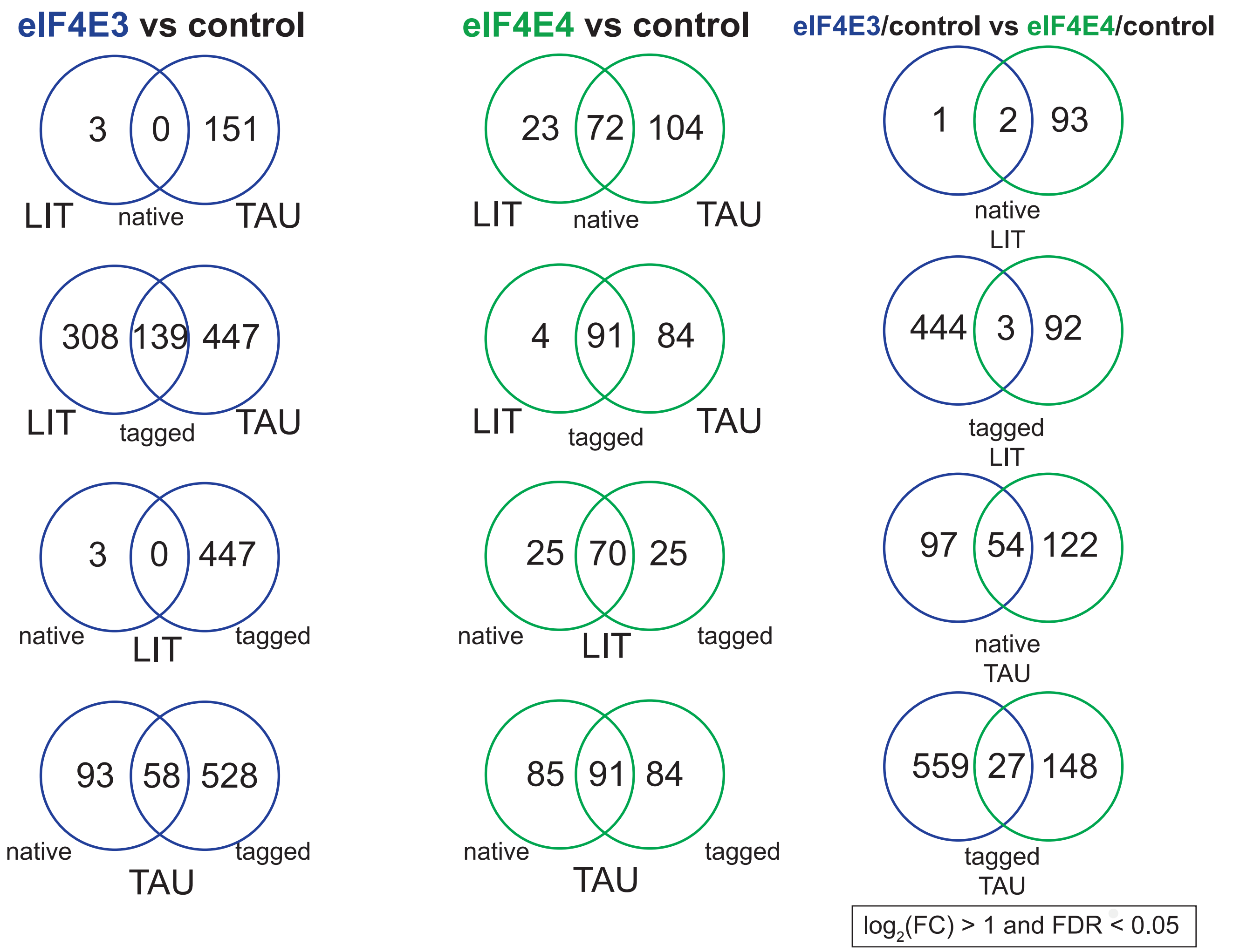

### Figure S7

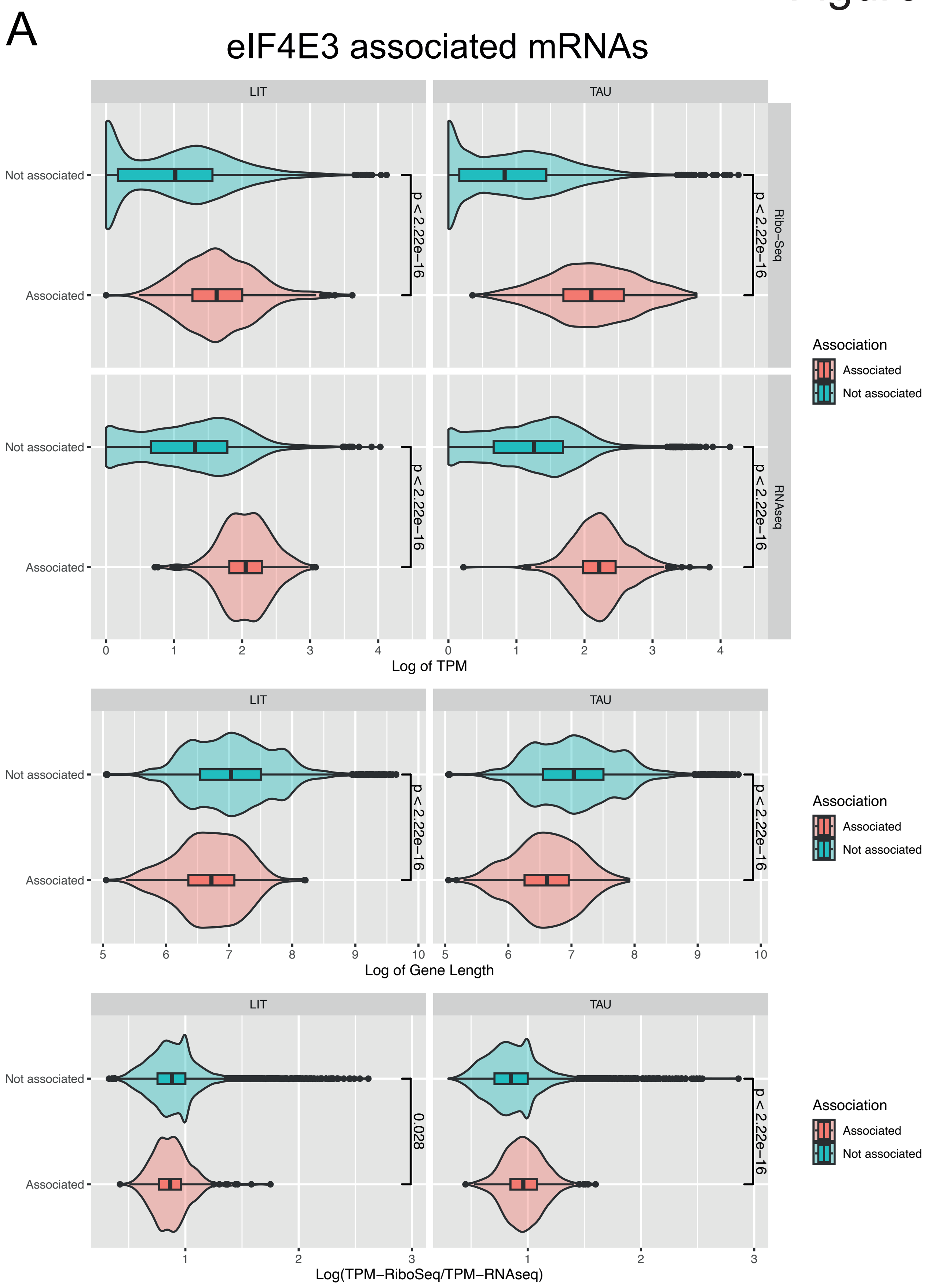

#### Figure S7 (cont.)

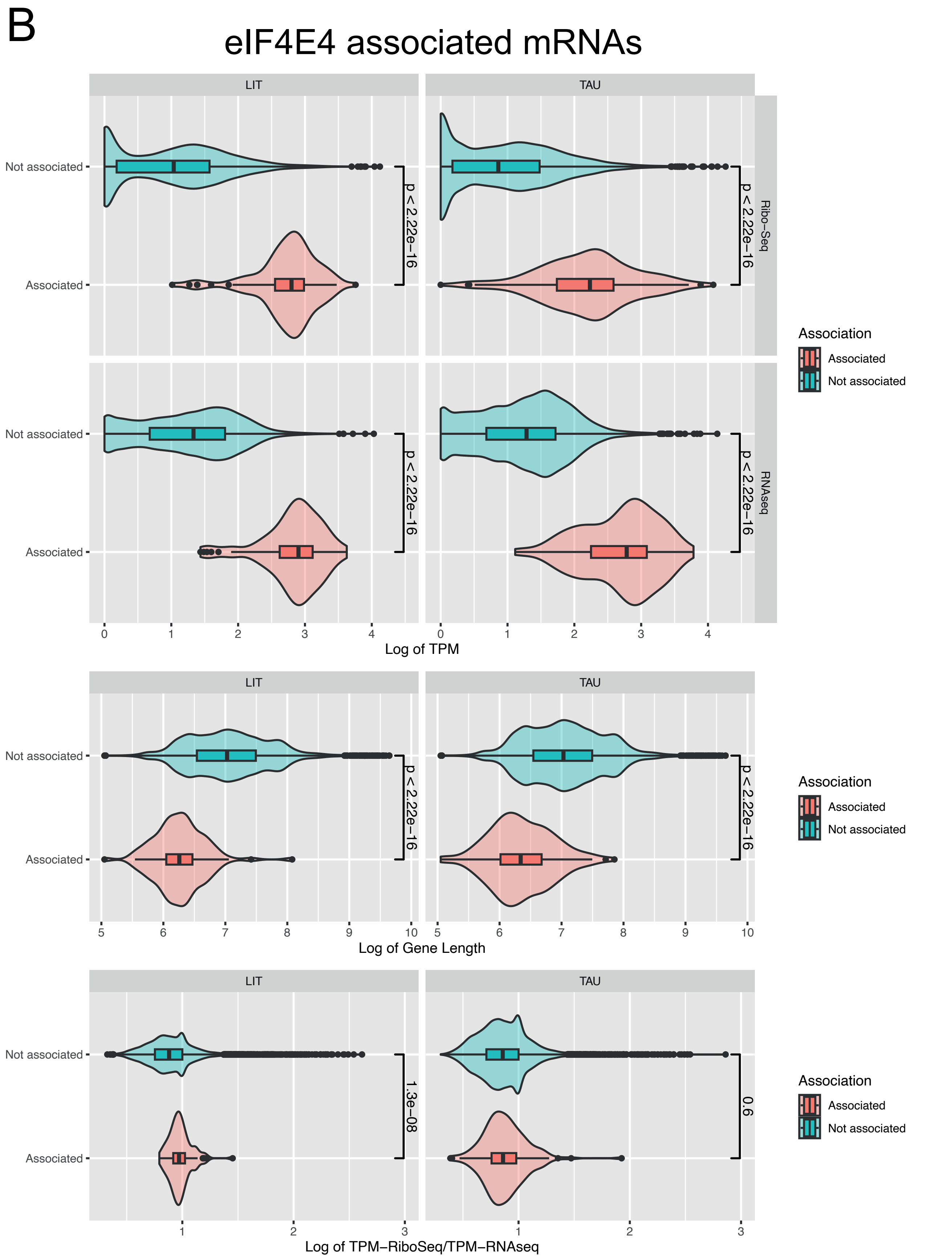

### Figure S8

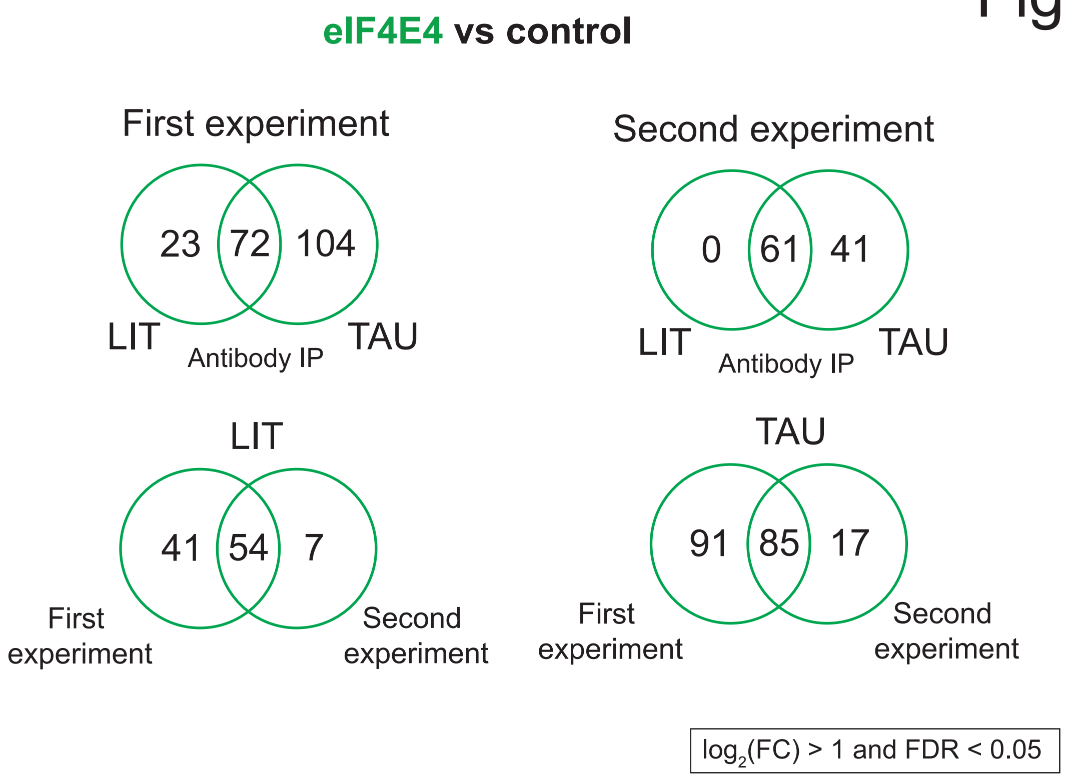

### majSeq.png

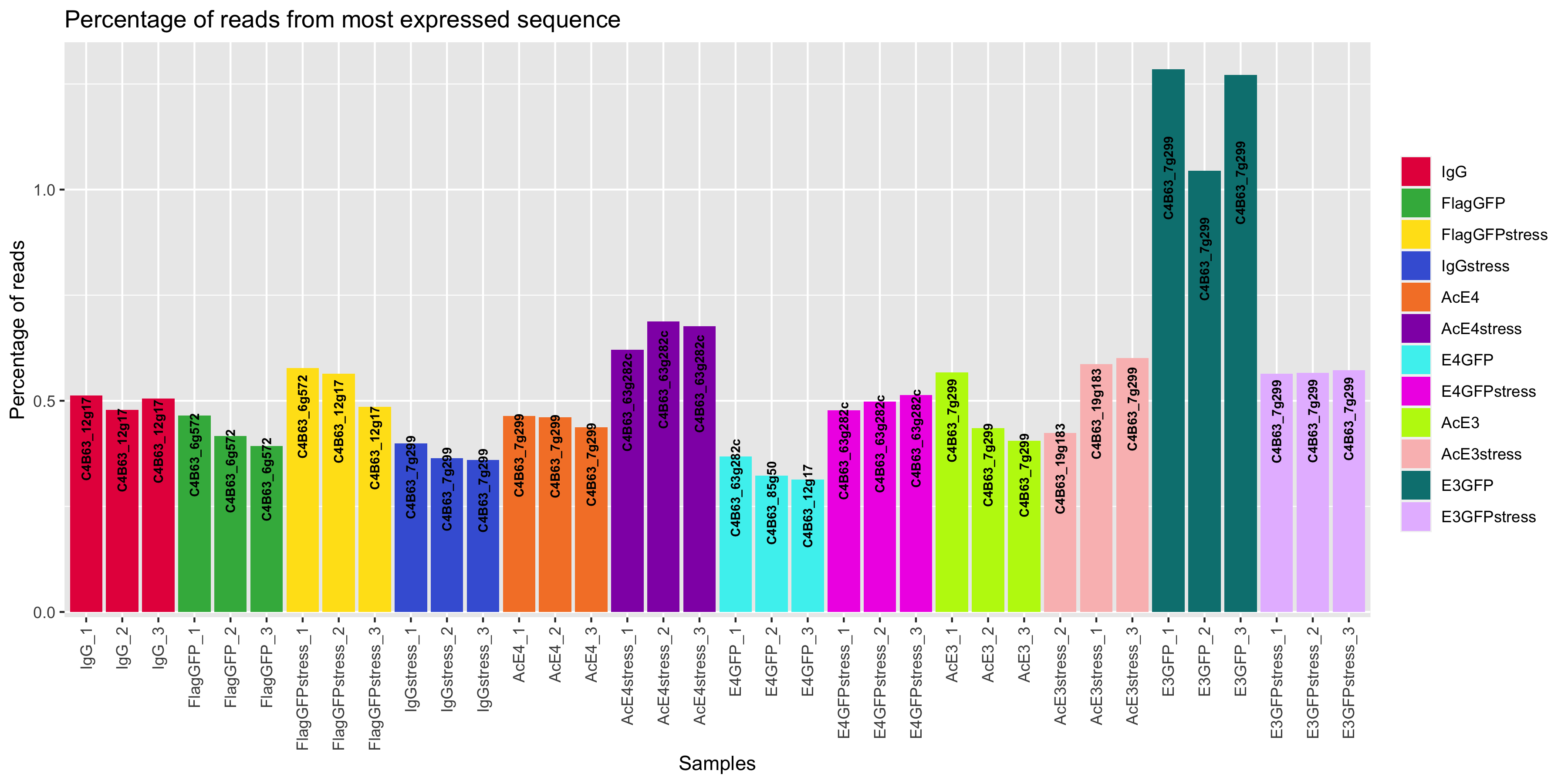

### majSeq.png

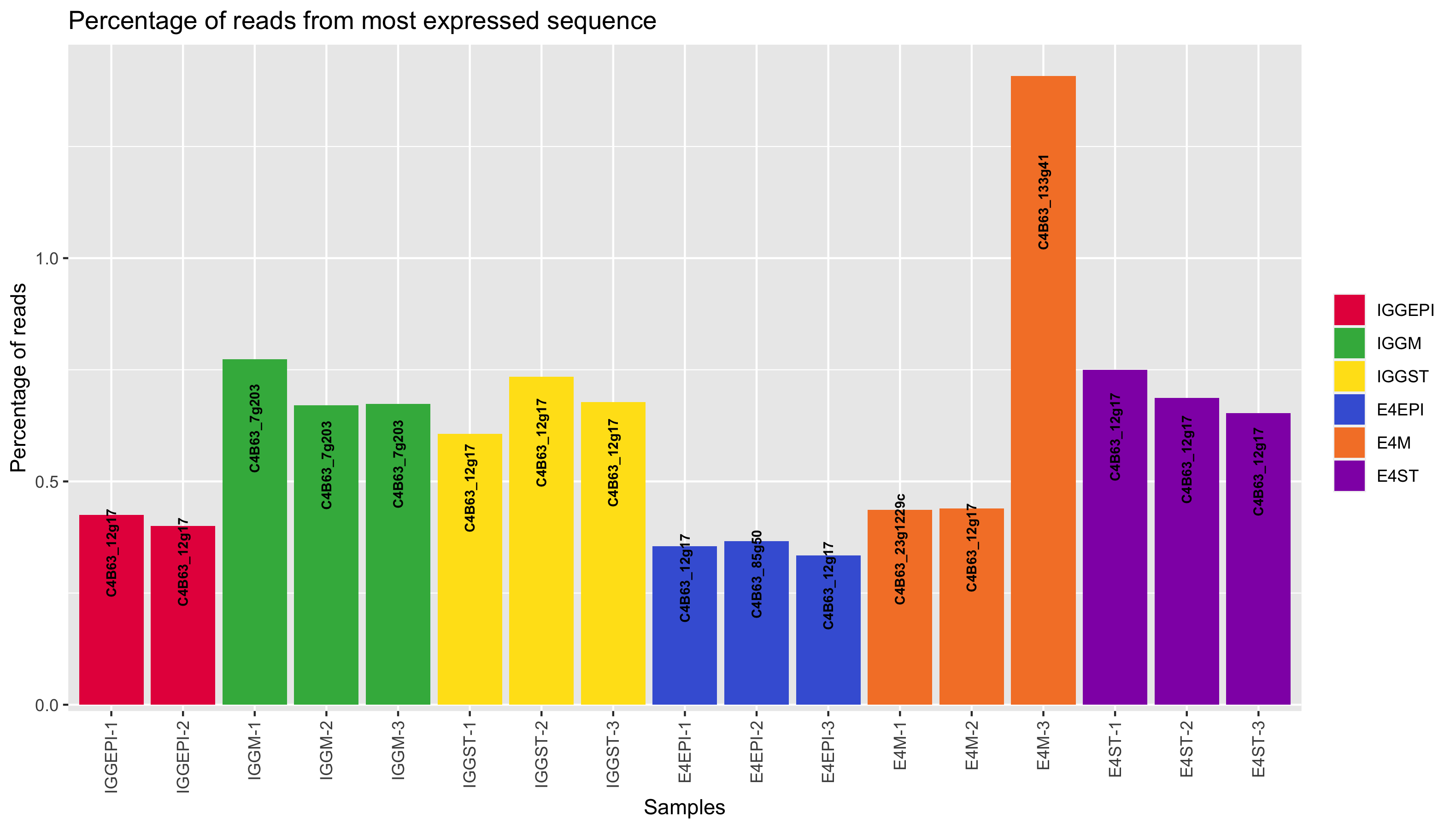

### majSeq.png

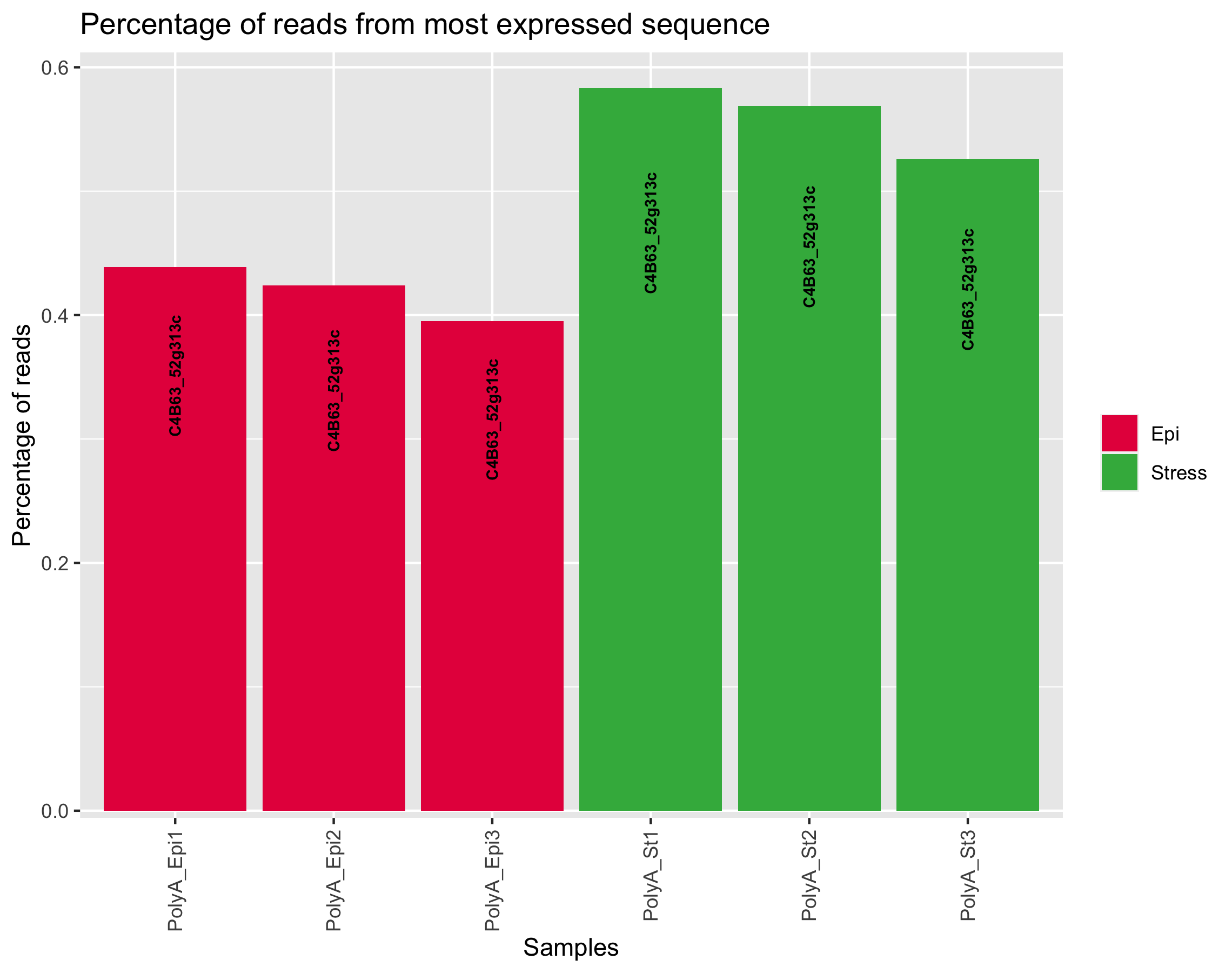

### majSeq.png

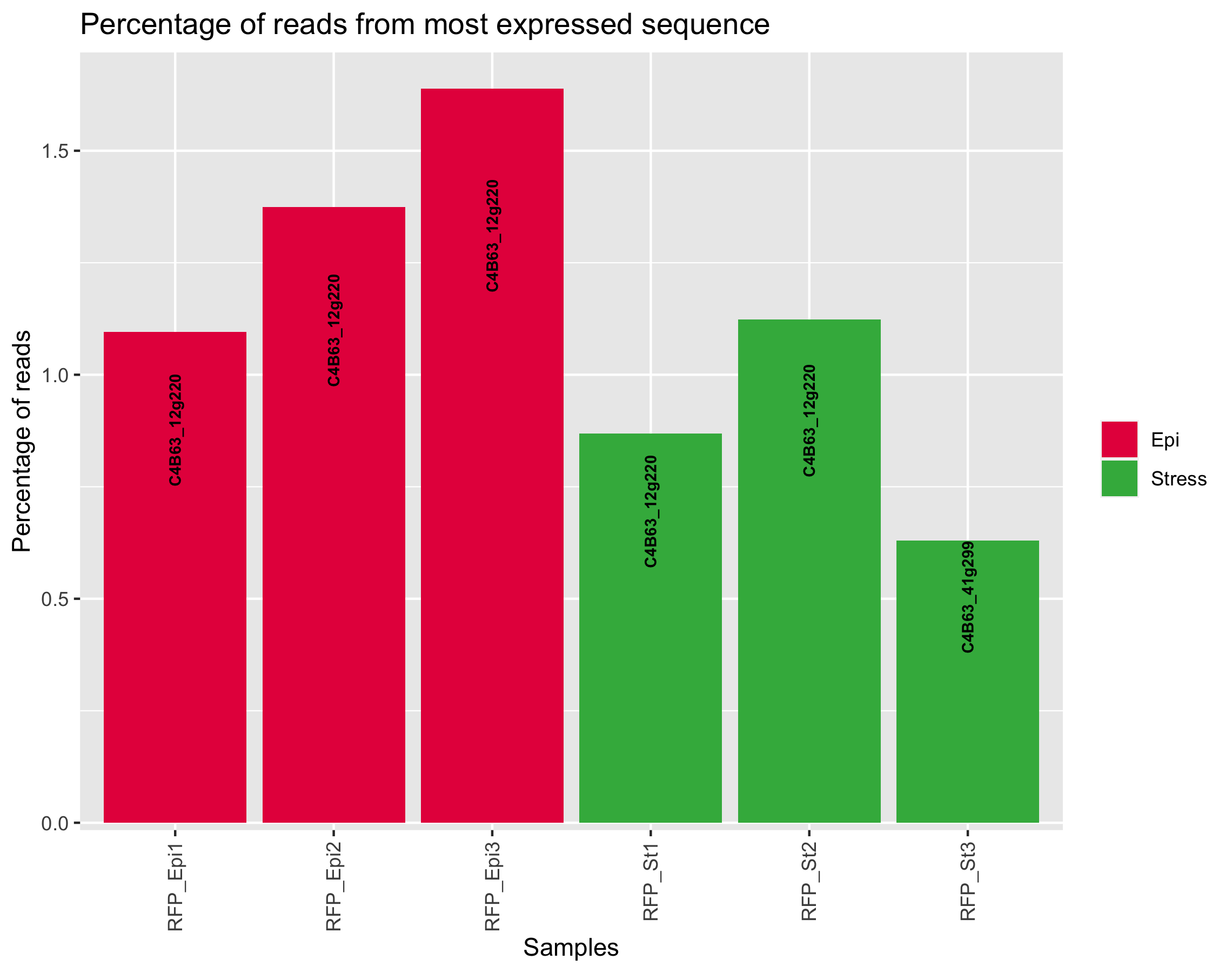

### MAPlot.png

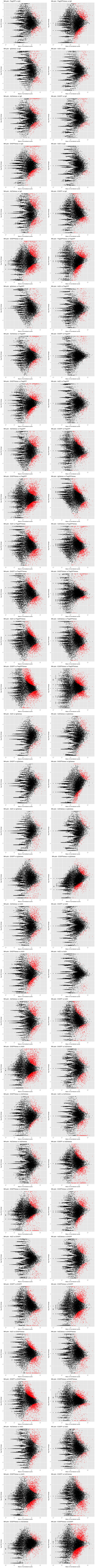

### MAPlot.png

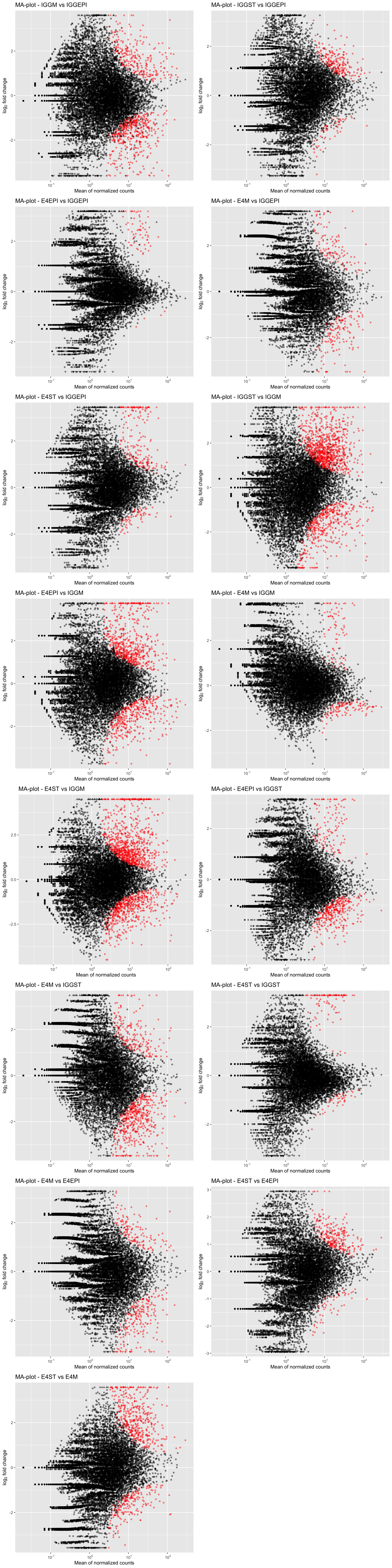

### MAPlot.png

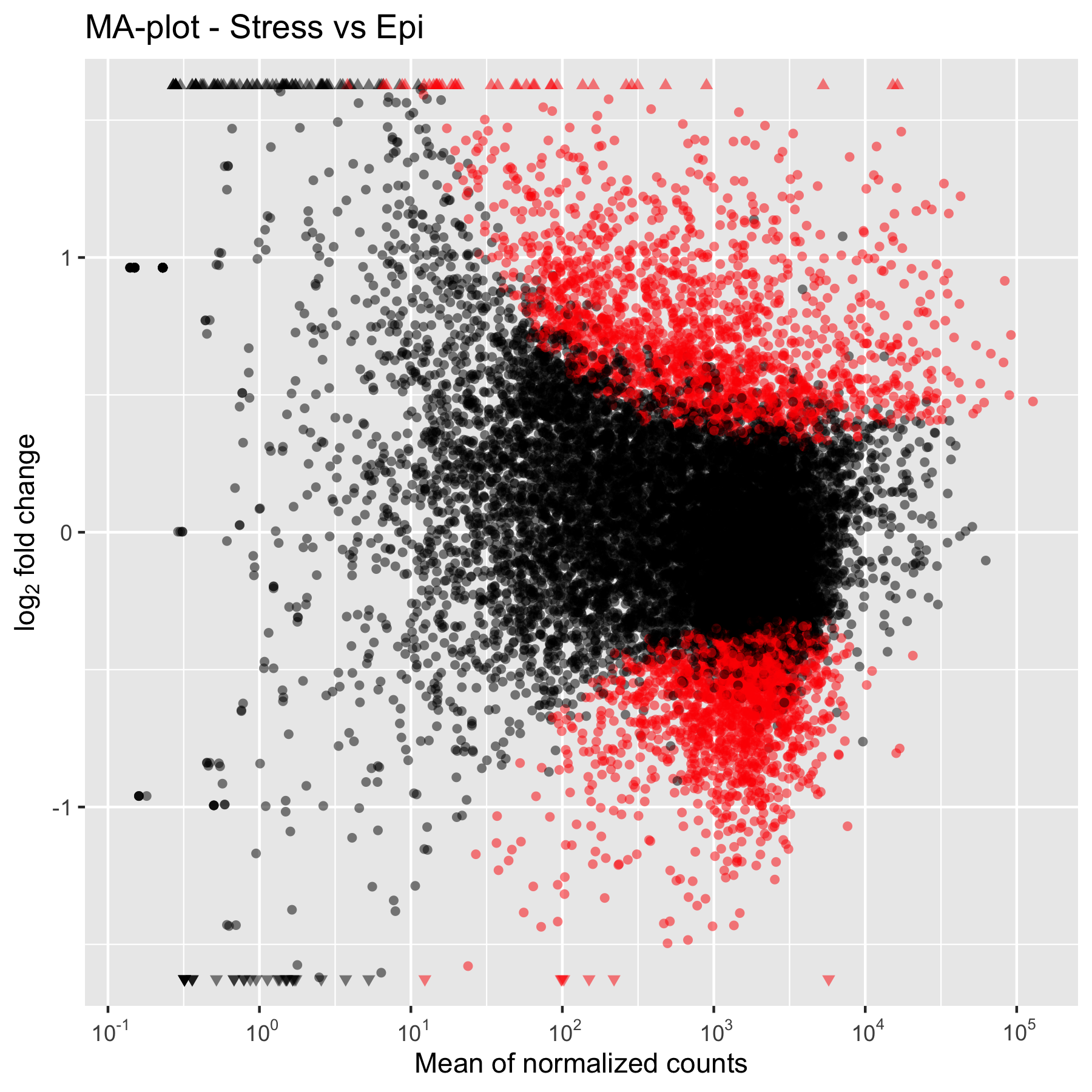

### MAPlot.png

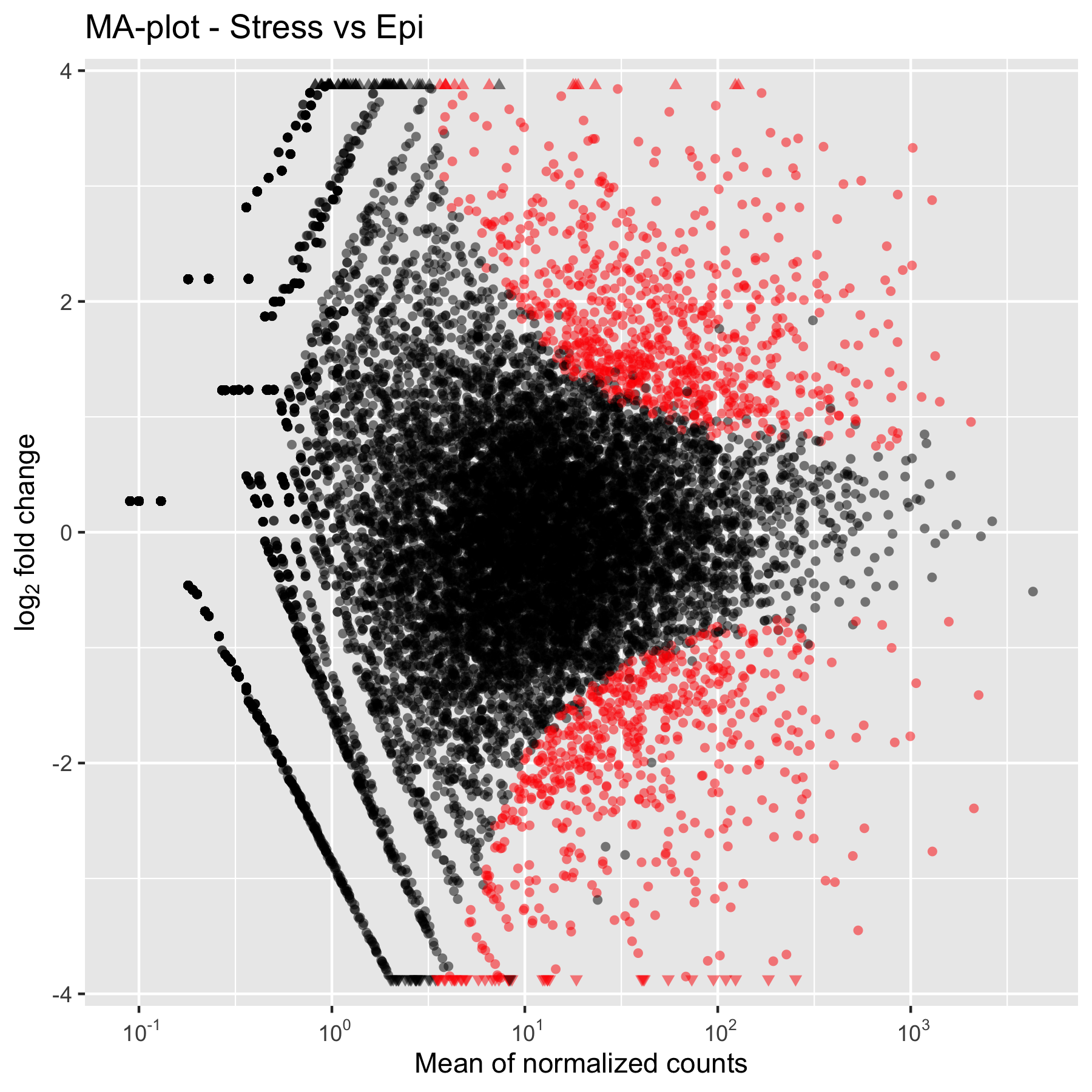

### pairwiseScatter.png

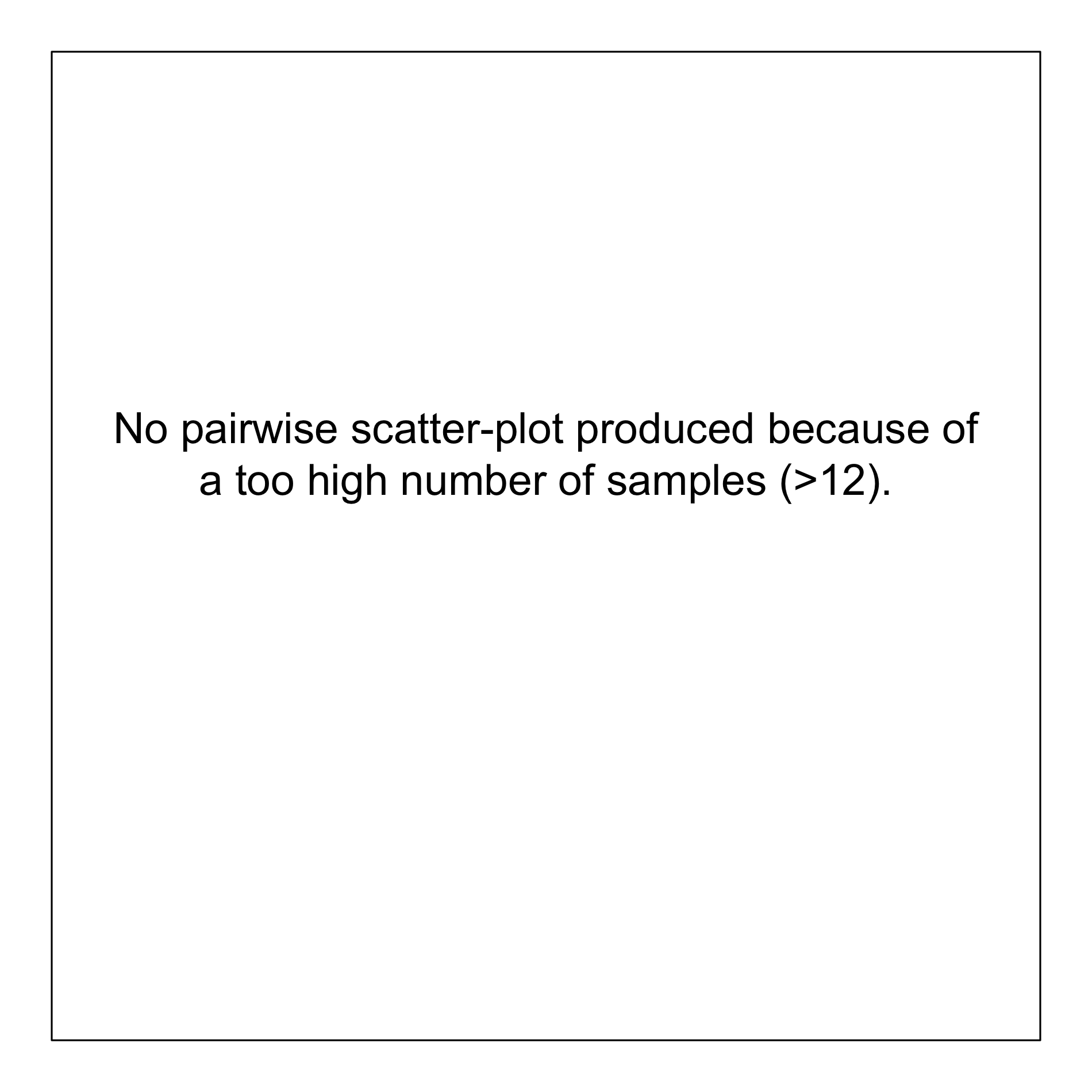

### pairwiseScatter.png

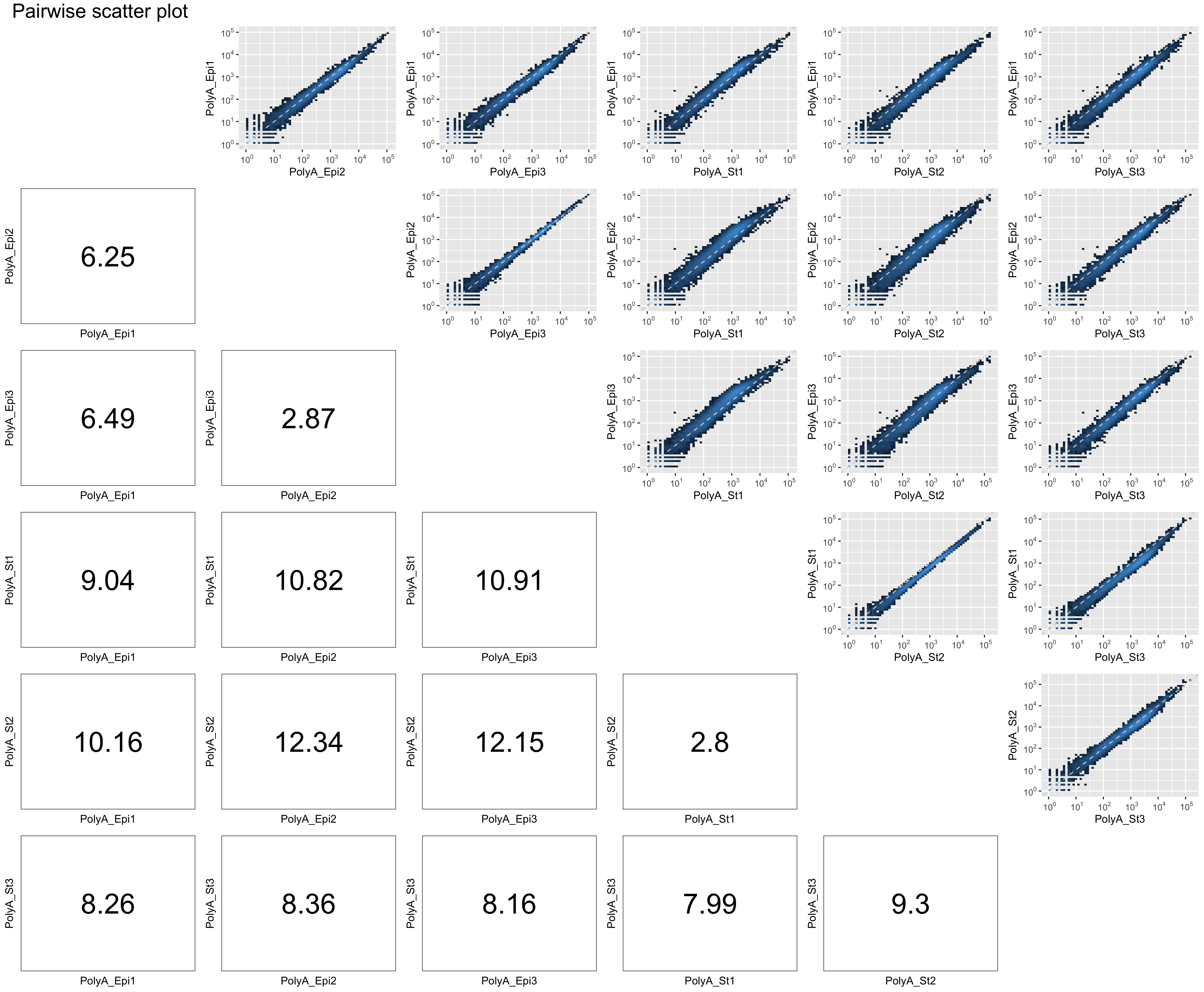

### pairwiseScatter.png

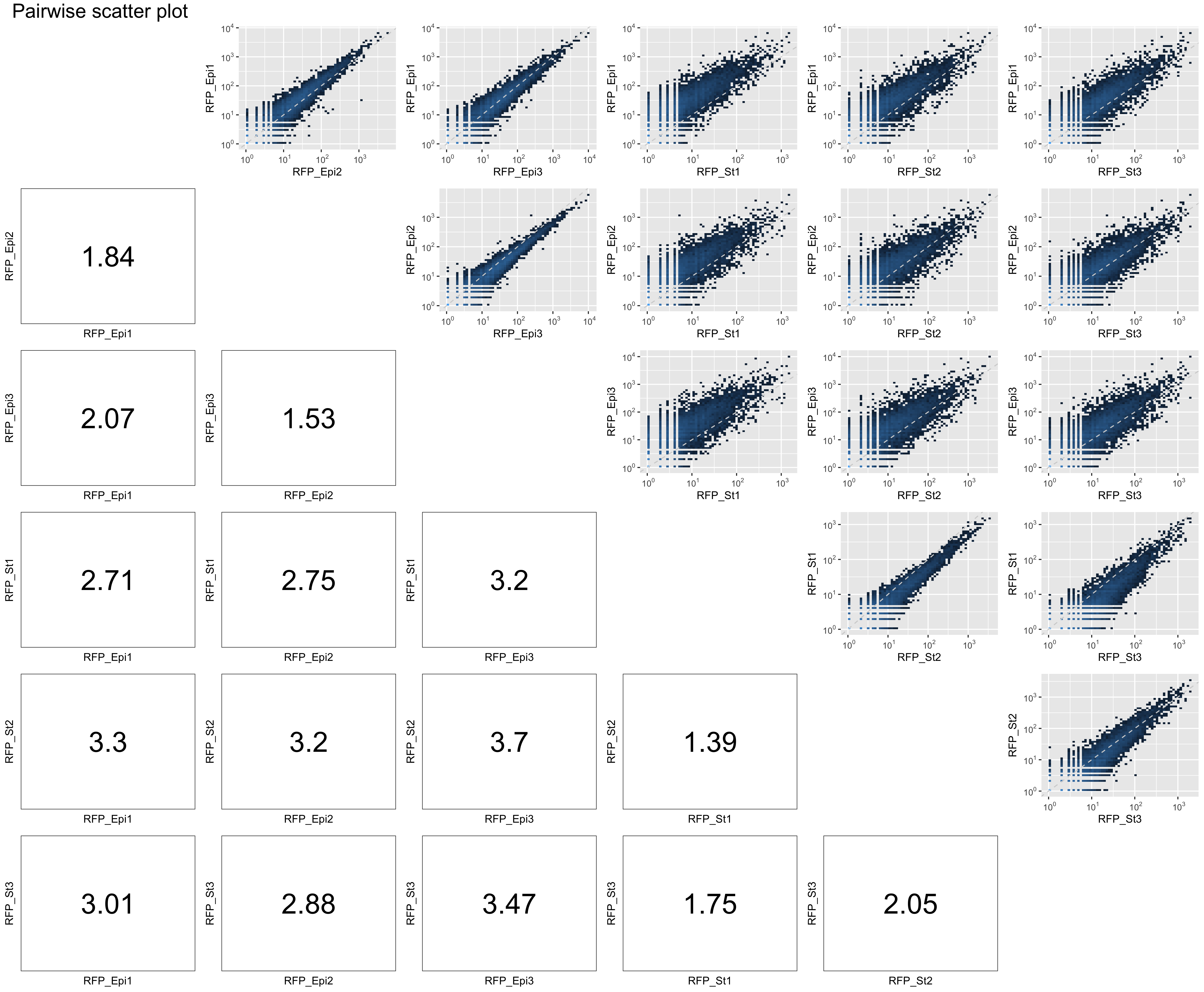

### PCA.png

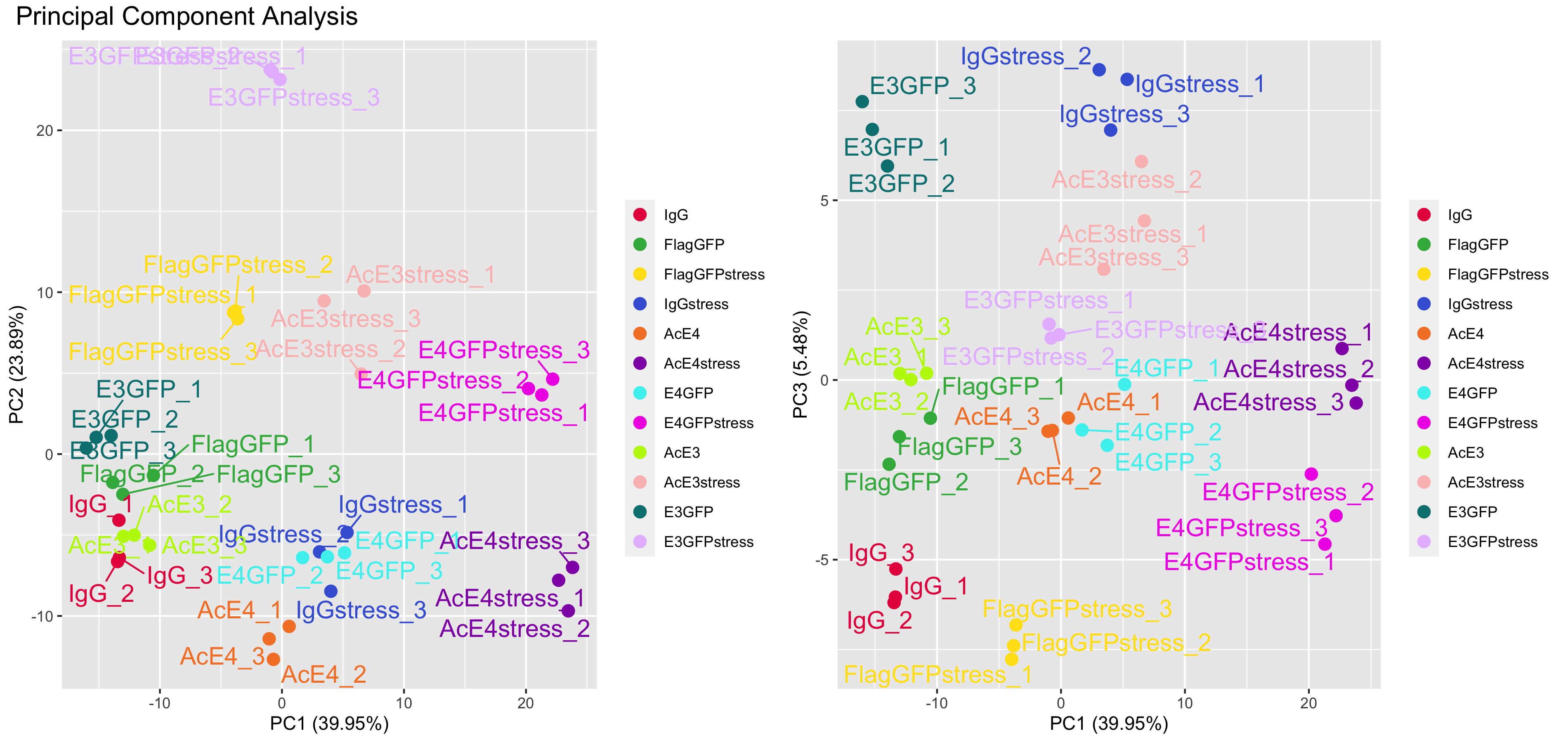

### PCA.png

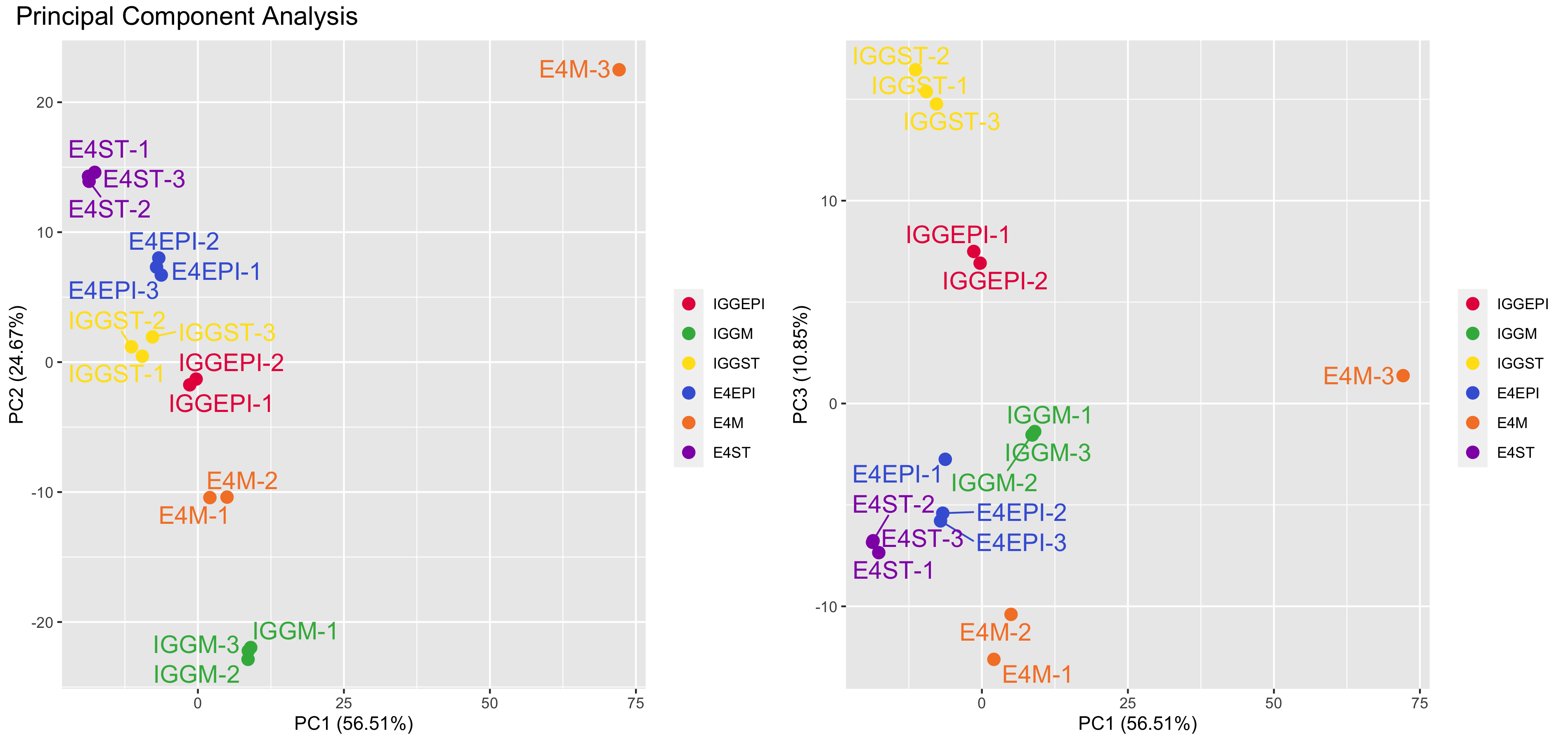

### PCA.png

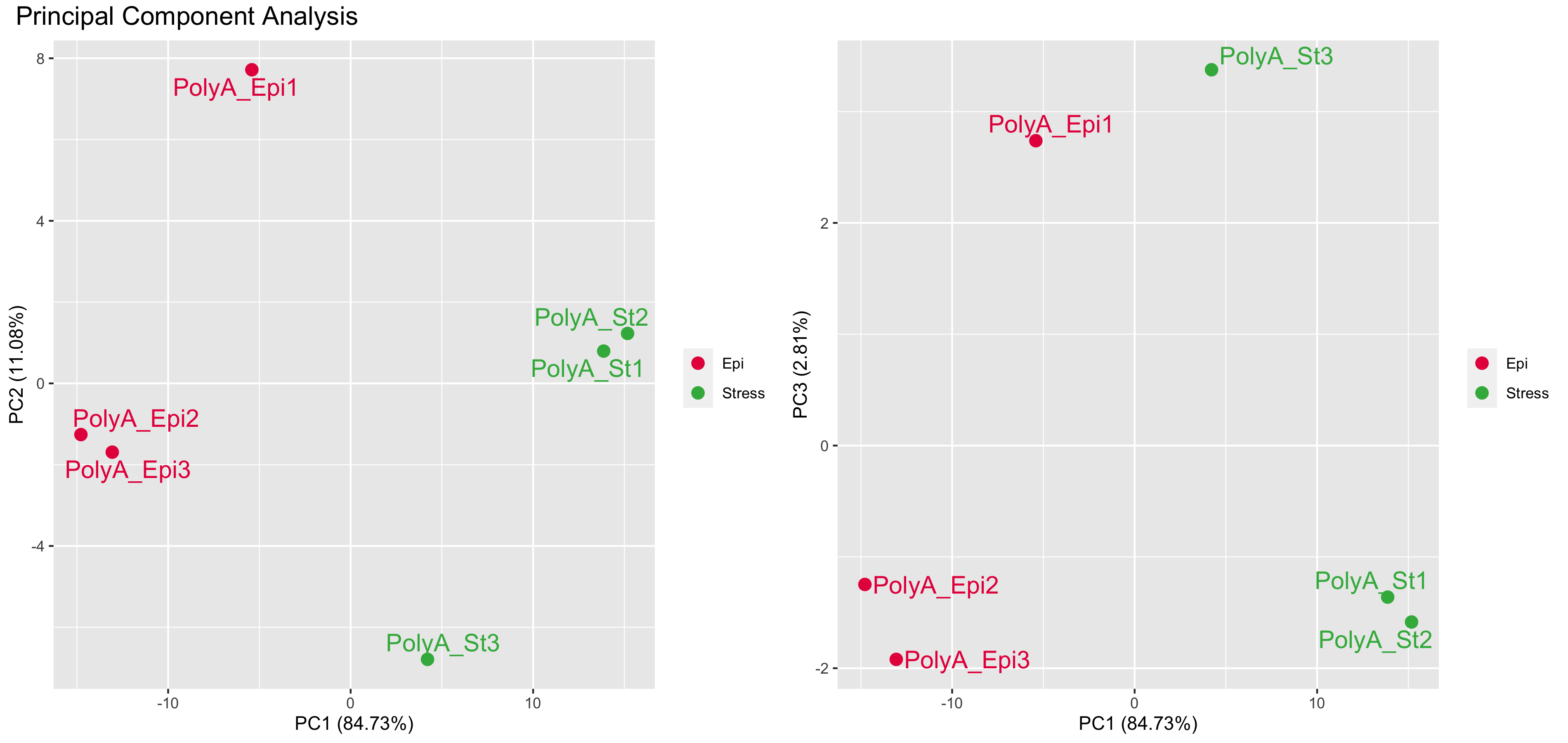

### PCA.png

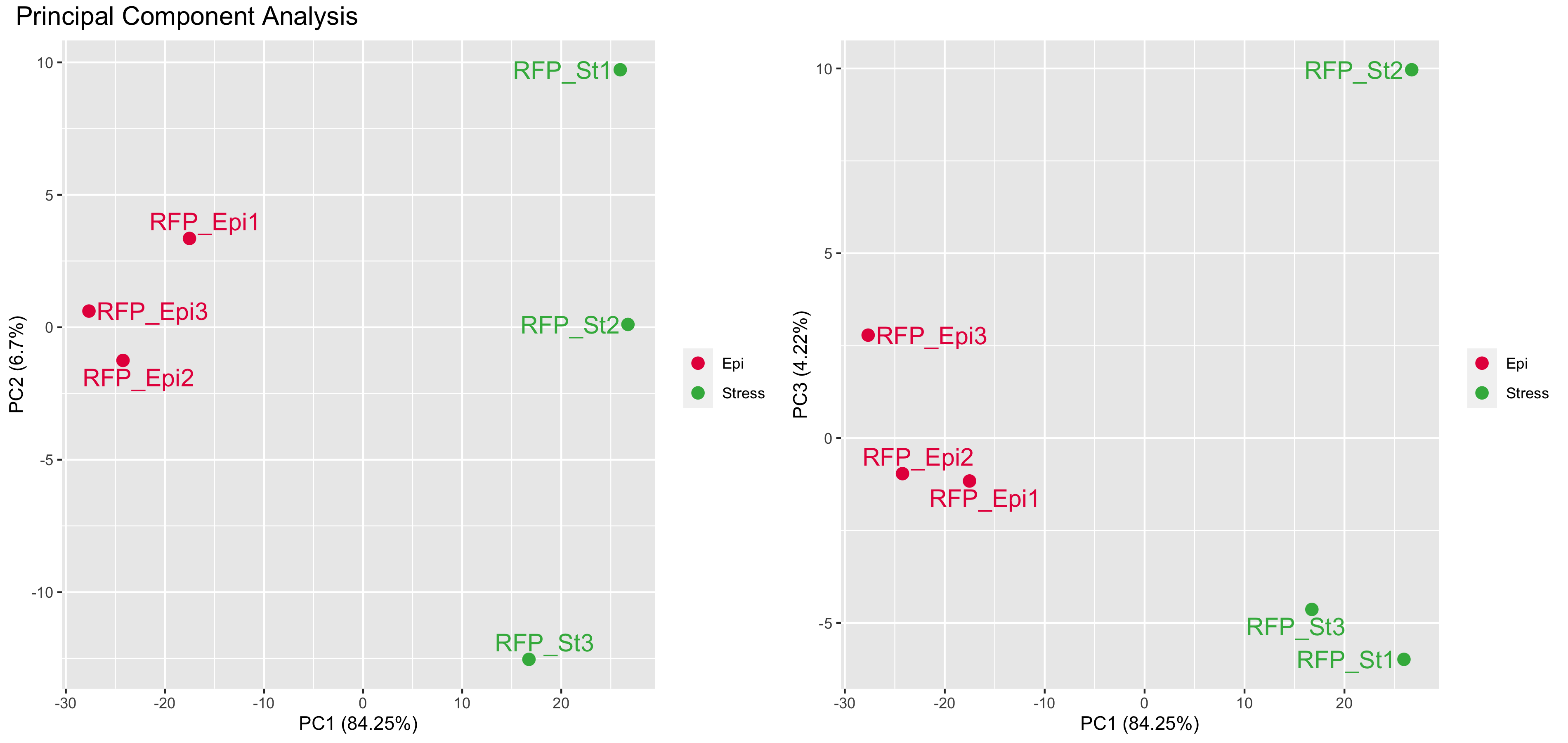

### rawpHist.png

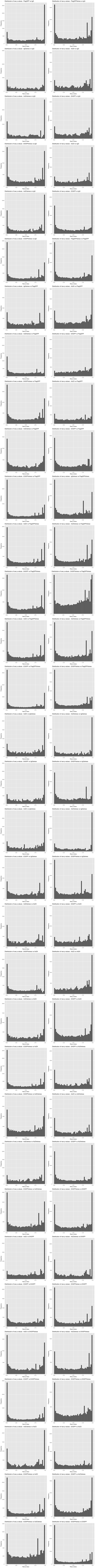

### rawpHist.png

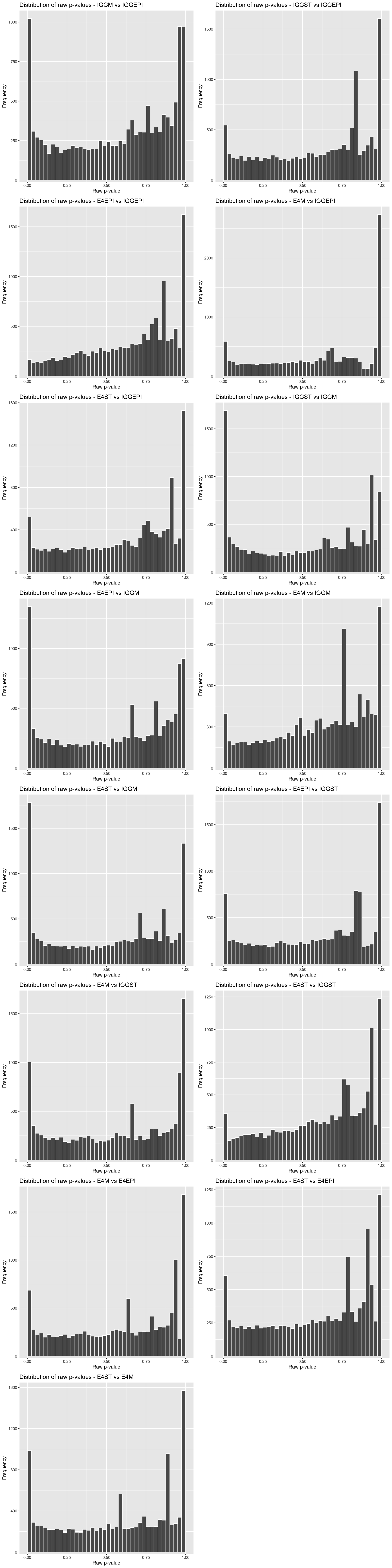

### rawpHist.png

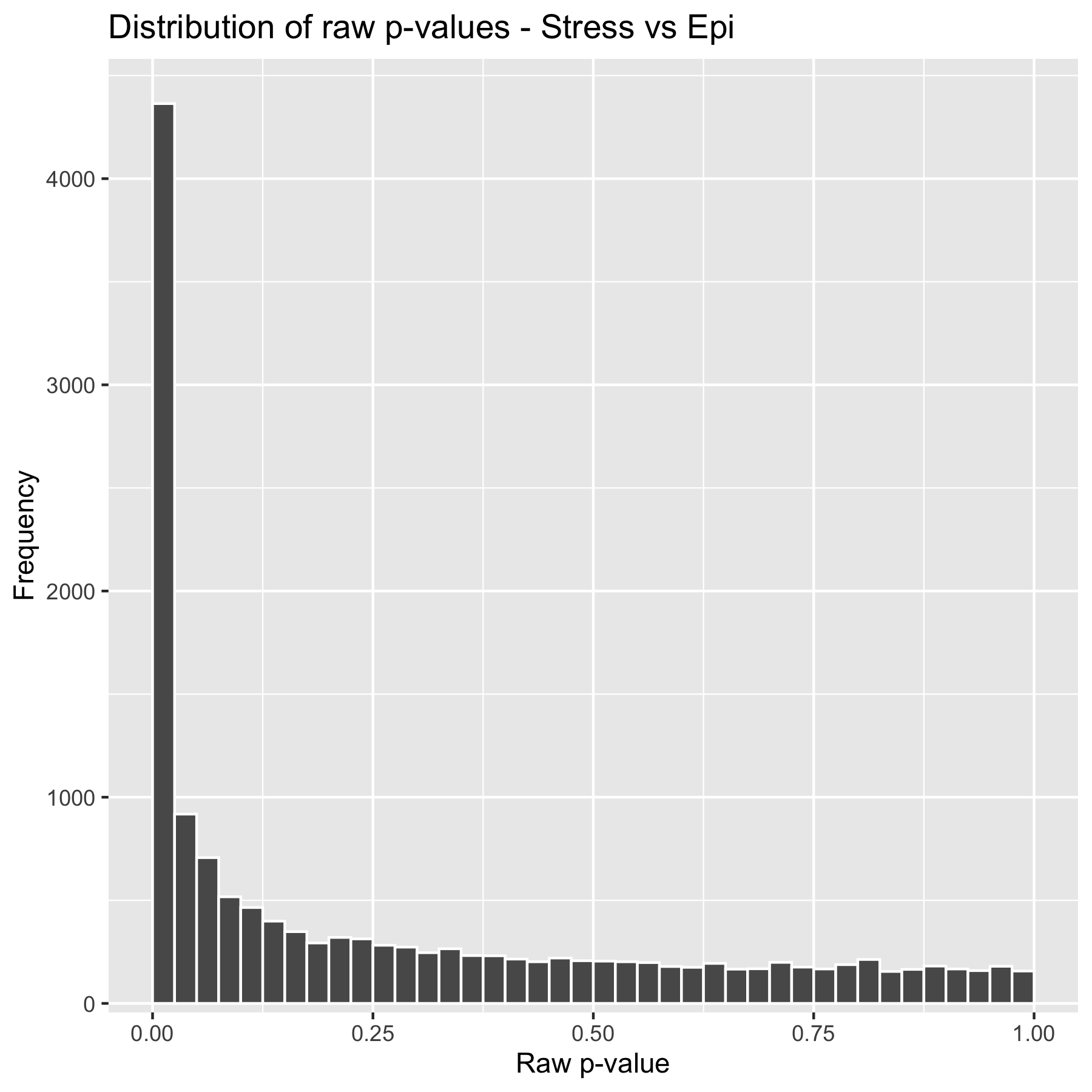

### rawpHist.png

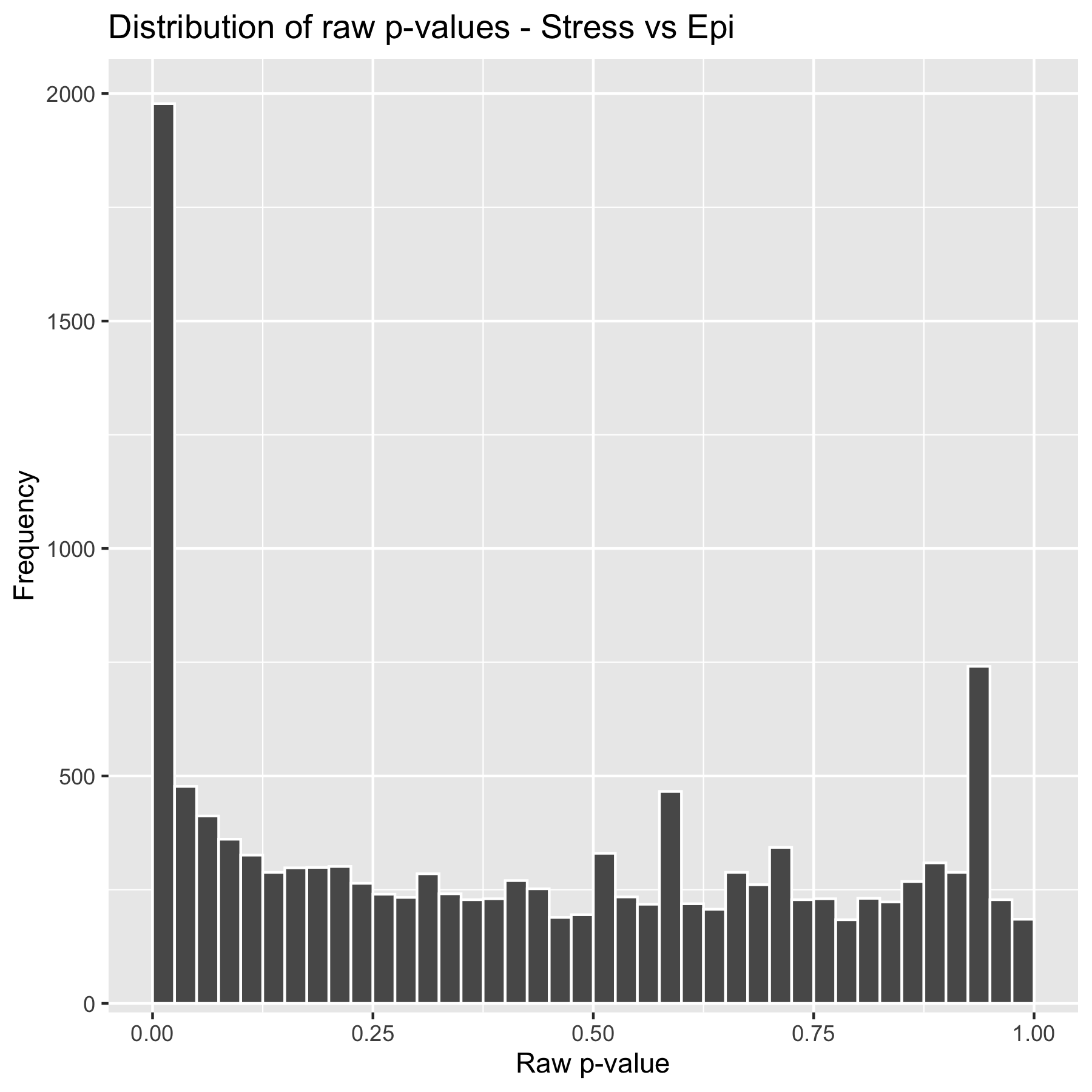

### volcanoPlot.png

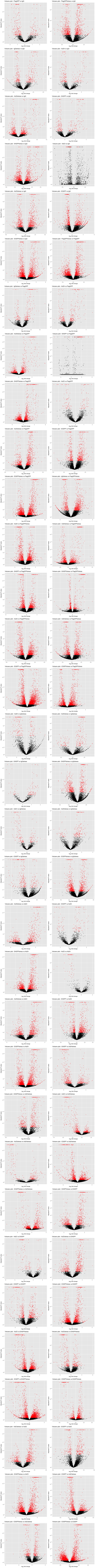

### volcanoPlot.png

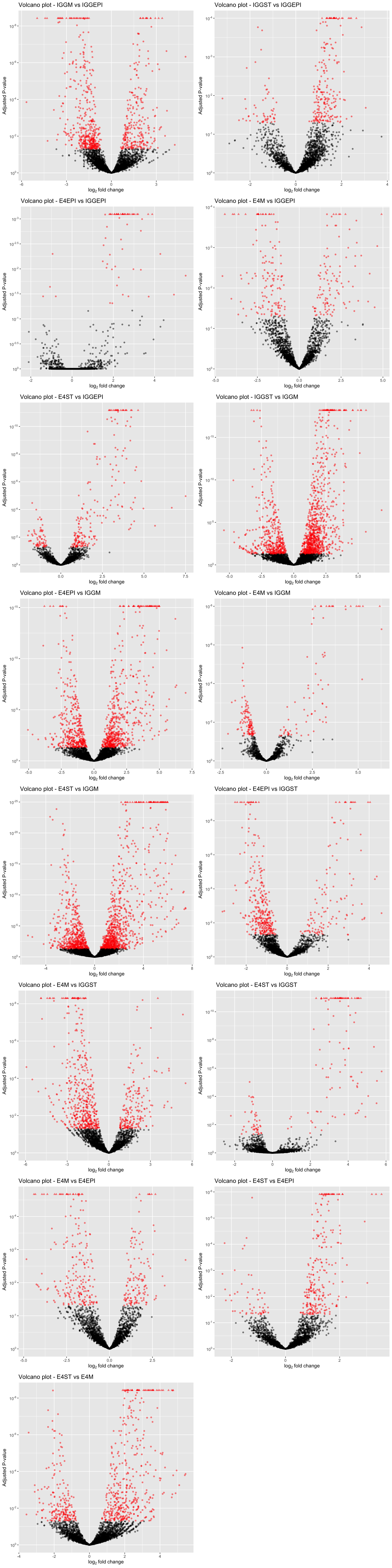

### volcanoPlot.png

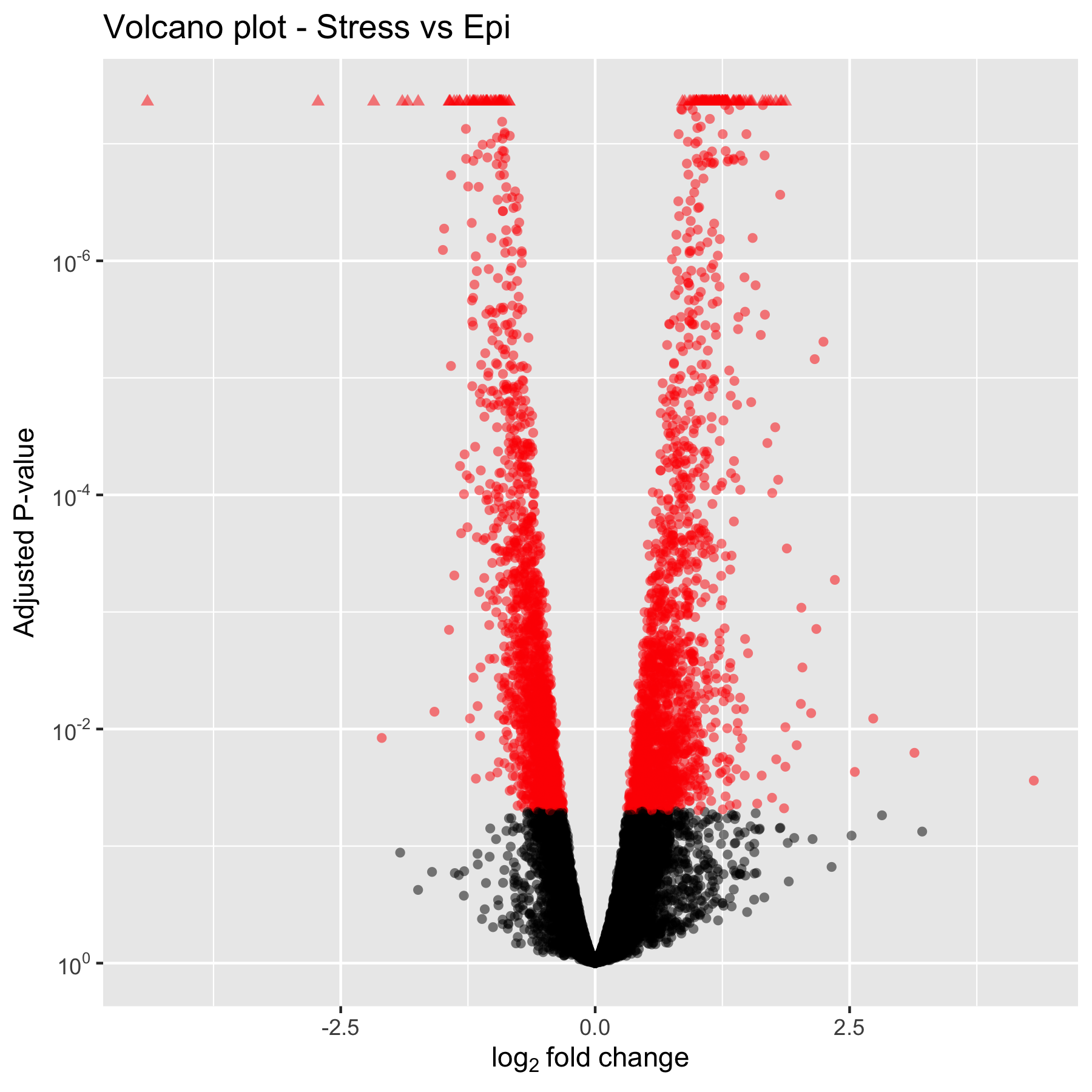

### volcanoPlot.png

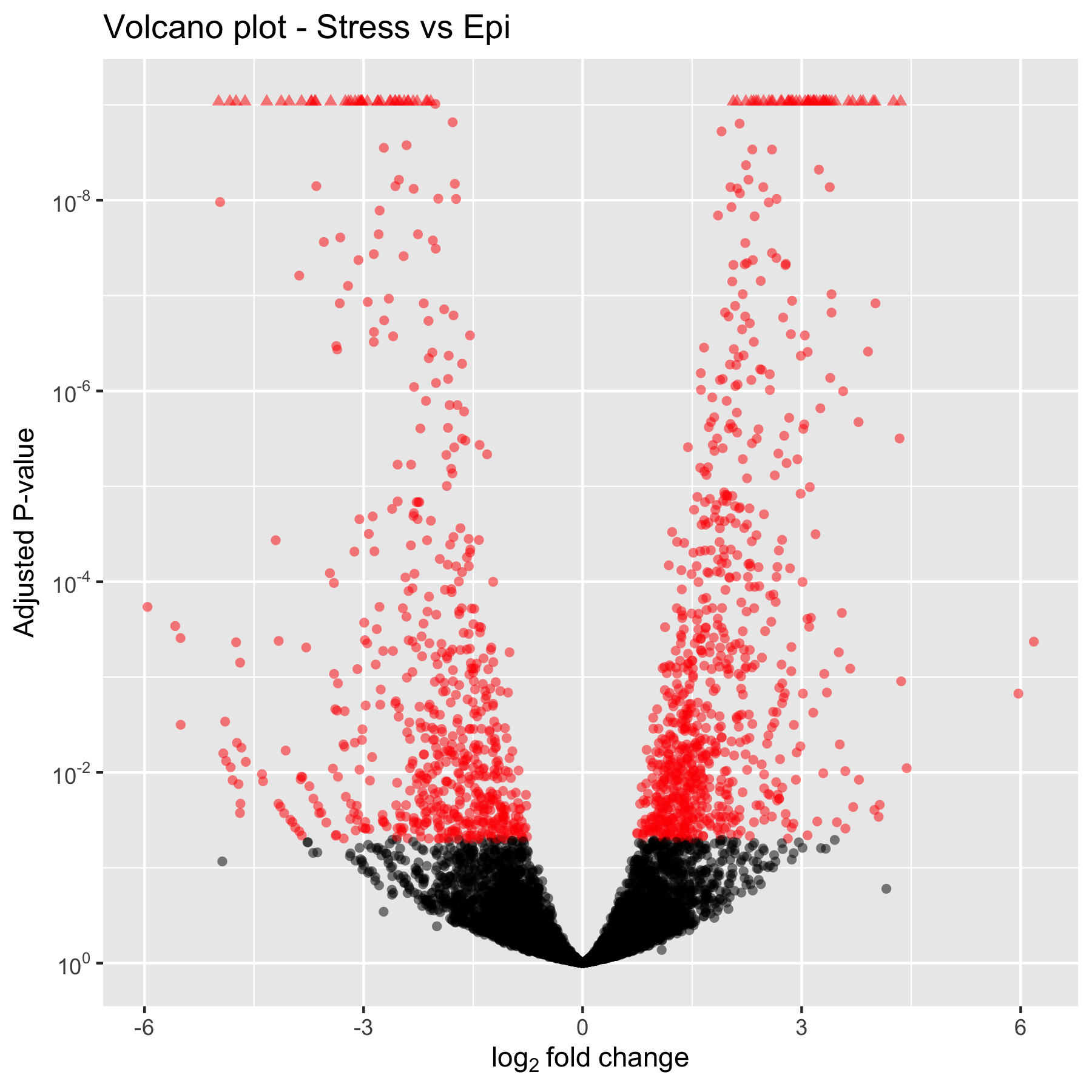
