## Supplementary material for "Two distinct *Trypanosoma* eIF4F complexes co-exist, bind different mRNAs and are regulated during nutritional stress": Figure S9

| | | 1 | 2 | 3 | 4 | 5 | 6 | 7 | 8 | 9 | 10 | 11 | 12 | 13 | 14 | 15 | | |
|---|---|---|---|---|---|---|---|---|---|---|---|---|---|---|---|---|---|---|
| **Consensus PAM2 motif** | | **x** | **x** | **Φ** | **N** | **X** | **X** | **A** | **x** | **E** | **F** | **X** | **P** | **x** | **x** | **x** | | |
| TcG1-IP1 | 1 | | M | L | N | I | N | A | K | A | F | V | P | Q | S | A | 15 | Cytoplasmic cap-modifying enzymes |
| TcG5-IP | 1 | M | S | L | N | P | N | A | M | P | W | N | M | P | E | S | 16 | |
| TcCE1 | 2 | S | A | F | N | P | D | A | P | A | F | I | P | T | F | L | 17 | |
| TcZC3H32 | 187 | R | E | L | N | P | Q | A | Q | P | F | V | P | S | P | L | 201 | RNA binding proteins |
| TcDRBD17 | 109 | S | T | L | N | P | D | A | K | E | F | Q | L | P | S | G | 123 | |
| TcRBP35 | 478 | S | T | M | N | A | R | A | R | E | F | V | P | S | S | T | 492 | |
| TcPCD6 | 61 | H | E | P | N | P | N | A | P | A | F | V | P | A | E | I | 75 | |
| Uncharacterized protein | 1 | | | M | N | P | D | A | P | A | F | I | P | A | R | D | 15 | |
