## Supplemental Files 1-4 for "Two distinct *Trypanosoma* eIF4F complexes co-exist, bind different mRNAs and are regulated during nutritional stress": 2023-06-27-PolyA_Multicounts_combined_samples_report.html

Statistical report of project 2023-06-27-PolyA\_Multicounts\_combined\_samples: pairwise comparison(s) of conditions with DESeq2


### Statistical report of project 2023-06-27-PolyA\_Multicounts\_combined\_samples: pairwise comparison(s) of conditions with DESeq2

###### MGF

#### 2023-08-25

The SARTools R package which generated this report has been developped at PF2 - Institut Pasteur by M.-A. Dillies and H. Varet. Thanks to cite H. Varet, L. Brillet-Guéguen, J.-Y. Coppee and M.-A. Dillies, *SARTools: A DESeq2- and EdgeR-Based R Pipeline for Comprehensive Differential Analysis of RNA-Seq Data*, PLoS One, 2016, doi: http://dx.doi.org/10.1371/journal.pone.0157022 when using this tool for any analysis published.

### 1 Introduction

The analyses reported in this document are part of the 2023-06-27-PolyA\_Multicounts\_combined\_samples project. The aim is to find features that are differentially expressed between Epi and Stress. The statistical analysis process includes data normalization, graphical exploration of raw and normalized data, test for differential expression for each feature between the conditions, raw p-value adjustment and export of lists of features having a significant differential expression between the conditions.

The analysis is performed using the R software [1], Bioconductor [2] packages including DESeq2 [3,4] and the SARTools package developed at PF2 - Institut Pasteur. Normalization and differential analysis are carried out according to the DESeq2 model and package. This report comes with additional tab-delimited text files that contain lists of differentially expressed features.

For more details about the DESeq2 methodology, please refer to its related publications [3,4].

### 2 Description of raw data

The count data files and associated biological conditions are listed in the following table.

Table 1: Data files and associated biological conditions.

| Sample | File | group |
| --- | --- | --- |
| PolyA\_Epi1 | Epi1.txt | Epi |
| PolyA\_Epi2 | Epi2.txt | Epi |
| PolyA\_Epi3 | Epi3.txt | Epi |
| PolyA\_St1 | St1.txt | Stress |
| PolyA\_St2 | St2.txt | Stress |
| PolyA\_St3 | St3.txt | Stress |

After loading the data we first have a look at the raw data table itself. The data table contains one row per annotated feature and one column per sequenced sample. Row names of this table are feature IDs (unique identifiers). The table contains raw count values representing the number of reads that map onto the features. For this project, there are 15319 features in the count data table.

Table 2: Partial view of the count data table.

|  | PolyA\_Epi1 | PolyA\_Epi2 | PolyA\_Epi3 | PolyA\_St1 | PolyA\_St2 | PolyA\_St3 |
| --- | --- | --- | --- | --- | --- | --- |
| C4B63\_100g14 | 288 | 317 | 254 | 174 | 250 | 266 |
| C4B63\_100g16 | 9 | 18 | 13 | 14 | 25 | 18 |
| C4B63\_100g24 | 75 | 74 | 82 | 56 | 74 | 67 |
| C4B63\_100g30 | 370 | 347 | 291 | 474 | 728 | 682 |
| C4B63\_100g31 | 14 | 6 | 8 | 11 | 19 | 12 |
| C4B63\_100g32 | 25 | 12 | 8 | 10 | 16 | 22 |

Looking at the summary of the count table provides a basic description of these raw counts (min and max values, median, etc).

Table 3: Summary of the raw counts.

|  | Min. | 1st Qu. | Median | Mean | 3rd Qu. | Max. |
| --- | --- | --- | --- | --- | --- | --- |
| PolyA\_Epi1 | 0 | 123 | 770 | 1499 | 1954 | 100716 |
| PolyA\_Epi2 | 0 | 126 | 885 | 1798 | 2337 | 116685 |
| PolyA\_Epi3 | 0 | 127 | 902 | 1739 | 2316 | 105346 |
| PolyA\_St1 | 0 | 110 | 600 | 1315 | 1428 | 117410 |
| PolyA\_St2 | 0 | 170 | 922 | 1963 | 2162 | 171031 |
| PolyA\_St3 | 0 | 162 | 1023 | 2042 | 2559 | 164449 |

Figure 1 shows the total number of mapped and counted reads for each sample. We expect total read counts to be similar within conditions, they may be different across conditions. Total counts sometimes vary widely between replicates. This may happen for several reasons, including:

- different rRNA contamination levels between samples (even between biological replicates);
- slight differences between library concentrations, since they may be difficult to measure with high precision.

Figure 1: Number of mapped reads per sample. Colors refer to the biological condition of the sample.

Figure 2 shows the percentage of features with no read count in each sample. We expect this percentage to be similar within conditions. Features with null read counts in the 6 samples are left in the data but are not taken into account for the analysis with DESeq2. Here, 708 features (4.62%) are in this situation (dashed line). Results for those features (fold-change and p-values) are set to NA in the results files.

Figure 2: Percentage of features with null read counts in each sample.

Figure 3 shows the distribution of read counts for each sample (on a log scale to improve readability). Again we expect replicates to have similar distributions. In addition, this figure shows if read counts are preferably low, medium or high. This depends on the organisms as well as the biological conditions under consideration.

Figure 3: Density distribution of read counts.

It may happen that one or a few features capture a high proportion of reads (up to 20% or more). This phenomenon should not influence the normalization process. The DESeq2 normalization has proved to be robust to this situation [Dillies, 2012]. Anyway, we expect these high count features to be the same across replicates. They are not necessarily the same across conditions. Figure 4 and table 4 illustrate the possible presence of such high count features in the data set.

Figure 4: Percentage of reads associated with the sequence having the highest count (provided in each box on the graph) for each sample.

Table 4: Percentage of reads associated with the sequences having the highest counts.

|  | C4B63\_52g313c | C4B63\_333g19 | C4B63\_54g30 | C4B63\_2g166c | C4B63\_333g15 |
| --- | --- | --- | --- | --- | --- |
| PolyA\_Epi1 | 0.44 | 0.31 | 0.30 | 0.24 | 0.27 |
| PolyA\_Epi2 | 0.42 | 0.27 | 0.30 | 0.28 | 0.21 |
| PolyA\_Epi3 | 0.40 | 0.23 | 0.27 | 0.23 | 0.20 |
| PolyA\_St1 | 0.58 | 0.48 | 0.45 | 0.24 | 0.47 |
| PolyA\_St2 | 0.57 | 0.51 | 0.38 | 0.21 | 0.49 |
| PolyA\_St3 | 0.53 | 0.28 | 0.34 | 0.22 | 0.26 |

Table 5: Normalization factors.

|  |  |  |  |  |  |  |
| --- | --- | --- | --- | --- | --- | --- |
| Size factor | 0.92 | 1.05 | 1.04 | 0.73 | 1.12 | 1.23 |

The histograms (figure 8) can help to validate the choice of the normalization parameter (“median” or “shorth”). Under the hypothesis that most features are not differentially expressed, each size factor represented by a red line is expected to be close to the mode of the distribution of the counts divided by their geometric means across samples.

Table 6 reports the thresholds used for each comparison and the number of features discarded by the independent filtering. Adjusted p-values of discarded features are then set to NA.

Table 6: Number of features discarded by the independent filtering for each comparison.

| Test vs Ref | BaseMean Threshold | # discarded |
| --- | --- | --- |
| Stress vs Epi | 3.28 | 991 |

#### 5.6 Final results

A p-value adjustment is performed to take into account multiple testing and control the false positive rate to a chosen level \(\alpha\). For this analysis, a BH p-value adjustment was performed [8 and BY2001] and the level of controlled false positive rate was set to 0.05.

Table 7: Number of up-, down- and total number of differentially expressed features for each comparison.

| Test vs Ref | # down | # up | # total |
| --- | --- | --- | --- |
| Stress vs Epi | 1763 | 1797 | 3560 |

Figure 13 represents the MA-plot of the data for the comparisons done, where differentially expressed features are highlighted in red. A MA-plot represents the log ratio of differential expression as a function of the mean intensity for each feature. Triangles correspond to features having a too low/high \(\log\_2(\text{FC})\) to be displayed on the plot.

Parameter values used for this analysis are:

- workDir: /Users/ferrarini/Desktop/Artigos/Ongoing/2023-Bernardo/SARTools/multicounts/
- projectName: 2023-06-27-PolyA\_Multicounts\_combined\_samples
- author: MGF
- targetFile: colData\_PolyA.txt
- rawDir: countData
- featuresToRemove: alignment\_not\_unique, ambiguous, no\_feature, not\_aligned, too\_low\_aQual
- varInt: group
- condRef: Epi
- batch: NULL
- fitType: parametric
- cooksCutoff: TRUE
- independentFiltering: TRUE
- alpha: 0.05
- pAdjustMethod: BH
- typeTrans: VST
- locfunc: median
- colors: #e6194b, #3cb44b, #ffe119, #4363d8, #f58231, #911eb4, #46f0f0, #f032e6, #bcf60c, #fabebe, #008080, #e6beff, #9a6324, #fffac8, #800000, #aaffc3, #808000, #ffd8b1, #000075, #808080, #ffffff, #000000

### Bibliography

1.

R Core Team. *R: A language and environment for statistical computing*. Vienna, Austria : R Foundation for Statistical Computing, 2017 :

2.

Gentleman RC, Carey VJ, Bates DM, *et al.* Bioconductor: Open software development for computational biology and bioinformatics. *Genome Biology* 2004 ; 5 : R80.

3.

Anders S, Huber W. Differential expression analysis for sequence count data. *Genome Biology* 2010 ; 11 : R106.

4.

Love MI, Huber W, Anders S. Moderated estimation of fold change and dispersion for RNA-seq data with DESeq2. *Genome Biology* 2014 ; 15 : 550.

5.

Schulze SK, Kanwar R, Gölzenleuchter M, *et al.* SERE: Single-parameter quality control and sample comparison for RNA-seq. *BMC Genomics* 2012 ; 13 : 524.

6.

Cook RD. Detection of influential observation in linear regression. *Technometrics* 1977 ; 19 : 15–18.

7.

Cox DR, Reid N. Parameter orthogonality and approximate conditional inference. *Journal of the Royal Statistical Society. Series B (Methodological)* 1987 ; 49 : 1–39.

8.

Benjamini Y, Hochberg Y. Controlling the false discovery rate: A practical and powerful approach to multiple testing. *Journal of the Royal Statistical Society. Series B (Methodological)* 1995 ; 57 : 289–300.
